# Structural reconstruction of individual filaments in Aβ42 fibril populations assembled *in vitro* reveal rare species that resemble *ex vivo* amyloid polymorphs from human brains

**DOI:** 10.1101/2023.07.14.549001

**Authors:** Liam D. Aubrey, Liisa Lutter, Kate Fennell, Tracey J. Purton, Natasha Ward, Louise C. Serpell, Wei-Feng Xue

## Abstract

Structural polymorphism has been demonstrated for both *in vitro* and *ex vivo* amyloid fibrils associated with disease. The manner in which different filament structures are assembled from common building blocks remains unclear but the assembly environment is likely to be a key determinant. To address this, three-dimensional reconstruction of individual filament structures was conducted from atomic force microscopy images to map the structural polymorphism landscape of Aβ_42_ amyloid fibril populations formed *in vitro* under most frequently used buffer conditions. The data show sensitivity of Aβ_42_ fibril polymorphism to the assembly environment in both the magnitude of heterogeneity and the types of filament species formed. However, some conserved fibril polymorphs were observed across the experimental conditions. Excitingly, by matching individual filament structures to cryo-electron microscopy derived structural data, rare species in these heterogeneous population clouds that show remarkable similarity to Aβ_42_ amyloid polymorphs purified from human patient brains were discovered. These results link *in vitro* experimental approaches with structures formed *in vivo*, and highlight the polymorph distribution, and the type and magnitude of structural variations within these heterogeneous molecular distributions as important factors in amyloid biology.

## Introduction

Protein misfolding and the formation of amyloid fibril structures is associated with pathology in numerous diseases ^1, 2^. This includes, but is not limited to, neurodegenerative diseases such as Alzheimer’s disease (AD), in which extracellular deposits known as amyloid plaques are found in brains of patients. The plaques are composed of amyloid fibrils made from cleavage products of the amyloid precursor protein (APP) ^3^. Amongst these fragments, the 42 residue amyloid β 1-42 peptide (Aβ_42_) is considered to be an important aggregation prone fragment that has been shown to produce species which are toxic to neuronal cells ^4, 5^ and fibrils that are part of insoluble deposits in human brains ^6^. Aβ_42_ amyloid fibrils, like all amyloid fibrils, are defined by their core cross-β molecular architecture composed of β strands that stack perpendicular to the long fibril axis to form protofilaments up to several microns in length ^7, 8^. Multiple protofilaments can laterally associate to form twisted fibrils with a hydrophobic core ^9–12^. Various experiments have been performed to determine the specific structural properties of Aβ fibrils. Atomic detailed structures have been generated from solid state NMR (ssNMR) (e.g. ^13–17^) and cryo-transmission electron microscopy (cryo-TEM) (e.g. ^6, 18–26^). Interestingly, different structural polymorphs have been observed for fibrils formed from both amyloid β 1-40 peptide (Aβ_40_) ^25, 26^ and Aβ_42_ ^6, 22^. In one of the studies of *ex-vivo* Aβ_42_ fibrils purified from human brains ^6^, multiple left-hand twisted polymorphs were observed in the same samples originating from the same patients (**Figure 1a and 1b**). Contrastingly, in patient brain derived Aβ_40_ fibrils from a meningeal sample ^25^, the most populous fibril structure was a right-hand twisted fibril with two laterally associated protofilaments. In a separate studies with *ex-vivo* seeded samples, a Aβ_40_ and an Aβ_42_ structure displayed a left-handed twist while another Aβ_42_ structure displayed a right-handed twist, all three with two protofilaments laterally associated ^22, 26^. These studies indicate the potential for different structural variation arising in different local environments, such as different brain regions. However, the range of structural polymorphs Aβ_42_ amyloid fibrils can adopt has not been mapped.

**Figure 1.**
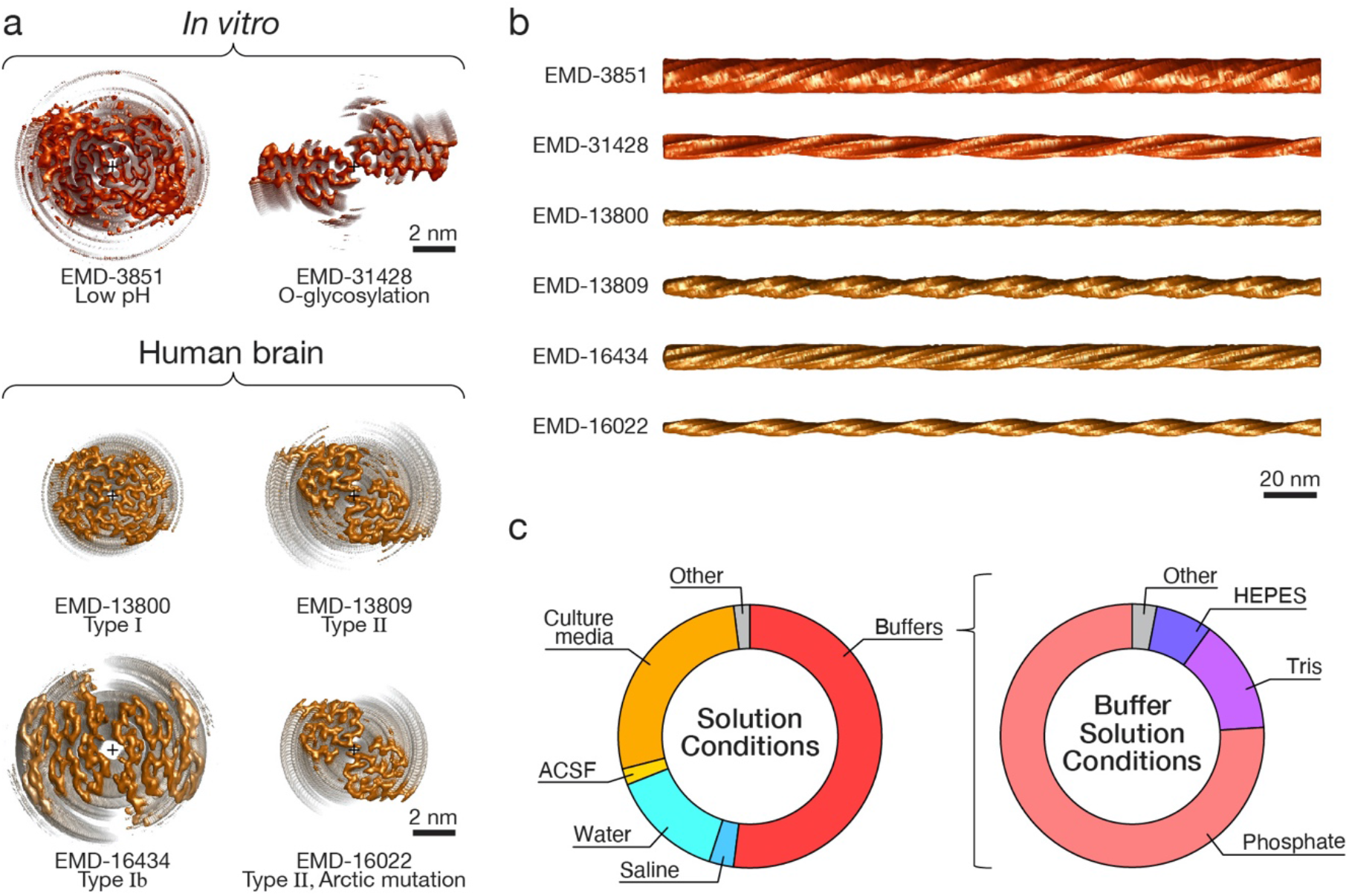
Aβ_42_ amyloid core structures of human brain derived and *in vitro* assembled fibrils demonstrate diverse structures and suggest assembly condition-dependent structural polymorphism. (a) Examples of recent cryo-TEM derived maps of Aβ_42_ amyloid filament core deposited in the EMDB with reported resolution of 4Å or better. The filament cross-sections are shown for two *in vitro* assembled fibril polymorphs (EMD-3851 ^21^ and EMD-31428 ^69^ coloured in bronze) and the Type I, Ib and II fibril polymorphs seen in human patient brain samples (EMD-13800, EMD-16434 and EMD-13809/EMD-16022 ^6, 24^, respectively, coloured in gold). All cross-sections are shown with identical scale, with scale bars indicating 2 nm. (b) The surface envelope of same Aβ_42_ fibril polymorphs shown in (a). The surface envelope structures are generated using axis aligned and extended cryo-TEM derived maps for comparing the long-range and helical features of the fibril polymorphs ^35^. The filaments are shown with identical scale, with scale bars indicating 20 nm. (c) Buffers used in highly cited publications between 2005 and 2020. The most highly cited publications involving Aβ_42_ from each of these years were tabulated and the buffer conditions in which monomer was incubated in which substantial polymerisation could occur were recorded. Pie charts to the left display the breakdown of various different types of solution conditions used and to the right the specific buffer salts in buffered conditions used.

In order to quantify the structural variation of fibril populations formed from Aβ_42_ and to map the extent of possible fibril structures that can be formed, *individual* fibrils in a sample population must be resolved at sufficiently high-detail to distinguish the structural properties of each individual fibril. Atomic force microscopy (AFM) imaging has previously been used to demonstrate that the distribution of morphometric features can be obtained by analysis of each individual fibril in a sample (e.g. ^27–33^). Recently, we have demonstrated that 3D models of individual fibril structures reconstructed from AFM image data can be used to map the polymorph distribution and to quantify the structural variations within a population ^29, 34^. Here, we obtained detailed images of fibrils formed from recombinant Aβ_42_ grown in solution conditions with three most prevalently used buffer salts sodium phosphate, HEPES and Tris at neutral pH *in vitro*, and mapped the polymorphic assembly landscape of Aβ_42_ by applying our individual fibril structural analysis approach. We show that across all conditions tested, certain classes of fibril polymorphs are conserved. However, the overall diversity of fibril structures *is* sensitive to small changes in the assembly conditions in both the magnitude of the population heterogeneity and the types of filament species formed. Excitingly, we identified rare fibril species in these heterogeneous populations that closely resemble Aβ_42_ amyloid purified from human brains by matching individual filament structures to cryo-EM derived structural data ^35^. These results suggest that differences in local *in vivo* environments may lead to different sets of fibril structures in different heterogenous fibril populations. Therefore, understanding structures of Aβ_42_ fibrils formed under different conditions could highlight how different Aβ fibril structures may occur and persist in clinical populations or in different cellular and tissue environments, and these conclusions thrust the biological roles of structural heterogeneity of amyloid populations into focus.

## Results

### Characterisation of purified recombinant monomeric Aβ_42_

To ensure assembly of high-quality amyloid samples for structural studies of individual fibrils, high-quality monomeric Aβ_42_ samples was first produced. Production of recombinant monomeric Aβ_42_ was optimised from previous methods ^36, 37^ to include multi-step size exclusion chromatography (SEC), to ensure high reproducibility of assembly. The final size exclusion chromatography step was performed in multiple steps until the resulting chromatogram contained a single main peak (**Supplementary Figure S1a**) that was collected and used immediately for assembly. SDS-PAGE of the eluted monomer (**Supplementary Figure S1b**) further confirmed the purity of the monomer samples. The formation of fibrils from the monomer solution was monitored using Thioflavin T fluorescence (**Supplementary Figure S1c**), demonstrating the expected concentration dependent sigmoidal kinetics profile for amyloid assembly. The expected formation of β-rich structures during assembly was also confirmed by CD (**Supplementary Figure S1d**). To verify that the samples made using the recombinant Aβ_42_ protein show the expected toxicity properties, primary hippocampal neurons were exposed to 10µM monomer equivalent concentration of oligomeric Aβ_42_ formed during assembly under previously reported conditions ^38^ and cell viability was assessed using a Live/Dead fluorescence assay (**Supplementary Figure S1e**). As expected, cell death was significantly more prevalent in cultures treated with the Aβ_42_ sample (p < 0.001), confirming its expected neurotoxicity potential ^38^. Taken together, these data show that we have generated monomeric protein samples that demonstrate all the biophysical and biochemical properties expected of assembly competent Aβ_42_.

### Selection of assembly conditions for polymorphic mapping of Aβ_42_

To investigate how solution conditions affect the extent of the polymorphism for Aβ_42_ amyloid assembly, we first identified commonly used solution conditions through an analysis of published experimental Aβ_42_ studies in the literature. In Alzheimer’s patients, Aβ amyloid fibrils are found in extracellular plaques in the brain. In the early stages of Alzheimer’s, the hippocampal region of the brain is especially vulnerable although as the disease progresses other areas of the brain can be impacted resulting in different symptoms. Different brain regions are likely to have different localised conditions resulting in different Aβ aggregation propensity, behaviour and potentially disease phenotype. This behaviour is also observed in other amyloid fibrils such as tau in which *ex vivo* structures from patients with different tauopathies have varying structures as determined by cryo-TEM ^39^. For *in vitro* studies, the conditions used to utilise or to resolve Aβ_42_ fibril structures vary considerably between reports, and an array of different Aβ_42_ fibril preparation methods are also employed in the field. In particular, different solution conditions are used to generate Aβ_42_ amyloid samples for molecular, structural, biophysical and cellular studies. We selected the top 20 most cited publications from each of the years (according to Google Scholar and Web of Science) running up to May 2020. Selected publications (presented in a tabulated form in the **Supplementary Table S1**) were included on the basis that they contained at least one Aβ_42_ experiment in which some form of amyloid assembly had occurred. A quantitative analysis of the buffer conditions used in this literature dataset is shown in **Figure 1c**. As seen in **Figure 1c** and in **Supplementary Table S1**, phosphate buffers were the most used (primarily sodium phosphate or phosphate buffered saline which is a mixture of sodium and potassium phosphates weighted heavily towards sodium) although Tris and HEPES buffers were also frequently used. Based on this analysis, to test the effect of solution conditions on the extent of Aβ_42_ fibril polymorphism and the heterogeneity of Aβ_42_ fibril populations, the three most prevalently used buffer salts: sodium phosphate, HEPES and Tris, were selected. Subsequently, Aβ_42_ fibril formation in the presence of these buffer salts was carried out at 37°C and pH 7.4. In order to compare variation in the extent of the structural polymorphism present in the amyloid fibril population that arises through altering the buffer salts used for amyloid assembly, fibril samples were prepared in an identical manner, except that the final monomer purification steps were performed in the appropriate buffer at pH 8.0 to avoid aggregation on the column ^36^. Subsequently, an appropriate solution was added to the monomer to ensure that 20 mM of the buffer salt and a pH of 7.4 were achieved in the final assembly reaction solutions and sodium phosphate buffer at pH 8 were also included in the set of conditions used.

### Assembly of Aβ_42_ result in amyloid fibril samples with high degree of structural heterogeneity

Previous work to determine the structure of amyloid fibrils formed from Aβ has demonstrated their morphology to be highly variable, even for amyloid fibrils assembled under identical conditions in the same sample or when extracted from Alzheimer’s patients ^6, 39–41^. Therefore, to understand how Aβ_42_ amyloid structures link with disease aetiology, the assembly landscape relating to the fibril polymorphs that can form must first be mapped and understood.

In order to map the polymorphic assembly landscape of Aβ_42_, we next employed peak-force tapping mode AFM imaging (ScanAsyst imaging mode, Bruker), which allows for topographical imaging with a high level of control over the imaging force applied. Low magnification AFM images shows that the Aβ_42_ fibrils were dispersed fairly randomly with some noticeable lateral association between filaments as well as some bundling of fibrils (**Figure 2a**). Qualitatively, the overall suprastructural appearance of the fibrils show subtle differences in HEPES and Tris compared to the samples in phosphate buffers. In phosphate buffer, particularly at pH 8.0, fibrils appeared to promote more tightly packed bundles of fibrils than HEPES pH 7.4. Upon magnification, the variety in fibril structures formed under different experimental conditions is striking (**Figure 2b**). In addition, it was not uncommon to observe fibrils which progressed from one type of polymorph to another across the length of a single fibril often with a noticeable change in average height and twist pattern. This intra-fibril variation strongly supports the view that variation in protofilament assembly and organisation is an important molecular mechanism of polymorphism ^42^. In summary, considerable structural variations and heterogeneity are observed in all samples across all conditions tested. These data demonstrate that Aβ_42_ fibril assembly have a high propensity for polymorphism and produce highly heterogeneous fibril populations.

**Figure 2.**
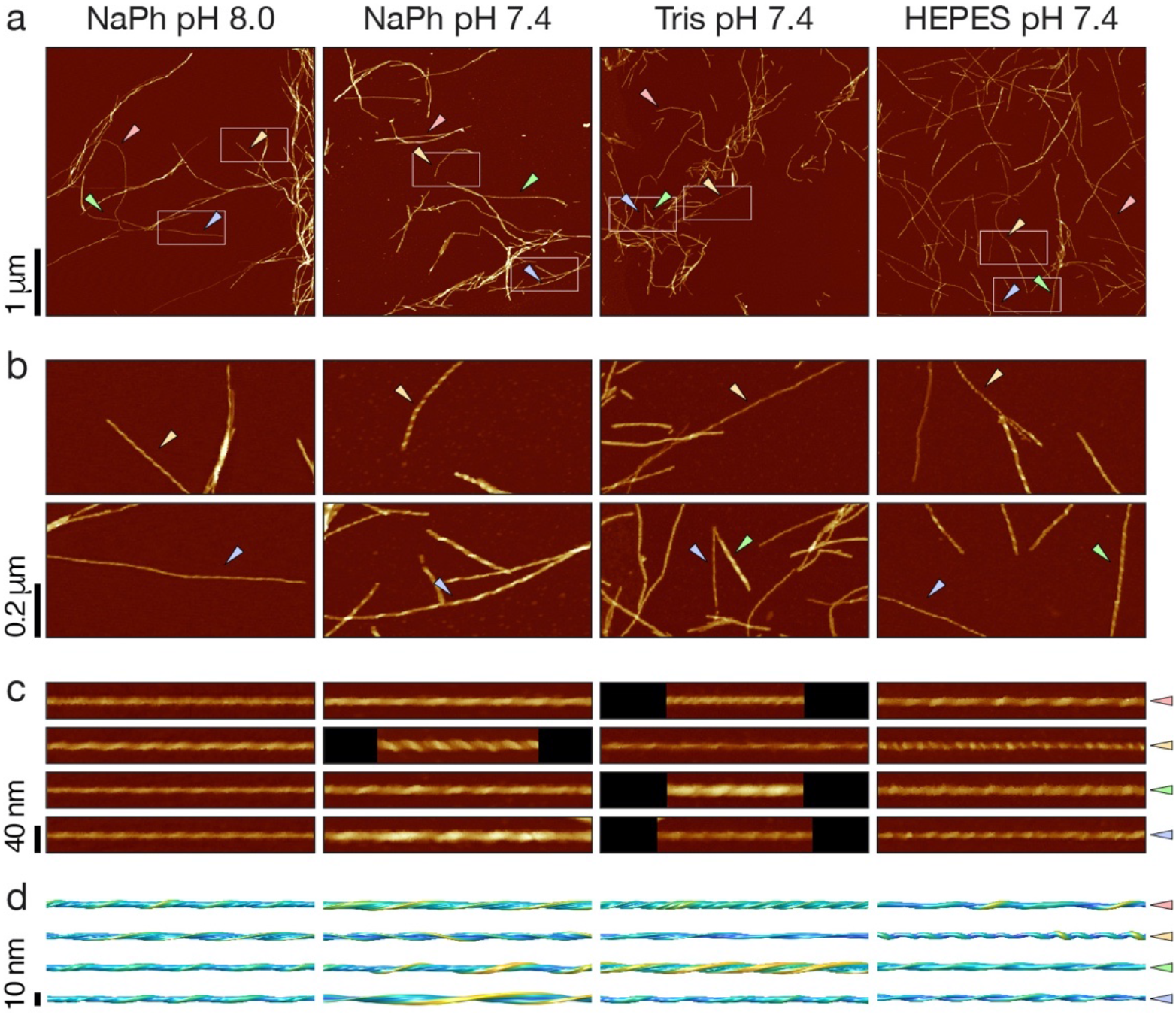
Topographical AFM height images of Aβ_42_ fibrils reveal heterogeneous and polymorphic fibril populations formed under different experimental solution conditions. AFM height images of Aβ_42_ fibrils formed in 20mM sodium phosphate (NaPh) pH 8.0, NaPh pH 7.4, Tris pH 7.4 and HEPES pH 7.4 respectively. (a) AFM height images of 2 x 2 µm areas. Qualitatively, all four types of sample appear to generate a variety of different structural polymorphs in heterogeneously mixed populations within each sample. (b) 4x magnified height images representing 1 x 0.5 µm areas shown in the white rectangles in (a) The different structural polymorphs are clearly visible at this magnification, including different twist patterns and height profiles. (c) Examples of individual fibrils traced from the images in (a) The individual fibrils are indicated by coloured triangles in (a) and (b) and up to a 400 nm digitally straightened segments are shown. (d) 200 nm segments of 3D surface envelopes reconstructed from the same fibril images shown in (c), demonstrating the diversity of structures observed in the samples. In all AFM images, the colour scale represents a height range from −5 to 10 nm.

### Quantification of structural variation in Aβ_42_ fibril populations reveal high degree of structural polymorphism that is sensitive to assembly conditions

To quantitatively assess and compare the structural variation in the fibril populations, we next analysed the three-dimensional structures of individual filaments reconstructed from the topographical AFM images using the contact point reconstruction (CPR-AFM) method described previously ^34, 43^. Individual fibrils (those which are clearly distinguishable from a bundle of fibrils) with no overlap with other fibrils, and with at least 3 repeating cross-overs and 150 nm in uninterrupted length, were selected. One hundred such fibrils from each of the four assembly conditions were traced and their individual 3D surface envelopes were reconstructed for subsequent analysis. Digitally straightened fibril images, reconstructed 3D surface envelope structural models and cross-sectional surface contact point density maps, as well as the morphometric parameters for each of the 400 fibrils resolved by AFM can be found in the **Supplementary Figures S2, S3, S4**, and **Supplementary Table 2**, respectively. The average fibril height distributions of the fibril populations formed in each of the assembly conditions (**Figure 3a**) suggest differential heterogeneity of the populations despite the similarity in the assembly conditions used. The distinctive spread of fibril structures formed in each of the different conditions used is unequivocally demonstrated by the 2-dimensional (2D) contour maps of average fibril heights against the directional periodic frequency (*dpf* = - 1/helical pitch for left-hand twisted filaments or +1/helical pitch for right-hand twisted filaments ^29^), which allows visualisation of the distinct polymorph distributions in each of the conditions (**Figure 3b**). Height and *dpf* contour map analysis, in particular, demonstrated how the fibril populations varied between the different samples. Fibrils formed in phosphate buffer, particularly at pH 8, tended to form wider filaments with a lower frequency of twist, likely promoted by a higher frequency of inter-protofilament association (**Figure 3b**). In contrast, Tris buffered reactions at pH 7.4 produced a greater range of twist patterns than the other conditions tested, which may reflect modulation of inter-protofilament interactions. Lesser variation in average height was also observed in Tris buffered reactions compared with phosphate buffered assembly. Assembly in HEPES buffer, on the other hand, produced the overall least variation in fibril width and twist compared to the other conditions tested. Both left and right handedness of twist is observed in the data, both of which have been observed in the four different Aβ samples. However, right-hand twisted fibrils remain rarely observed across all four assembly conditions, accounting for only 10.3% (41) of the 400 fibrils analysed. Overall, these polymorph distribution maps suggest that both the magnitude of heterogeneity and the types of filament species formed are sensitive to assembly conditions. This is especially remarkable considering the apparent similarity of the assembly conditions used.

**Figure 3.**
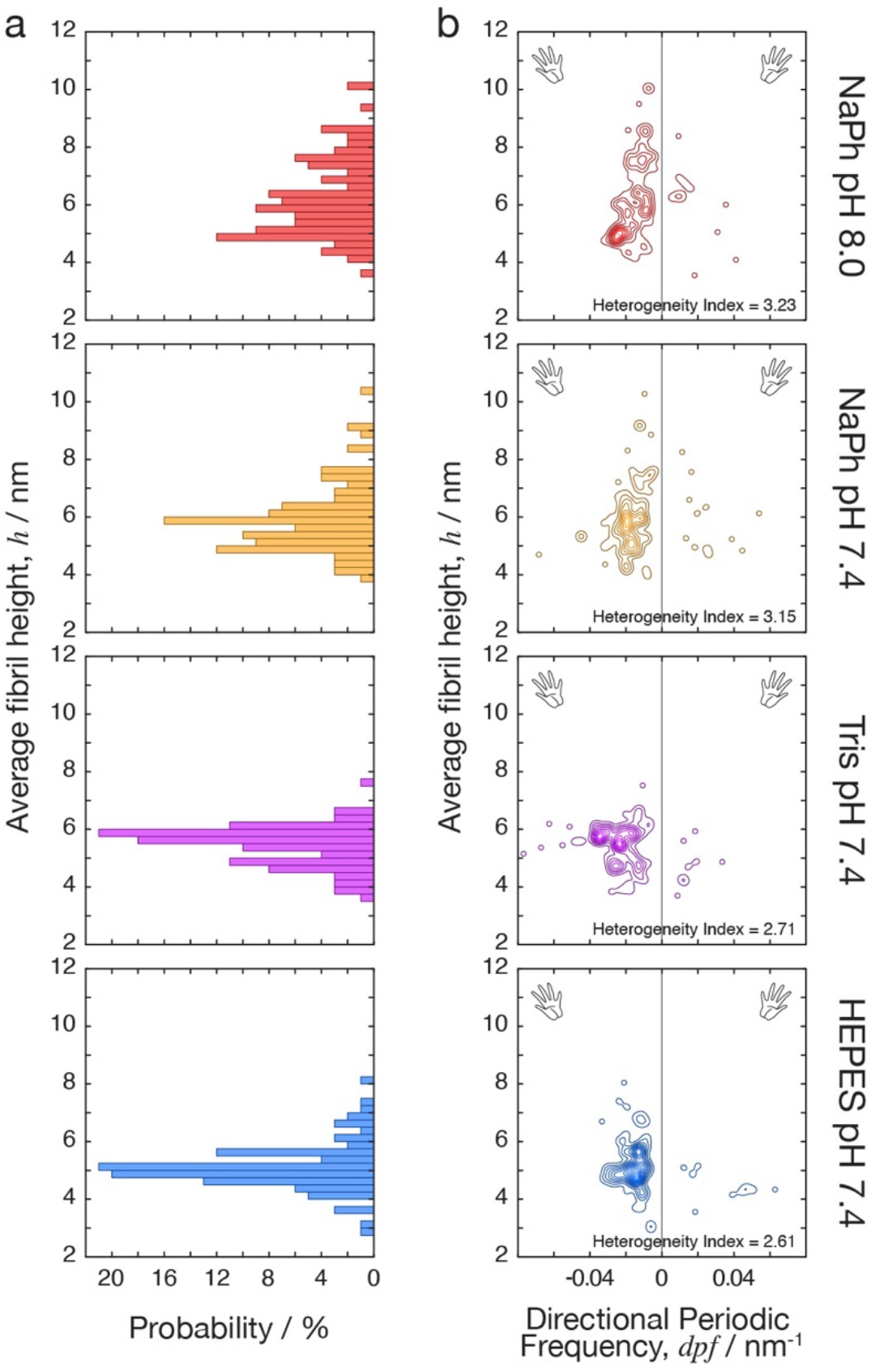
The distribution of structural polymorphs of Aβ_42_ fibrils is sensitive to the assembly solution condition. Morphometric analysis of Aβ_42_ fibrils formed in 20mM sodium phosphate (NaPh) pH 8.0, sodium phosphate pH 7.4, Tris pH 7.4, and HEPES pH 7.4, respectively. (a) The distribution of average fibril height for the heterogeneous fibril populations formed in each of the assembly conditions tested. (b) Contour maps displaying average height of individual fibrils (describing their cross-sectional width) plotted versus their individual *dpf* values (describing their helical properties of handedness and pitch). The structural variation of Aβ_42_ fibrils formed in each of the four assembly conditions are visualised and compared through these contour maps by spreading the characteristics of their structures onto two dimensions. The heterogeneity index, HI (larger value indicating higher heterogeneity as defined in the methods section), for each of the population distributions is also shown.

To visualise, organise and identify natural partitions within the single-fibril level structural data that may represent distinct classes of fibril structures, we carried out an agglomerative hierarchical clustering analysis ^29^. For this structure based clustering analysis, we developed a distance function *d_ξ_* (see Methods) that takes into account the detailed cross-sectional shape ^35^ as well as the helical symmetry properties described by *dpf* parameter ^29^. This pairwise *d_ξ_* distance score, where low numerical value indicate high degree of structural similarity and high numerical value indicate dissimilarity for a pair of individual fibrils, were calculated for all 79,800 pairings possible for 400 individual fibril structures analysed. This approach allowed us to quantify and objectively compare the heterogeneity of each of the four fibril populations, and to detect if the fibril structures can be divided into classes that correlate with the assembly conditions.

For quantitative comparison of the magnitude of heterogeneity of the four fibril populations, we defined a heterogeneity index: *HI* = *RMS_dξ_*, based on the quadratic mean of *d_ξ_* distance for all fibril pairs within each sample population. The HI value of the observed fibril population at each assembly condition was subsequently calculated and ranked objectively, revealing the order of high to low heterogeneity: sodium phosphate buffer at pH 8 (HI = 3.23), sodium phosphate buffer at pH 7.4 (HI = 3.15), Tris buffer at pH 7.4 (HI = 2.71) and HEPES buffer at pH 7.4 (HI = 2.61). These result corroborated the qualitative assessment of fibril population heterogeneity seen in the height vs. *dpf* contour map analysis (**Figure 3b**).

In terms of the species present in each of the four fibril populations, the agglomerative hierarchical clustering analysis of the entire dataset of 400 fibrils using the pairwise *d_ξ_* distance measure (**Figure 4a**) demonstrated that fibrils of similar structures that may constitute a distinct fibril polymorph class can often be found in more than one condition examined. This can be seen in the colour bar to the right of the dendrogram in **Figure 4a**, showing that each cluster in the data with distance *d_ξ_* < 1 is almost always composed of individual fibrils observed from two or more assembly conditions. This result, again, corroborates with the qualitative assessment of the fibril populations seen in the height vs. *dpf* contour map analysis (**Figure 3b**) where there is a considerable overlap of fibril structures observed in a region of the maps with left-handed twist with 10-40 repeating units per µm (0.01-0.04 repeating units per nm) and an average fibril height between 4-6 nm across the different conditions tested. Whereas the coarse contour map analysis is not able to separate fibril structures with further detail than their width and helical morphometrics, the hierarchical clustering analysis of 3D-reconstructed filament envelopes is able to analyse and cluster filaments based on their cross-sectional shapes in addition to their helical pitch and handedness (**Supplementary Figures S4**). Thus, this analysis suggests that some fibrils of similar structures that may constitute a common fibril polymorph were able to form in some or all of the buffer salt or pH conditions examined. Overall, these results allow us to confirm, in a quantifiable manner, that Aβ_42_ fibrils are able to form a cloud of different structural polymorphs under identical conditions in the same sample, and that each and every fibril displays structural individuality, as previously seen in the assembly of short amyloid forming peptides ^29^. These results also confirm the observations made in earlier works that Aβ_42_ amyloid fibrils are highly polymorphic, and the amyloid sample populations are highly heterogeneous *in vitro* and *ex vivo* from patients ^6, 16, 22^.

**Figure 4.**
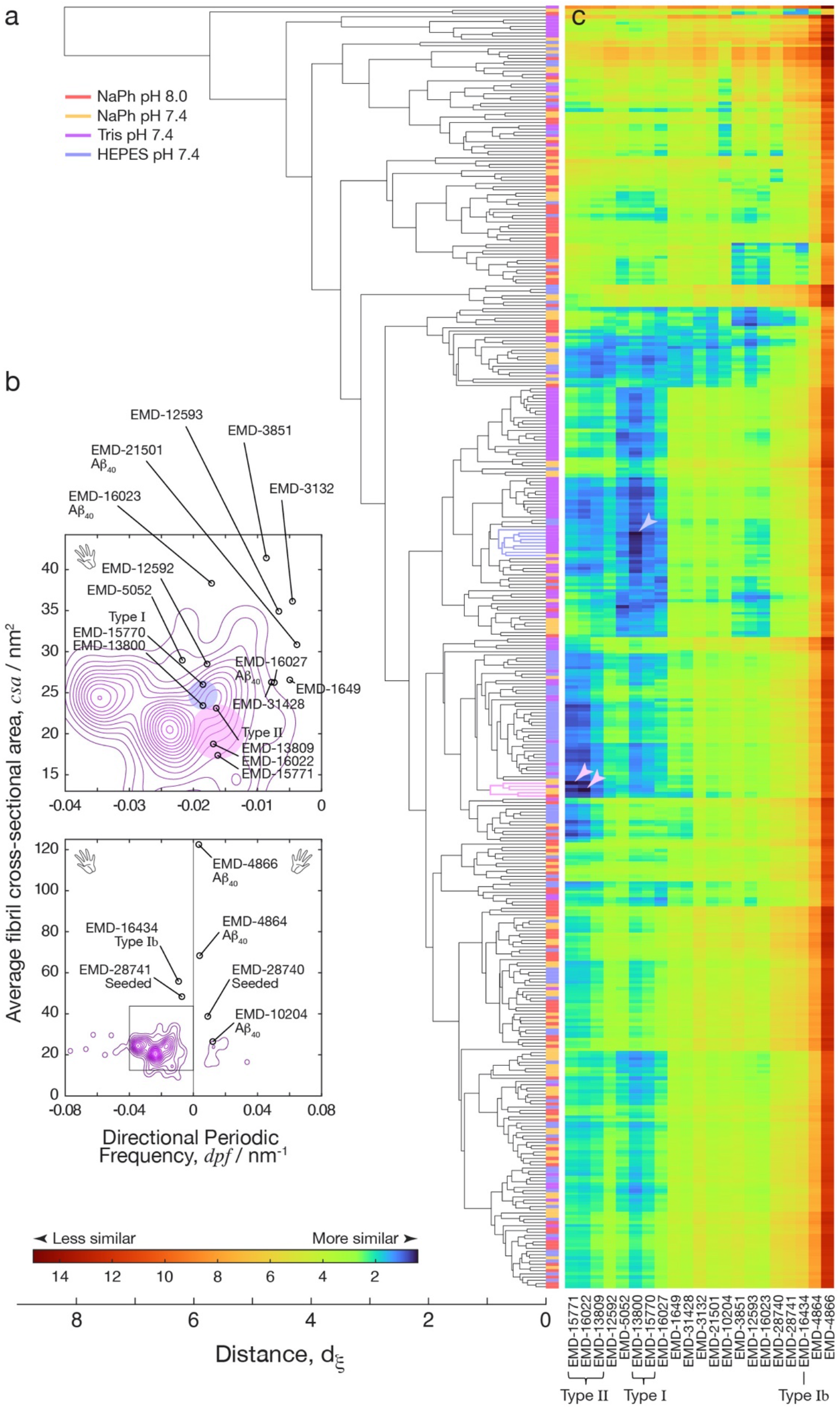
Comparison of individual Aβ_42_ fibril structures observed by AFM to cryo-TEM derived structural maps suggest rare fibril structures in the heterogeneous populations match fibril polymorphs seen in human patient brain samples. (a) Agglomerative hierarchical clustering analysis of all 400 individual fibrils analysed from the four assembly conditions. This analysis organises the fibril based on their individual surface envelopes obtained from 3D reconstructions using individual filament level AFM data, with the order of the filament from bottom to the top of the dendrogram matching the order of individual fibrils in **Supplementary Figures S2, S3 and S4**. Colour bar to the right of the dendrogram indicate the origin of each of the filaments in terms of their assembly condition, indicating that fibril polymorphs were generally observed across the experimental assembly conditions tested. (b) Contour maps displaying average AFM tip-accessible cross-sectional area of individual fibrils (csa, describing their cross-sectional area) plotted versus their individual *dpf* values (describing their helical properties of handedness and pitch) for the polymorph distribution of fibrils assembled in Tris pH7.4. The cross-section area (*csa*) and the directional periodic frequency (*dpf*) of cryo-TEM derived Aβ_40_ and Aβ_42_ structural data entries in the EMDB are also mapped onto the contour maps for comparison. Type I and Type II Aβ_42_ fibril polymorphs from human brains ^6^are highlighted by light blue and pink circles, respectively. The bottom contour map is a zoom-out of the region represented by top map highlighted by black square. (c) The similarity between each of the 400 individual fibrils and the cryo-TEM derived structural maps quantified by the pairwise *d_ξ_* distance scores. All of the pairwise scores are visualised by the colour scale shown to the left, with low *d_ξ_* distance scores indicating high similarity. The rows of this image represent each of the 400 fibrils in the same order as shown in the dendrogram in (a) and the columns represent an EMDB-entry with their identity labelled at the bottom. The identity of known Aβ_42_ fibril polymorphs purified from human brains are also indicated. Arrow heads show the top three ranked matches, and the light blue or pink colour of the arrow heads and clusters refer to matches with Type I and Type II Aβ_42_ fibril polymorphs ^6^, respectively.

### The *in vitro* population cloud of polymorphic Aβ_42_ structures encompasses fibril structures found from *ex vivo* human patient brains

Finally we investigated if the heterogeneous Aβ_42_ fibril populations we observed encompasses amyloid fibril structures that can be found in humans. There are now several different experimentally determined structural models describing the filament core structures of Aβ_42_ fibrils. In particular, recent reports detail cryo-TEM based structure determination that revealed the core structures of several Aβ_40_ and Aβ_42_ fibril polymorphs from brain samples of human patients suffering from Alzheimer’s disease and related dementias in atomic detail ^6, 22–26^. In order to validate whether the maps of the polymorphic Aβ_42_ assembly landscape we observed encompasses known Aβ_42_ polymorphs characterised to high detail by cryo-TEM, we calculated the distance score, *d_ξ_* (see Methods), between every one of the 400 fibril structures we 3D-reconstructed and analysed by AFM, and all of the recent Aβ_42_ cryo-TEM maps (**Table 1**) deposited in the Electron Microscopy Data Bank (EMDB). We also included recent cryo-TEM maps of Aβ_40_ fibrils found in the EMDB as controls since current evidence strongly shows that fibrils formed of Aβ_42_ are different to those of Aβ_40_ (**Table 1**). Comparison and structural distance calculations between the Coulomb potential maps from cryo-EM experiments in the EMDB and the individual filament envelopes from our AFM data was accomplished by modelling the rigid-body interaction of an AFM tip of average sharpness used in our experiments with each of cryo-TEM map to generate simulated AFM images of the fibril polymorphs seen by cryo-TEM (**Figure 5a**). The shapes of the external envelope of the fibrils were determined by calculating the contact points between a simulated AFM tip of average sharpness and the author-recommended iso-surface of the cryo-TEM derived structural maps that are denoised, axis-aligned and extended using the reported twist and rise information in each of the EMDB entries ^35^. These simulated images demonstrate the considerable variety of the Aβ structures and the considerable morphometric differences between each. The twist and rise information was also used to calculate the *dpf* values of each cryo-TEM derived data entry. In **Figure 4b**, a scatter plot of these data is overlaid onto the assembly contour map from all the individual fibrils formed in the Tris buffered condition used here. Interestingly, many cryo-TEM derived entries, including the Aβ_40_ entries, do not fall within the contour boundaries, but Type I and Type II Aβ_42_ polymorphs observed from human brain samples were located well within the mapped polymorph contour boundaries.

**Figure 5.**
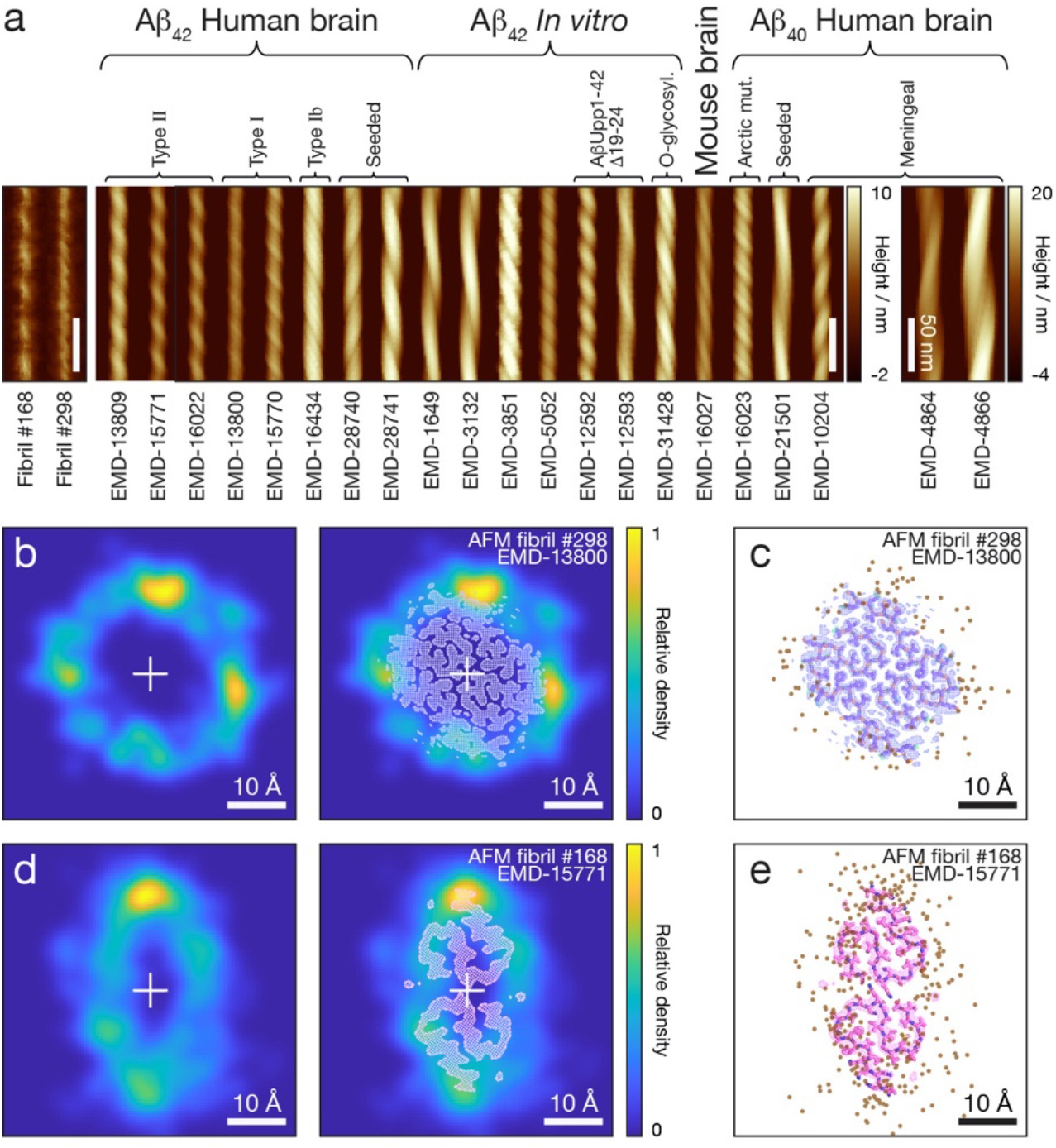
Structural comparison of Type I and Type II Aβ_42_ fibril polymorphs from human patient brain samples with their respective best matched individual fibrils seen in the heterogeneous fibril populations. (a) Comparison of the straightened AFM height image of the two Aβ_42_ fibrils best matched to cryo-EM derived structural maps with simulated topographical AFM images of Aβ amyloid fibril polymorph data entries found in the EMDB. The Aβ fibril polymorphs were simulated with a tip radius of 5.5 nm matching to the average tip radius estimate of the AFM data using the iso-surfaces of axis-aligned and extended cryo-EM structural maps. A 180 nm segment is shown for each fibril image and the scale is identical for all images except EMD-4864 and EMD-4866 due to their large cross-section as indicated by their separate colour scale. The scale bars indicate 50 nm in all images. (b) AFM cross-sectional contact-point density map of the Aβ_42_ fibril best matched to Type I polymorph from human patient brains ^6^. The AFM-derived density map is shown with and without the cross-section of cryo-TEM derived structural map superimposed for comparison. (c) The fibril cross-section of the Type I polymorph from human patient brains shown as cryo-TEM Coulomb potential map superimposed with the fitted structural model (PDB: 7Q4B) together with the cross-sectional coordinates of AFM tip-filament contact-points (dots) along the top of the fibril contour deconvoluted and 3D-reconstructed from the topographical AFM data. The RMSD between the tip-accessible cross-sectional cryo-TEM derived iso-surface and AFM-derived surface envelope is 1.38Å. (d)-(e) The same as (b) and (c) but for the Aβ_42_ fibril best matched to Type II polymorph from human patient brains (EMD-15771 and PDB: 8BFZ ^6^) with a RMSD between the tip-accessible cross-sectional cryo-TEM derived iso-surface and AFM-derived surface envelope of 1.65Å. All scale bars in (b)-(e) indicate 10Å in length.

**Table 1:** Cryo-TEM structural data entries of polymorphic Aβ amyloid fibrils in the EMDB. Structural maps of Aβ_42_ amyloid fibrils from 2009 and recent Aβ_40_ amyloid fibrils from 2019 with resolution around 7Å or better in the EMDB included in this study are listed in the order of release year.

| EMDB ID | 40/42 | Fibril type | Resolution / Å | Year Released | Publication |
| --- | --- | --- | --- | --- | --- |
| EMD-1649 | Aβ<sub>42</sub> | in vitro | 15 | 2009 | Schmidt et al., 2009 <sup>19</sup> |
| EMD-5052 | Aβ<sub>42</sub> | in vitro | 10 | 2010 | Zhang et al., 2009 <sup>20</sup> |
| EMD-3132 | Aβ<sub>42</sub> | in vitro | 7 | 2015 | Schmidt et al., 2015 <sup>18</sup> |
| EMD-3851 | Aβ<sub>42</sub> | in vitro | 4 | 2017 | Gremer et al., 2017 <sup>21</sup> |
| EMD-12592 | Aβ<sub>42</sub> | in vitro, AβUpp1-42Δ19-24 | 5.7 | 2021 | Pagnon de la Vega et al., 2021 <sup>68</sup> |
| EMD-16434 | Aβ<sub>42</sub> | Human brain | 3.5 | 2022 | Yang et al., 2022 <sup>6</sup> |
| EMD-12593 | Aβ<sub>42</sub> | in vitro, AβUpp1-42Δ19-24 | 5.1 | 2022 | Pagnon de la Vega et al., 2021 <sup>68</sup> |
| EMD-31428 | Aβ<sub>42</sub> | in vitro, dsAβ<sub>42</sub> Tyr10 O-glycosylation | 3.1 | 2022 | Liu et al., 2021 <sup>69</sup> |
| EMD-28740 | Aβ<sub>42</sub> | Human brain, seeded | 2.83 | 2022 | Lee et al., 2023 <sup>22</sup> |
| EMD-28741 | Aβ<sub>42</sub> | Human brain, seeded | 2.76 | 2022 | Lee et al., 2023 <sup>22</sup> |
| EMD-15770 | Aβ<sub>42</sub> | Human brain | 2.9 | 2022 | Stern et al., 2023 <sup>23</sup> |
| EMD-15771 | Aβ<sub>42</sub> | Human brain | 3.7 | 2022 | Stern et al., 2023 <sup>23</sup> |
| EMD-13800 | Aβ<sub>42</sub> | Human brain | 2.5 | 2022 | Yang et al., 2022 <sup>6</sup> |
| EMD-13809 | Aβ<sub>42</sub> | Human brain | 2.8 | 2022 | Yang et al., 2022 <sup>6</sup> |
| EMD-16022 | Aβ<sub>42</sub> | Human brain, E22G | 2.8 | 2023 | Yang et al., 2023 <sup>24</sup> |
| EMD-10204 | Aβ<sub>40</sub> | Human brain, meninges | 4.4 | 2019 | Kollmer et al., 2019 <sup>25</sup> |
| EMD-4864 | Aβ<sub>40</sub> | Human brain, meninges | 5.56 | 2019 | Kollmer et al., 2019 <sup>25</sup> |
| EMD-4866 | Aβ<sub>40</sub> | Human brain, meninges | 7.01 | 2019 | Kollmer et al., 2019 <sup>25</sup> |
| EMD-21501 | Aβ<sub>40</sub> | Human brain, seeded | 2.77 | 2021 | Ghosh et al., 2021 <sup>26</sup> |
| EMD-16023 | Aβ<sub>40</sub> | Human brain, E22G | 1.99 | 2023 | Yang et al., 2023 <sup>24</sup> |
| EMD-16027 | Aβ<sub>40</sub> | AppNL–G–F mouse brains, E22G | 3.5 | 2023 | Yang et al., 2023 <sup>24</sup> |

To individually match the 400 AFM-derived 3D-reconstrructed fibril structures to the cryo-TEM derived data (**Table 1**), the *d_ξ_* distance values were subsequently calculated and ranked for all 8400 pairwise comparisons between the 21 cryo-TEM derived structural maps and 400 individual filament envelopes reconstructed from our AFM data (**Figure 4c**). Strikingly, of the top 3 out of the 8400 ranked pairs showing the highest pair-wise similarities with lowest *d_ξ_* distance scores below 0.5, two are individual fibrils that match to the Type II polymorph structure and one is an individual fibril that matches to the Type I polymorph structure of Aβ_42_ seen in the brain samples from human patients ^6, 23, 24^ (arrowheads in **Figure 4c**). The clusters of fibrils (defined in the Methods section) containing the top matched individuals (**Figure 4a** blue and purple clusters matching to Type I and Type II human brain Aβ_42_ polymorphs, respectively) are all ranked in the top 20 for each matched cryo-TEM entry. The individual fibril and its parent cluster that matches to the Type I polymorph (**Figure 4a** blue cluster) account for 2.25% (9 out of 400) and the two individual fibrils and their parent cluster that matches to the Type II polymorph (**Figure 4a** purple cluster) account for 1.25% (5 out of 400) of the individual fibril observations. Thus, these clusters represent rare sub-populations within the heterogeneous Aβ_42_ amyloid fibril population clouds formed *in vitro* under the assembly conditions used that closely resemble Aβ_42_ fibril structures found in human patient brain samples.

To further validate the structural similarity between the rare fibril species found in the heterogenous *in vitro* assembled fibril populations and the Aβ_42_ fibril structures from human patient brain samples, the top matched individual fibril seen with AFM were further validated by comparing their fibril images with all of the simulated images made using cryo-TEM derived structural maps (**Figure 5a**). The cross-sectional contact-point density maps of the best matched individuals were also overlaid with the cross-sections of cryo-TEM derived structural maps of Type I and Type II Aβ_42_ amyloid from human brain samples and their fitted molecular models, respectively (**Figure 5b-e**), demonstrating the remarkable similarities. Interestingly, no other of the 400 individually analysed fibril structures by AFM matched well with any other cryo-TEM derived structural maps included in the analysis. Importantly, no Aβ_40_ structural matches were identified. In addition, the individual fibril 3D-structural reconstruction were independently and blindly performed since the original AFM image data were acquired prior to 2021 while the closely matched cryo-TEM data entries were released in 2022 and 2023. Therefore, despite that only the surface envelopes of the individually analysed fibrils were reconstructed and matched, these results taken together demonstrate strong evidence suggesting that the synthetic fibrils formed *in vitro* from recombinant Aβ_42_ can represent, in rare parts, disease relevant fibril structures formed *in vivo*. These conclusions demonstrate that a common set of polymorphs, including disease related polymorph structures, do populate and overlap across the different condition *in vivo* and *in vitro* despite the sensitivity of the assembly landscape.

## Discussion

When compared to conventional protein folding, in which structural polymorphism is rare, amyloid assemblies demonstrate a remarkable ability to have a single sequence of amino acids fold and assemble in numerous ways to make up the cross-β core regions of fibrils. One clear example is the range of *ex vivo* fibril structures reported from various patients with different diseases in which tau fibril formation is reported ^39^. Structural variation could be an important factor in amyloid disease as distinctive structures could arise from different local conditions, requiring different treatment options dependent on the local environment and the structure formed in that disease. Equally, different structural polymorphs could each have a distinctive set of biophysical and biochemical properties, display dissimilar surfaces and therefore contribute to the local biological environment in different ways, therefore having diverse impacts in different diseases. Finally, structural variation itself may result in a dynamic and continuously evolving cloud of fibril species which could result in a variety of behaviours and can change upon changes to the local environment, making different disease states difficult to predict and to target. It is already known that Aβ can form polymorphic structures which has been demonstrated in both *in vitro* and *ex vivo* samples (**Table 1**). However, the polymorphic extent and the heterogeneity of Aβ fibril assembly has not been quantified to establish the ruggedness of the landscape of polymorphic Aβ assembly. Furthermore, the effect of changing the assembly conditions on Aβ polymorphism has not previously been quantified despite a plethora of different conditions were used in the literature to date (**Figure 1c** and **Supplementary Table 1**). Finally, it remains hitherto unclear as to whether a subset of polymorphs is conserved across conditions *in vitro* and *in vivo* or whether changing the assembly conditions resulted in entirely different set of structures.

Due to its importance in Alzheimer’s disease, Aβ_42_ is a highly researched amyloidogenic peptide sequence. In terms of the amyloid fibril structures, numerous different structural polymorphs have been determined from various cryo-TEM and ssNMR studies. Rather than adding evidence to confirm and to improve previous structural models of Aβ_42_ amyloid, each new study has often revealed entirely new structures, despite investigating the same primary amino acid sequence. This has cemented the idea that Aβ_42_ amyloid assembly can be highly polymorphic. Differences in the assembly conditions used *in vitro* have often been cited as contributing factors towards the variations in experimental results. Unsurprisingly, different experiments can often require different experimental conditions. For example, the strength and integrity of the signal obtained using circular dichroism is highly dependent on the solution used ^44^. Differences in the type of experiment being performed, along with research group specific preferences have resulted in a vast array of experimental conditions in which Aβ_42_ polymerisation has been studied (**Supplementary Table S1**). Therefore, in this work, we sought to understand how different experimental conditions contribute to the structural differences of the fibrils generated as well as the heterogeneity of the fibril populations formed. To do so, we have collated commonly used experimental conditions from highly cited publications from the last 15 years involving Aβ_42_ assembly and developed a unique method to quantify, by structural analysis at individual fibril level, the extent of structural polymorphs that arose when forming fibrils using the different conditions adopted by the field.

We have observed a highly polymorphic landscape of structural possibilities for Aβ_42_ fibril formation which is not simply constrained to one or few types of structural polymorph per assembly condition. Both right and left-hand twisted fibrils were also observed under all experimental conditions although left-handed fibrils account for around 90% of the individual filaments we analysed. Whilst phosphate buffers appeared to promote variation in fibril width, the Tris buffered condition promoted variations in the helical twist. These data have confirmed that Aβ_42_ fibril samples are highly polymorphic, and its rugged polymorphic landscape of assembly is surprisingly sensitive to assembly condition, as the polymorph distribution can be shifted by small changes to the assembly conditions both in terms of the species formed and the overall heterogeneity of the population distribution.

The supra-structural features of amyloid fibril populations, including both the individual fibril level structural parameters and the population level interconnectivity, are likely to impact upon their biological activity. For example, filaments with a larger surface area to volume ratio could be more efficient for secondary nucleation or other surface catalysed reactions resulting in the formation of toxic oligomeric species ^45^. Thin fibrils with a low frequency of twisting may be susceptible to breaking and therefore fragment more frequently, which is known to increase the rate of amyloid uptake into cells ^46^. Indeed, different polymorphic structures resulting from the same amyloid forming protein may considerably alter the fibrils’ growth, division by fragmentation and chaperone interactions ^47–49^. Here we observe variations in Aβ_42_ supra-structural features that are dependent on the experimental conditions. Phosphate buffer, in particular at pH 8.0, promotes a higher degree of inter-protofilament association and ultimately thicker and wider fibrils. Since phosphate is a polyprotic base, it could provide greater local electrostatic shielding between filaments compared to the other two buffer salts at the same molar concentration, resulting in some thicker fibrils, similar to the way in which high ionic strength solutions have been reported to shield electrostatic repulsion between monomers and fibrils in turn catalysing secondary nucleation effects ^50^. Fibrils formed in Tris buffer displayed more variable twist patterns but lesser variation in average height between fibrils. This could be due to perturbed intra-filament interactions resulting in no single favourable periodicity in these conditions which in turn could result in fewer thicker filaments as favourable orientations for inter-filament interactions may be reduced. In this context, secondary nucleation is likely to be impacted by the molecular surface differences due to the supra-structural features of the fibril populations. Here, our data indicate that the supra-structure, including structural variation between filaments and heterogeneity of the fibril population, is sensitively influenced by the experimental conditions. Therefore, it is reasonable to assume that differing conditions, for example in different brain regions, can alter the pathological contribution of Aβ_42_ fibrils by means of changing their supra-structures. Toxicity of Aβ_42_ amyloid species is thought to be size dependent. Typically, smaller species or small fibril fragments rather than large fibrils or plaques are considered to more acutely toxic ^3, 32, 38, 51^. Furthermore, a general feature of amyloid fibrils is that fragmented fibrils can produce toxic effects ^32, 52, 53^ although intermediate oligomeric species of Aβ_42_ may yet possess a more potent cytotoxic potential ^54^. These cases highlight the potential for Aβ fibrils to act as a long-term source of small species with cytotoxic potential. Aβ_42_ fibrils can also induce a positive feedback loop of increased toxic potential through secondary nucleation events that results in smaller species capable of toxic effects ^55^. However, the relationship between the structure of Aβ_42_ fibril polymorphs and their function with respect to their individual capacity for fragmentation or catalysing secondary nucleation is not well understood. Amyloid fibrils have been suggested to show prion-like properties in which they propagate from cell to cell ^56–58^. Examples of this behaviour is analogous to epigenetic gene regulation in yeast by yeast prions ^57, 59^ and the spread of toxic particles in various prion diseases such as Creutzfeldt-Jakob disease ^60^. Aβ_42_ has also been shown to be able to spread in a prion-like manner ^61^. Prion structures that spread can have a structural dependence, resulting in some cells populated with different ‘strains’ of fibrils ^56–58, 62, 63^. Structural polymorphism can therefore have an impact on the prion-like behaviour as higher variation in the fibril populations could result in a higher likelihood of fibrils proliferating with structural features which promote fragmentation and/or spreading. Smaller particles for example have been observed to link with the infective potential of prion particles ^46^ indicating that having at least one subset of structural polymorphs which are prone to fragmentation in a heterogeneous polymorph distribution may increase the infective potential of a population of amyloid fibrils. Hence, using experimental conditions that encourage populations of fibrils that have a thin, easily broken morphology could result in significantly different outcomes in experiments measuring the spread of Aβ_42_ between cells.

Amyloid fibrils from the same precursor protein can be implicated in numerous diseases. One example of such is Tau which is implicated in numerous tauopathies, including in Alzheimer’s disease. Cryo-TEM of *ex vivo* Tau fibrils from patients with different tauopathies has demonstrated that the core structure of tau fibrils varies in a disease dependent manner ^39^. From current structural evidence, it is not yet clear if Aβ, which is also associated with several different diseases as well as Alzheimer’s disease, demonstrates similar behaviour compared to tau in forming disease specific structures, or if Aβ fibrils formed in different brain regions form specific structures. It is also not clear if the lower levels of heterogeneity and structural variations in amyloid populations seen in the structural studies of *ex vivo* brain samples (**Table 1**) compared to the high level of structural variations observed here are due to specific difference in the *in vivo* biological environments, long timescale properties of the assembly mechanism or methodologically linked observation biases. For example, a recent study where Aβ fibril plaques from mouse brains were imaged *in situ* by cryo-electron tomography revealed that, whilst fibrils extracted from the brains had the same atomic structure solved by cryo-TEM as those previously identified, *in situ* the observed fibrils were quantifiably thinner indicating the presence of a different polymorph that existed *in vivo* ^64^. In early-stage Alzheimer’s disease the hippocampal region is considered to be particularly at risk before other regions of the brain become impacted in later stages ^65^. Structural features such as the ability to fragment and spread or act as a catalysing surface for secondary nucleation or other surface interactions could contribute to the biological behaviours of amyloid populations propagating from early stages to late stages of disease. Structural variation in Aβ fibrils could increase the likelihood of fibrils that are more prone to spreading resulting in the deposition of Aβ plaques found throughout the brains of patients with late-stage Alzheimer’s disease.

In summary, we have demonstrated that Aβ_42_ amyloid fibril populations are highly polymorphic and sensitive to assembly conditions. We have also demonstrated that by applying AFM imaging and individual filament 3D structural analysis by CPR-AFM, the structural variations, polymorphism and heterogeneity within amyloid populations can be quantified. Furthermore, we present evidence that rare species in the heterogeneous fibril populations formed *in vitro* matches those observed in *ex vivo* human patient brain samples, giving support to the view that disease and physiologically relevant amyloid species can be formed *in vitro*. The approach we present gives complimentary population level information that links to high-detailed structural analysis methods such as cryo-TEM and ssNMR which can be used to determine the average core structures of one or few dominant Aβ_42_ polymorphs to atomic detail ^35, 66^. Small changes in assembly conditions applied here revealed that the rugged landscape of polymorphic amyloid assembly can easily be shifted, although our data demonstrate that a common set of polymorphs, including disease related polymorph structures, can exist across different conditions despite the sensitivity of the polymorph distribution. Therefore, understanding the precise *in vitro* conditions that generate *in vivo* disease associated structures in rarity or in dominance will be important for resolving the assembly mechanisms of these structures in a medical disease setting and providing valuable experimental tools for diagnostics and therapeutics research and development. Overall, the conclusions from this study link *in vitro* and *in vivo* structural data and put into focus the structural variations, diversity, heterogeneity and molecular population level properties of amyloid in general. Determining the factors that control the formation of dominant amyloid polymorph structures, control the formation of rare species and control the extent of structural variations and population heterogeneity, are all required for a fuller understanding of the roles that amyloid molecular populations play in biology, as cause or consequence of disease or as structures required for functions.

## Materials and Methods

### Expression, purification and characterisation of monomeric Aβ_42_

Recombinant Aβ_42_ was expressed and purified as described elsewhere ^36, 37^. Briefly, *E. coli* BL21 cells were transformed with the pET-sac Aβ_42_ plasmid. Colonies were grown on and selected from LB-agar plates to form starter cultures in LB media. 1 L volumes of culture were then grown until they reached an OD_600_ of 0.6 at which point protein expression was induced using IPTG and the cultures were left for 4 more hours before cell pellets were harvested by centrifugation. The protein was expressed in 4 x 1 L batches and purified 1 L at a time. Aβ_42_ containing inclusion bodies were purified by sonicating frozen cell pellets in 20 mM Tris buffer at pH 8 followed by centrifugation at 18000 x g. The inclusion bodies were further sonicated in Tris buffer to dissolve the pellet followed by immediate immersion in 8M UREA to induce denaturing conditions. The protein was purified by ion exchange chromatography using DEAE sepharose followed by size exclusion chromatography using a 60 ml Superdex 75 increase column (GE healthcare) in 6M GuHCl. Chromatograms and SDS-PAGE data (**Supplementary Figure S1a and b**) provided evidence of purified monomeric Aβ_42_. Finally, immediately before use, monomeric Aβ_42_ was further purified using a 30 ml Superdex 75 increase column (GE healthcare). Alternately, several aliquots of Aβ_42_ were concentrated using centrifugal filtration units (Amicon) and then purified using the 30 ml Superdex 75 increase column twice. Either way the sample was eluted into the appropriate buffer solution of 20mM sodium phosphate (NaPh) pH 8.0, sodium phosphate pH 7.4, Tris pH 7.4 or HEPES pH 7.4 (at pH 8.0 followed by adjustment to pH 7.4 if necessary once eluted from the column). The ability of the expressed and purified Aβ_42_ monomers to assemble and form β-rich and ThT positive amyloid fibrils was confirmed by CD and ThT kinetics assay, respectively (**Supplementary Figure S1c and d**). To assess the cytotoxic potential of the Aβ_42_ samples, monomeric protein solutions were diluted to a concentration of 50µM in 10mM HEPES buffer (10 mM HEPES, 50 mM NaCl, 1.6 mM KCl, 2 mM MgCl_2_, 3.5 mM CaCl_2_) and left at room temperature for 2–3 hours to oligomerise as detailed in earlier publication ^38^. These were added to cell culture media of mouse primary hippocampal neurons to a final concentration of 10µM and the cytotoxicity effect of the samples on the cells was assayed after 1 week using ReadyProbes^TM^ (Thermofisher). Cells were incubated for 15 mins with one drop each of ReadyProbes ^TM^ blue (all cells) and green (dead cells) and imaged using a Zeiss CO widefield microscope with DAPI and FITC filters, and the cytotoxic potential of the oligomeric species was shown to be comparable to a commercially produced Aβ_42_ peptide purchased from rPeptide (**Supplementary Figure S1e**).

### Analysis of published experimental conditions for amyloid assembly

A literature analysis was conducted where the most highly cited publications involving at least one experiment in which monomeric Aβ_42_ was present and assembled in some capacity, from 2005 to 2020 were identified. The experimental conditions from each of the identified publications was tabulated (**Supplementary Table S1**). These conditions were analysed, the types of experimental buffers used were annotated and their relative frequencies in the publication dataset quantified. Specifically, buffers that were intended as experimental buffers for Aβ_42_ polymerisation were recorded. For example, for the frequently employed strategy in which monomeric peptide was solubilised in NaOH or DMSO before being diluted into an experimental buffer or cell culture media, the particular buffer or media were included in the analysis not the initial step involving NaOH or DMSO. Another commonly used method is to use a “vehicle” buffer in which monomer is initially incubated before being transferred to cell culture. In this case the “vehicle” buffer *would* be recorded as it is the initial condition in which substantial polymerisation could occur. Included in the tabulated publication data but not the analysis are publications detailing Aβ_42_ fibril structures that can be found in the PDB but did not meet the criteria for being one of the most highly cited in the year it was published. The literature search was performed initially using Web of Science, followed by validation by cross-referencing using Google Scholar.

### *In vitro* fibril formation

To prepare amyloid fibril samples formed from the recombinant Aβ_42_, the monomer solution eluted from the final step of purification was immediately incubated at 37 °C without agitation. The solution was incubated for at least 1 week before being sonicated for 5 seconds. Freshly sonicated fibril fragments were subsequently used as seeds and were mixed with freshly prepared monomer to a final concentration of 15 µM monomer equivalent concentration with 1 % total protein mass in pre-formed seed fibrils. These second generation fibrils were incubated at 37 °C without agitation for at least 1 week in the appropriate buffer solution before used for AFM imaging.

### AFM specimen preparation and imaging

Each sample to be used for AFM imaging was diluted to 10 µM using a combination of the appropriate buffer and a solution of HCl that had been pre-determined to result in a final pH of 2 when mixed in a 1:20 ratio with the appropriate buffer for each sample. Immediately after dilution, 20 µl samples were deposited onto freshly cleaved mica surfaces (Agar scientific, F7013) and incubated for 5 minutes. This low pH deposition method was employed to adjust the surface charge of the mica surfaces to allow more efficient deposition of the fibrils. Following the 5 min incubation, the sample was washed with 1 ml of filter sterilised milli-Q water and then dried using a gentle stream of nitrogen gas. To confirm that the brief pH jump deposition method did not perturb the fibrils, fibrils that were subjected to the procedure were used to generate seeds for a seeded reaction in which the thioflavin T profile was found to be identical to that from a reaction using seeds made from fibrils which had not undergone the brief pH jump procedure. Fibrils were imaged using a Multimode 8 AFM with a Nanoscope V controller (Bruker) operating under peak-force tapping mode using ScanAsyst probes (silicon nitride triangular tip with tip height = 2.5-2.8 µm, nominal tip radius = 2 nm, nominal spring constant 0.4 N/m, Bruker). Each collected image had a scan size of 4 x 4 µm and 2048 x 2048 pixels or 8 x 8 µm and 4096 x 4096 pixels. Therefore, the same pixel density is maintained for all images within the dataset. A scan rate of 0.305 Hz was used for the 4 x 4 µm and 0.2 Hz for the 8 x 8 µm scans. A noise threshold of 0.5 nm was used, and the Z limit was reduced to 1.5 µm. When obtaining high-detail topographical maps (**Figure 2 and Supplementary Figure S2**), the fibrils were probed gently using ∼400 pN of force, which is substantially lower than the reported force required for the deformation of amyloid fibrils ^67^. Nanoscope analysis software (Version 1.5, Bruker) were used to process the image data by flattening the height topology data to remove tilt and scanner bow.

### Individual filament structural analysis

Aβ_42_ fibrils on the AFM height images were traced and digitally straightened using algorithms previously described ^34, 43^ and the height profile for each fibril was extracted from the centre contour line of the straightened fibrils. The periodicity of the fibrils was then determined using fast-Fourier transform of the height profile of each fibril. The final structural datasets consist of a varying multitude of images comprising of 100 individually traced fibrils per assembly condition used. In order to determine appropriate fibrils for analysis, selection criteria were applied in which single fibrils, with a free segment up to the point at which they either reached an end or overlapped with another fibril were used. Selected fibrils also had at least 3 repeating units and no visible breakages. Care was also taken to ensure that no more than one segment of the same fibril was included in the analysis unless a distinct change in morphology was observed in which case the segments were treated and treated as a separate fibrils. By excluding broken fibrils likely occurred upon deposition, it is possible that the polymorph distribution results were biased against thin filaments with a specific twist pattern.

Straightened fibril traces were corrected for the tip-sample convolution effect by CPR-AFM based on rigid-body geometric modelling of the tip-fibril contact points ^34, 43^. Briefly, the algorithm first corrects for the lateral dilation of nano-structures resulting from the finite dimensions of the AFM probe, without the loss of structural information, by resampling of the fibrils at tip-sample contact points. This results in recovering subpixel resolution of lateral sampling. Each pixel value in the straightened fibril data is then corrected in their x, y and z coordinates. Filament helical symmetry was estimated by building 3D models with various symmetries from the data, back calculating a dilated AFM image and comparing the angle of the fibril twist pattern with that of the straightened fibril trace in the simulated images and in the 2D Fourier transform of the simulated images. Then for construction of the 3D surface envelopes for each of the individual fibrils, the degree of twist per pixel along the y-axis was found by dividing 360° with the product of fibril periodicity and its symmetry number (e.g. 1 2 or 3 etc.). The 3D models were made assuming a helical symmetry using on a moving-window approach, in which a window, centred at a pixel *n*, contained the pixels *n – x* to *n + x* where *x* is the axial length covered by 180° twist. The central pixel *n* is not rotated while neighbouring pixels on both sides along the y-axis are rotated by a rotation angle, which is the product of the twist angle and the distance from n in pixels. Rotation angle values are negative in one direction from n and positive in the other direction, with the specific direction depending on the handedness of the fibril, determined by manual inspection of the straightened fibril image and its 2D Fourier transform image.

### Morphometric analysis of individual fibrils and polymorph distribution analysis

From each of 3D-reconstructed fibril models, several morphometric parameters ^29, 35^ were calculated. The average fibril heights, *h*, were measured on the central ridge of each fibril. The mean cross-over distance, *cod,* were measured between the peaks on the centre fibril height profile. The twist handedness of the fibril, *hnd*, is defined as −1 for left-hand twisted fibrils and +1 for right-hand twisted fibrils. The handedness of twist is determined for each filament by manual inspection of the straightened fibril image and its 2D Fourier transform image. The directional periodic frequency, *dpf*, is calculated as *dpf* = *hnd* / *helical pitch*. The filament mean AFM tip accessible cross-sectional area, *csa*, of the fibril is calculated by polar integration along the fibril axis of the reconstructed fibril 3D surface envelopes. The filament mean cross-sectional radius to the helical axis, *csr*, is calculated as the average distance from the filament’s AFM tip accessible surface to its helical axis. The filament cross-sectional mean second polar moment of area, *csjz*, is calculated by calculating the mean second moment of area with regard to the x and y-axis of the cross-section using the perpendicular axis theorem.

### Agglomerative hierarchical clustering

For the agglomerative hierarchical clustering analysis, a *L_1_* (Manhattan) distance measure, *d_ξ_*, was defined to measure the dissimilarity between a pair of fibrils *v* and *w*. The *d_ξ_* distance score takes into account the differences in the helical properties of the fibrils (along the helical z-axis) characterised by the *dpf* values, and the differences in the cross-sectional shape (in the x/y-cross-sectional plane) of the two fibrils Equation (1).

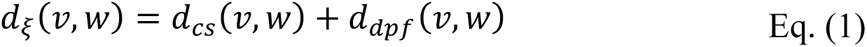

In Equation (1), the cross-sectional shape difference *d_cs_* is the standardised RMSD between the average tip-accessible fibril cross-sections defined according to Equation (2).

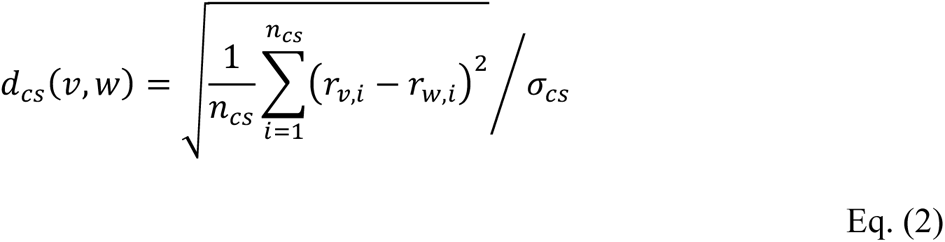

In Equation (2), *r* are the surface envelope coordinates in helical-polar coordinate system where the x-y axis plane rotates around the helical z-axis with the same periodicity as the helical periodicity of the individual helical filament. The parameter *n_cs_* is the number of sampling points, in this case, 1° intervals were sampled for a total of 360 points used. The parameter *σ_cs_* is the standardisation parameter, with the standard deviation of *d_cs_* of the whole dataset used here. The distance, *d_dpf_*, in Equation (1) is defined in Equation 3.

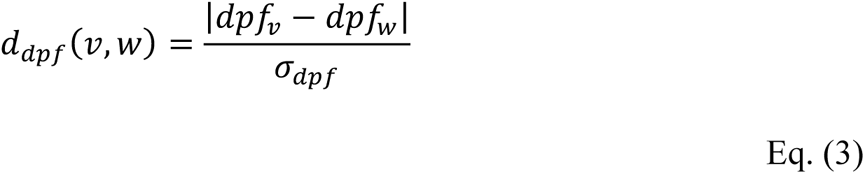

For the standardisation parameter *σ_dpf_* in Equation (3), the standard deviation of *d_dpf_* of the whole dataset used. The heterogeneity index HI is calculated over n*_dξ_* pairwise distances in a set of fibrils using Equation (4).

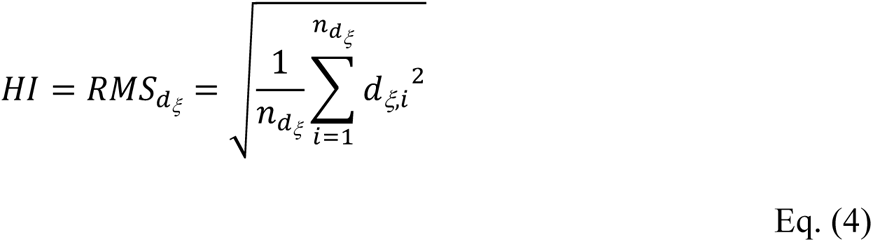

Agglomerative hierarchical clustering was performed using the average distance linkage function shown in equation (5) for cluster *r* and *s*.

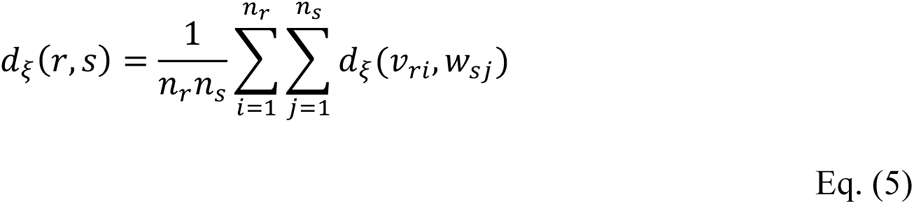

Explained briefly, the shortest distance, *d_ξ_*, between any two fibrils within a data set is found. Those two data points are then considered to be joined into a cluster and the two individual fibril data points are no longer used to determine the next cluster. Instead the average distance of both fibril data points to other fibrils or clusters are used. This is then repeated until all of the data is linked under one cluster.

### Comparative structural matching of individual fibril 3D-envelopes to cryo-TEM data

The pairwise similarity between each and every individually reconstructed 3D surface envelopes of the 400 Aβ_42_ fibrils analysed and cryo-TEM derived structural maps of Aβ_42_ and Aβ_40_ listed in **Table 1** were quantified. The structural maps were downloaded from the EMDB. The maps were axis aligned and their tip-accessible cross-sections calculated according to the method previously described ^35^ using an average tip radius of 5.5 nm. The pairwise *d_ξ_* distance scores in Equations (1)-(3) were subsequently calculated and ranked. The clusters identified to best match a cryo-TEM entry is found by finding individual fibrils with *d_ξ_* scores of less than 0.5 to the relevant cryo-TEM entry, and with other individual cluster members having *d_ξ_* scores of less than 1.0 to the same cryo-TEM entry and to each other within the cluster.

## Acknowledgements

We gratefully thank Sara Linse for the generous gift of Aβ_42_ expression plasmid used in this study. We thank the members of the Xue group and the Serpell group for helpful comments throughout the preparation of this manuscript. We also thank Ian Brown for technical support. This work was supported by funding from the Biotechnology and Biological Sciences Research Council (BBSRC), UK grant BB/S003312/1.

## Supplementary Information

**Supplementary Figure S1:**
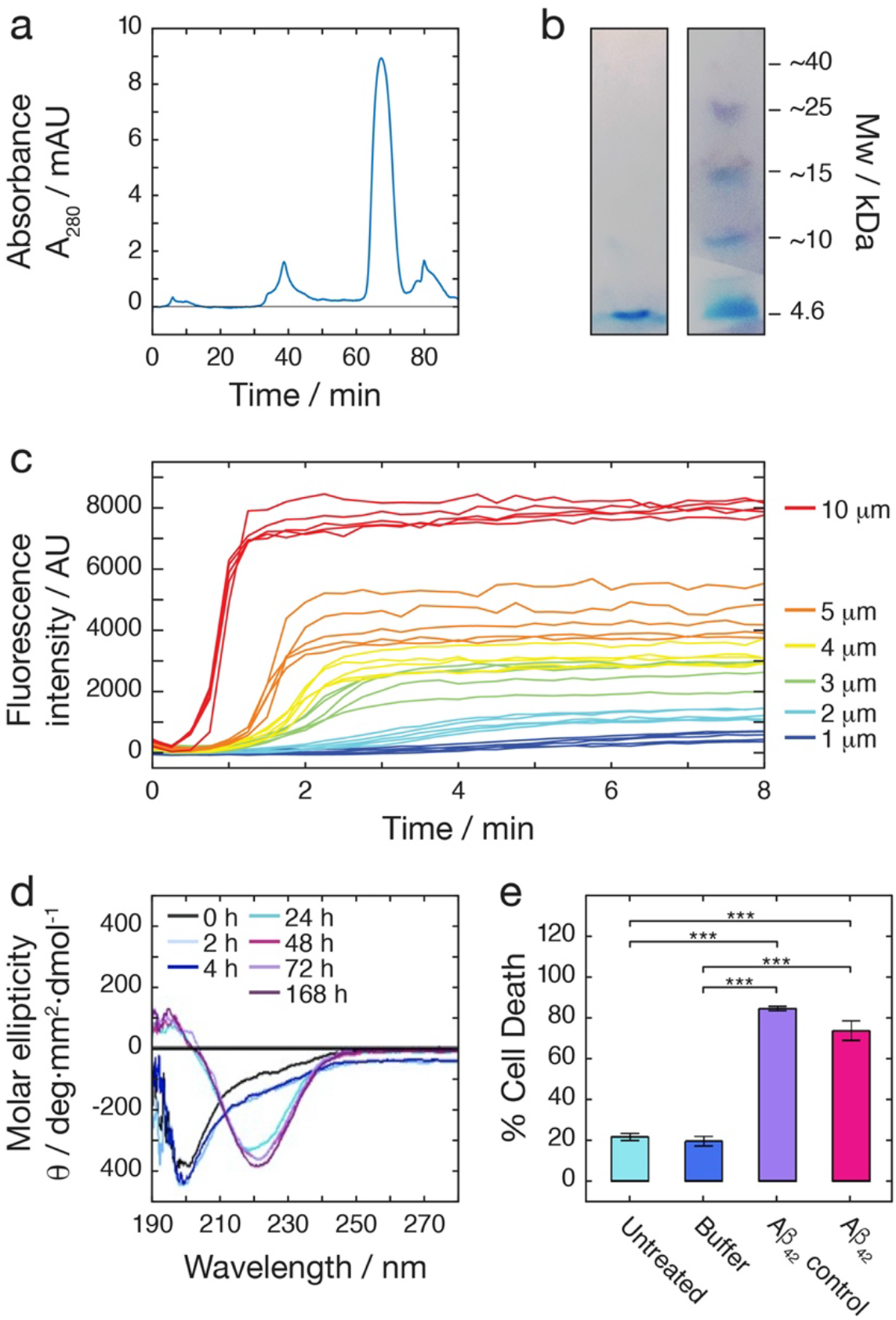
Expression and purification of Aβ_42_. (a) Size exclusion chromatography trace of monomeric recombinant Aβ_42_ elution immediately before use. Aβ_42_ was expressed and purified as described in the Methods section, with the last purification step consisting of multiple rounds of SEC, which were performed until only a single peak appeared on the chromatogram. (b) SDS-PAGE confirming the purified Aβ_42_ monomer sample used. Fibril formation was tracked by ThT assay (c) and by CD (d), demonstrating the expected assembly competency of the monomeric recombinant Aβ_42_ samples. (e) The cytotoxicity of small oligomeric species formed from the monomeric recombinant Aβ_42_ sample compared to species formed using a commercial sample (Aβ_42_ control) from rPeptide ^38^ demonstrate the expected cytotoxic potential for the recombinant Aβ_42_ samples.

**Supplementary Figure S2:**
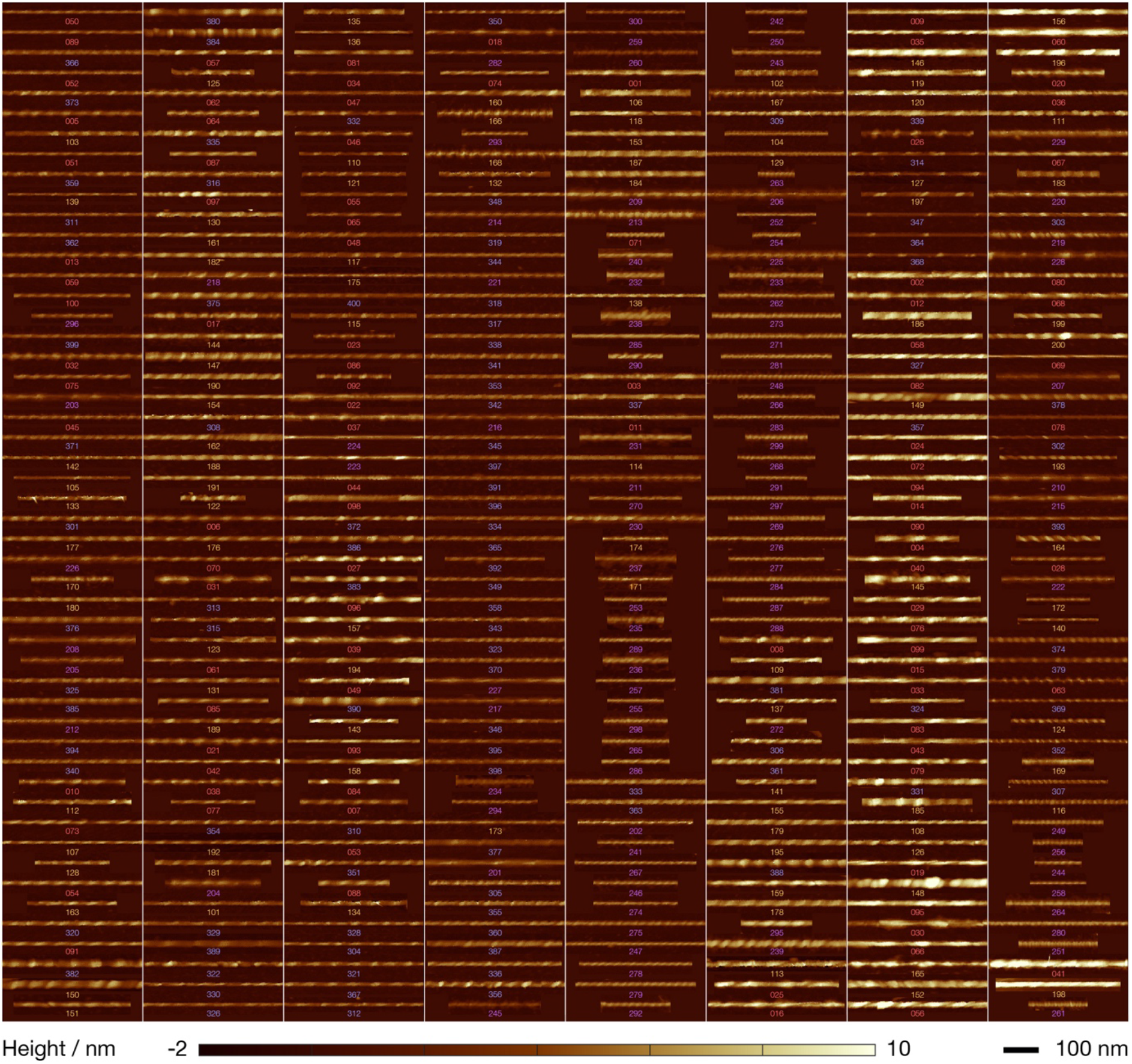
AFM image data for all of the 400 individually analysed Aβ_42_. fibrils. Up to a 400 nm of digitally straightened segment are shown for each individual fibril analysed. The fibrils are shown with their individual index numbers (see **Supplementary Table 2**) coloured coded according to the assembly condition they were found in (20mM of sodium phosphate pH 8: red #1-100, sodium phosphate pH 7.4: orange #101-200, Tris pH 7.4: purple #201-300, and HEPES pH 7.4: blue #301-400). The fibrils are arranged by similarity, in the same order from upper-left column-wise to lower-right as shown from bottom to top of the dendrogram in Figure 4. All fibril images are identically scaled for comparison, with the colour scale of the images represent height range of −2 to 10 nm as indicated by the bottom colour bar and length scale of 100 nm indicated by the scalebar.

**Supplementary Figure S3:**
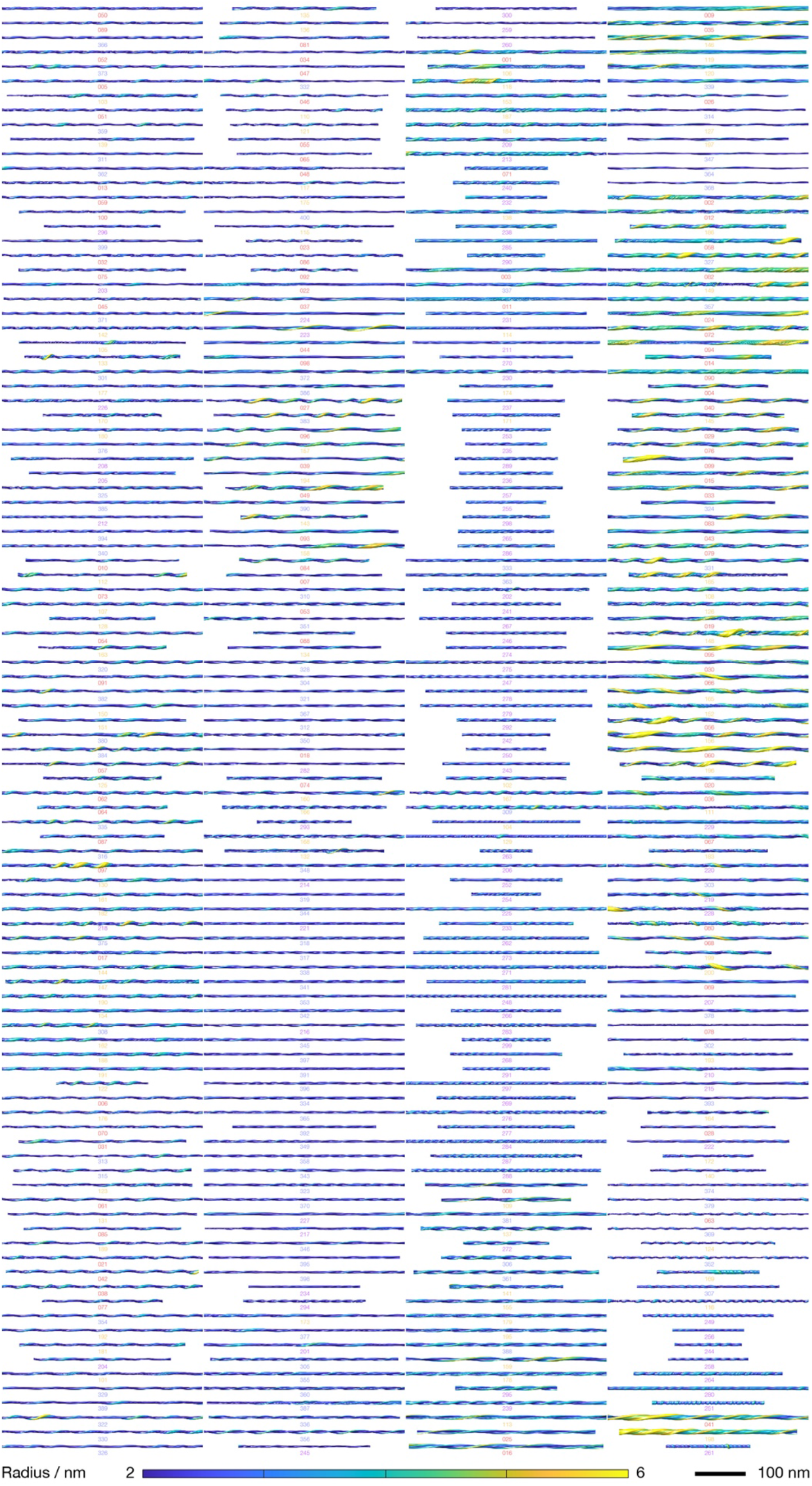
3D models of all of the 400 individually analysed Aβ_42_. fibrils. Up to a 400 nm of reconstructed fibril 3D surface envelopes are displayed for each individual fibril analysed. The fibrils are shown with their individual index numbers (see **Supplementary Table 2**) coloured coded according to the assembly condition they were found in (20mM of sodium phosphate pH 8: red #1-100, sodium phosphate pH 7.4: orange #101-200, Tris pH 7.4: purple #201-300, and HEPES pH 7.4: blue #301-400). The fibrils are arranged by similarity, in the identical order from upper-left column-wise to lower-right as shown in Supplementary Figure S2 and from bottom to top of the dendrogram in Figure 4. All of the models are shown with identical scales for comparison, with the colour represent the local radius to the screw axis according to the bottom colour bar for visualisation and length scale of 100 nm indicated by the scalebar.

**Supplementary Figure S4:**
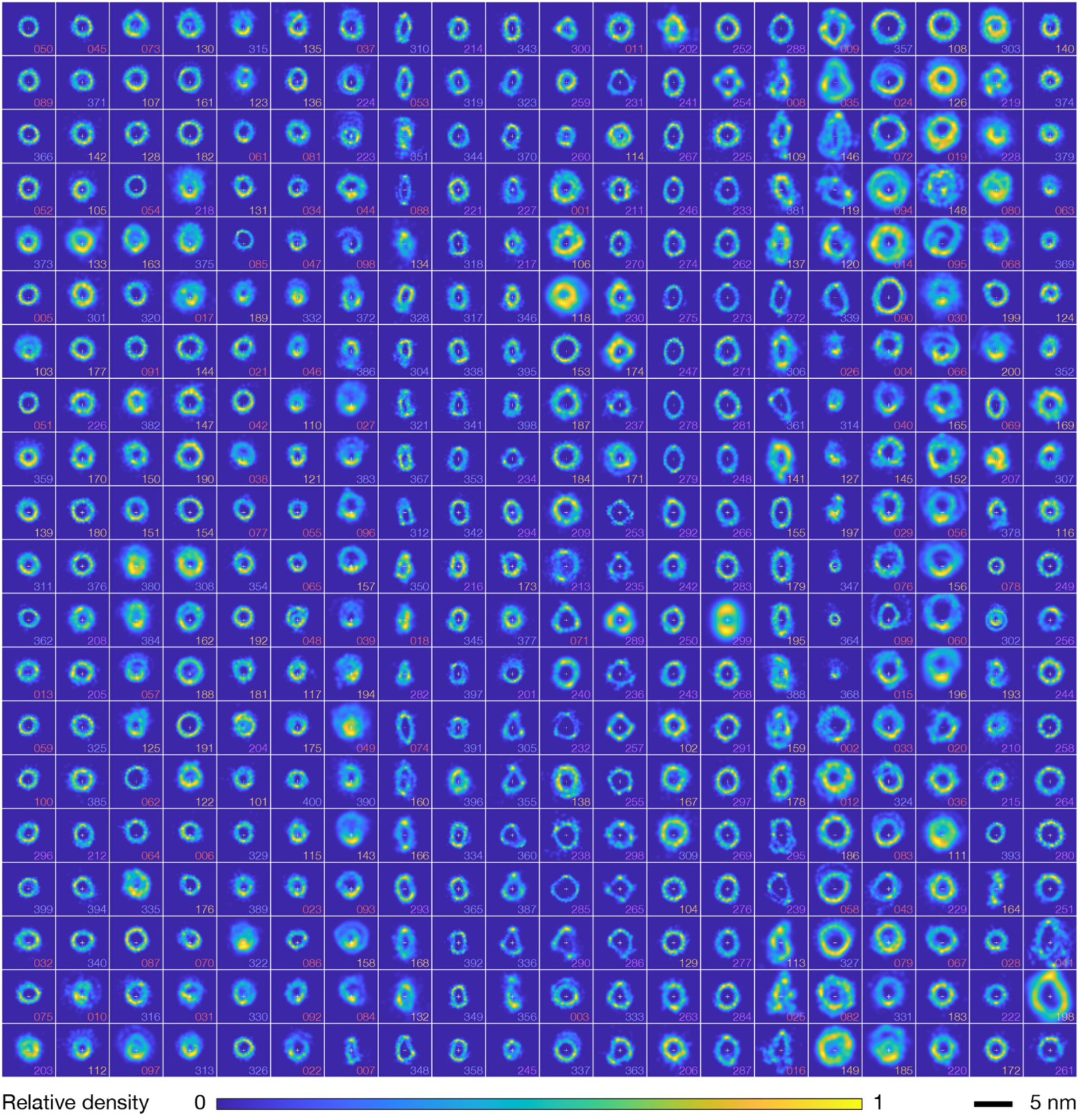
Cross-sectional contact-point density map for all of the 400 individually analysed Aβ_42_. fibrils. The cross-sectional contact-point densities are scaled between 0 to 1 relative to the highest density for each density map. The fibrils are shown with their individual index numbers (see **Supplementary Table 2**) coloured coded according to the assembly condition they were found in (20mM of sodium phosphate pH 8: red #1-100, sodium phosphate pH 7.4: orange #101-200, Tris pH 7.4: purple #201-300, and HEPES pH 7.4: blue #301-400). The fibrils are arranged by similarity, in the identical order from upper-left column-wise to lower-right as shown in Supplementary Figure S2 and S3, and from bottom to top of the dendrogram in Figure 4. All of the density maps are shown with identical scales for comparison, with the colour represent the relative density and length scale of 5 nm indicated by the scalebar.

**Supplementary Table S1:** Solution conditions commonly used for Aβ_42_ assembly. Solution conditions used in highly cited primary Aβ_42_ research publications between 2005 and May 2020. The 20 highly cited publications involving Aβ_42_ from each of these years were tabulated and the buffer conditions in which monomer was incubated in which substantial polymerisation could occur are listed.

| Title | Publication Year | Corresponding Author(s) | Solution Condition |
| --- | --- | --- | --- |
| A hybrid molecule that prohibits amyloid fibrils and alleviates neuronal toxicity induced by β-amyloid (1–42) | 2005 | Jaehoon Yu | 10 mM sodium phosphate at pH 7.4 |
| Certain Inhibitors of Synthetic Amyloid β-Peptide (Aβ) Fibrillogenesis Block Oligomerization of Natural Aβ and Thereby Rescue Long-Term Potentiation | 2005 | Dennis J. Selkoe | 100 mM Tris-HCl, pH 7.4 |
| Protection of rat primary hippocampal cultures from Aβ cytotoxicity by pro-inflammatory molecules is mediated by astrocytes | 2005 | Romy von Bernhardt | 12 mM Tris buffer, pH 7 |
| Amyloid-β Peptide Inhibits Activation of the Nitric Oxide/cGMP/cAMP-Responsive Element-Binding Protein Pathway during Hippocampal Synaptic Plasticity | 2005 | Ottavio Arancio | ACSF |
| Fucoidan inhibits cellular and neurotoxic effects of β-amyloid (Aβ) in rat cholinergic basal forebrain neurons | 2005 | S. Kar | ACSF/ Neurobasal |
| Novel Aβ peptide immunogens modulate plaque pathology and inflammation in a murine model of Alzheimer's disease | 2005 | Charles G. Glabe and Andrea J. Tenner | MilliQ water |
| γ-Glutamylcysteine ethyl ester protection of proteins from Aβ(1–42)-mediated oxidative stress in neuronal cell culture: A proteomics approach | 2005 | D. Allan Butterfield | MilliQ water |
| Modulation of the humoral and cellular immune response in Aβ immunotherapy by the adjuvants monophosphoryl lipid A (MPL), cholera toxin B subunit (CTB) and E. coli enterotoxin LT(R192G) | 2005 | Cynthia A. Lemere | MilliQ water |
| β-Amyloid (Aβ) causes detachment of N1E-115 neuroblastoma cells by acting as a scaffold for cell-associated plasminogen activation | 2005 | Martijn F.B.G. Gebbink | MilliQ water |
| Neurotoxicity and oxidative stress in D1M-substituted Alzheimer's Aβ(1–42): relevance to N-terminal methionine chemistry in small model peptides | 2005 | D. Allan Butterfield | MilliQ water |
| β-Amyloid-Induced Neuronal Apoptosis Involves c-Jun N-Terminal Kinase-Dependent Downregulation of Bcl-w | 2005 | Christian Pike | MilliQ water |
| Microglial Phagocytosis Induced by Fibrillar β-Amyloid and IgGs Are Differentially Regulated by Proinflammatory Cytokines | 2005 | Gary E. Landreth | MilliQ water |
| Proteomic identification of proteins specifically oxidized by intracerebral injection of amyloid β-peptide (1–42) into rat brain: Implications for Alzheimer's disease | 2005 | D. Allan Butterfield | MilliQ water |
| Differential effects of oligomeric and fibrillar amyloid-β1–42 on astrocyte-mediated inflammation | 2005 | Mary Jo LaDu | Oligomers - Ham's F-12 (phenol red-free) / Fibrils - 10mM HCl |
| β-Secretase-Cleaved Amyloid Precursor Protein Accumulates at Actin Inclusions Induced in Neurons by Stress or Amyloid $\beta$: A Feedforward Mechanism for Alzheimer's Disease | 2005 | James R. Branburg | Oligomers - Ham's F-12 (phenol red-free) / Fibrils - 10mM HCl |
| Proteomic identification of proteins oxidized by Aβ(1–42) in synaptosomes: Implications for Alzheimer's disease | 2005 | D. Allan Butterfield | PBS |
| Erythropoietin Requires NF-κB and its Nuclear Translocation to Prevent Early and Late Apoptotic Neuronal Injury During β-Amyloid Toxicity | 2005 | Kenneth Maiese | PBS |
| Oxidation of cholesterol catalyzed by amyloid β-peptide (Aβ)–Cu complex on lipid membrane | 2005 | Ryoichi Kuboi | PBS, pH 7.4 |
| Trehalose differentially inhibits aggregation and neurotoxicity of beta-amyloid 40 and 42 | 2005 | Michael R. Sierks | PBS, pH 7.4 |
| A 13 kDa carboxy-terminal fragment of ApoE stabilizes Abeta hexamers | 2005 | Christian Czech | PBS, pH 7.2 |
| Immunization with amyloid-β using GM-CSF and IL-4 reduces amyloid burden and alters plaque morphology | 2006 | JoAnne McLaurin | PBS, pH 7.5 |
| Antibodies against β-amyloid reduce aβ oligomers, glycogen synthase kinase-3β activation and $\tau$ phosphorylation in vivo and in vitro | 2006 | Greg M. Cole | 10 mM HEPES, pH 7.4 |
| PEGylated phospholipid nanomicelles interact with β-amyloid(1–42) and mitigate its β-sheet formation, aggregation and neurotoxicity in vitro | 2006 | Hayat Önyüksel | 10 mM HEPES |
| Interaction between Aβ Peptide and $\alpha$ Synuclein: Molecular Mechanisms in Overlapping Pathology of Alzheimer's and Parkinson's in Dementia with Lewy Body Disease | 2006 | Ratna Mandal | 100 mM SDS in 90% MilliQ H<sub>2</sub>O and 10% D<sub>2</sub>O at pH 7.2 |
| The cAMP-specific phosphodiesterase 4B mediates Aβ-induced microglial activation | 2006 | Patrick Tremblay | 5 mM Tris, pH 7.4 |
| The various aggregation states of β-amyloid 1–42 mediate different effects on oxidative stress, neurodegeneration, and BACE-1 expression | 2006 | Massimo Tabaton | DMEM |
| Role of toll-like receptor signalling in Aβ uptake and clearance | 2006 | Ken - ichiro Fukuchi | Ham's F-12 (phenol red-free) |
| Autoinsertion of soluble oligomers of Alzheimer's Aβ(1–42) peptide into cholesterol-containing membranes is accompanied by relocation of the sterol towards the bilayer surface | 2006 | Jeremy P. Bradshaw | Ham's F-12 (phenol red-free) |
| Autophagy of amyloid beta-protein in differentiated neuroblastoma cells exposed to oxidative stress | 2006 | Alexei Terman | Ham's F-12 (phenol red-free) |
| Soluble oligomers of amyloid-β peptide induce neuronal apoptosis by activating a cPLA2-dependent sphingomyelinase-ceramide pathway | 2006 | Thierry Pillot | Ham's F-12 (phenol red-free) |
| Alzheimer's amyloid β-peptide (1–42) induces cell death in human neuroblastoma via bax/bcl-2 ratio increase: An intriguing role for methionine 35 | 2006 | F. Misiti | MEM |
| Low molecular weight thiol amides attenuate MAPK activity and protect primary neurons from Aβ(1–42) toxicity | 2006 | Daphne Atlas | MilliQ water |
| β-Amyloid Stimulates Murine Postnatal and Adult Microglia Cultures in a Unique Manner | 2006 | Colin K. Combs | MilliQ water |
| The spirostenol (22R, 25R)-20α-spirost-5-en-3β-yl hexanoate blocks mitochondrial uptake of Aβ in neuronal cells and prevents Aβ-induced impairment of mitochondrial function | 2006 | Vassilios Papaopoulos | MilliQ water, pH 6 |
| Lack of glutathione peroxidase-1 exacerbates Ab-mediated neurotoxicity in cortical neurons | 2006 | R. C. Iannello | Neurobasal |
| Apoptosis is secondary to non-apoptotic axonal degeneration in neurons exposed to Aβ in distal axons | 2006 | Elena I. Posse de Chaves | Oligomers - Ham's F-12 (phenol red-free) / Fibrils - 10mM HCl |
| Active immunization trial in Aβ42-injected P301L tau transgenic mice | 2006 | Jürgen Götz | PBS |
| Immunization with fibrillar Aβ1–42 in young and aged canines: Antibody generation and characteristics, and effects on CSF and brain Aβ | 2006 | Carl W. Cotman | PBS, pH 7.5 |
| Development of new screening system for Alzheimer disease, in vitro Aβ sink assay, to identify the dissociation of soluble Aβ from fibrils | 2006 | Ryuichi Morishita | PBS, pH 7.5 |
| Copper-dependent inhibition of cytochrome c oxidase by Aβ1–42 requires reduced methionine at residue 35 of the Aβ peptide | 2006 | Ian A. Trounce | PBS, pH 8 |
| Aβ peptides can enter the brain through a defective blood–brain barrier and bind selectively to neurons | 2007 | Robert G. Nagele | 0.5 × PBS, 250 mM HEPES buffer, pH 8.5 |
| Single particle detection of Aβ aggregates associated with Alzheimer's disease | 2007 | Dieter Willbold | 140 mM NaCl; 2.7 mM KCl; 10 mM Na<sub>2</sub>HPO<sub>4</sub>, pH 7.4 |
| Requirement of aggregation propensity of Alzheimer amyloid peptides for neuronal cell surface binding | 2007 | Avijit Chakrabartty | 2 mM borate, 2 mM citrate and 2 mM phosphate, pH 6.5 |
| Lipid peroxidation and 4-hydroxy-2-nonenal formation by copper ion bound to amyloid-β peptide | 2007 | Masao Nakamura | 5 mM Hepes buffer, pH 7.4 |
| Neprilysin protects neurons against Aβ peptide toxicity | 2007 | Mark S. Kindy | 50 mM HEPES buffer, pH 7.2 |
| Dopamine release in prefrontal cortex in response to β-amyloid activation of α7* nicotinic receptors | 2007 | Robert A. Nichols | ACSF |
| Aβ Upregulates and Colocalizes with LGI3 in Cultured Rat Astrocytes | 2007 | Yasuhiro Yoshikawa | DMEM |
| Minocycline does not affect amyloid $\beta$ phagocytosis by human microglial cells | 2007 | Robert Veerhuis | DMEM |
| Natural Oligomers of the Alzheimer Amyloid-β Protein Induce Reversible Synapse Loss by Modulating an NMDA-Type Glutamate Receptor-Dependent Signaling Pathway | 2007 | Dennis J. Selkoe and Bernado L. Sabatini | Eluted in 50 mm ammonium acetate, pH 8.5 |
| Rapid, concurrent alterations in pre- and postsynaptic structure induced by naturally-secreted amyloid-β protein | 2007 | Shelley Halpain | Eluted in 50 mm ammonium acetate, pH 8.5 |
| Aβ Oligomer-Induced Aberrations in Synapse Composition, Shape, and Density Provide a Molecular Basis for Loss of Connectivity in Alzheimer's Disease | 2007 | William L. Klein | Ham's F-12 (phenol red-free) |
| Aβ Oligomers Induce Neuronal Oxidative Stress through an N-Methyl-D-aspartate Receptor-dependent Mechanism That Is Blocked by the Alzheimer Drug Memantine | 2007 | Sergio T. Ferreira and William L. Klein | Ham's F-12 (phenol red-free) |
| Lipoprotein Receptor-Related Protein-1 Mediates Amyloid-β-Mediated Cell Death of Cerebrovascular Cells | 2007 | Marcel M. Verbeek | MEM |
| High content screen microscopy analysis of Aβ1–42-induced neurite outgrowth reduction in rat primary cortical neurons: Neuroprotective effects of α7 neuronal nicotinic acetylcholine receptor ligands | 2007 | Jinhe Li | MEM - F12 |
| Lovastatin protects human neurons against Aβ-induced toxicity and causes activation of β-catenin–TCF/LEF signaling | 2007 | Francis Amara | MilliQ water |
| Dopamine and Aβ-Induced Stress Signaling and Decrements in Ca<sup>2+</sup> Buffering In Primary Neonatal Hippocampal Cells Are Antagonized by Blueberry Extract | 2007 | D. R. Fisher | MilliQ water |
| Soluble amyloid beta1-42 reduces dopamine levels in rat prefrontal cortex: Relationship to nitric oxide | 2007 | S. Govoni | MilliQ water |
| Aβ42 neurotoxicity in primary co-cultures: Effect of apoE isoform and Aβ conformation | 2007 | Mary Jo LaDu | Oligomers - Ham's F-12 (phenol red-free) / Fibrils - 10mM HCl |
| Oligomeric Amyloid Decreases Basal Levels of Brain-Derived Neurotrophic factor (BDNF) mRNA via Specific Downregulation of BDNF Transcripts IV and V in Differentiated Human Neuroblastoma Cells | 2007 | Margaret Fahnestock | Oligomers - Ham's F-12 (phenol red-free) / Fibrils - 10mM HCl |
| Interface peptide of Alzheimer's amyloid beta: Application in purification | 2007 | Alagiri Srinivasan | PBS |
| TGF-β1 protects against Aβ-neurotoxicity via the phosphatidylinositol-3-kinase pathway | 2008 | Agata Copani | 0.1 M phosphate-buffered saline |
| Amyloid-β Peptide (Aβ) Neurotoxicity Is Modulated by the Rate of Peptide Aggregation: Aβ Dimers and Trimers Correlate with Neurotoxicity | 2008 | Kevin J. Barnham | 10 mm Na<sub>2</sub>HPO<sub>4</sub> and NaH<sub>2</sub>PO<sub>4</sub>, pH 7.4 |
| Aβ Mediated Diminution of MTT Reduction—An Artefact of Single Cell Culture? | 2008 | Marcus Fändrich and Klaus G. Reymann | 20 mM NaH<sub>2</sub>PO<sub>4</sub>, 140 mM NaCl, 0.2% SDS, pH 7.4 |
| Lipids revert inert Aβ amyloid fibrils to neurotoxic protofibrils that affect learning in mic | 2008 | Frederic Rousseau | 50 mM Tris, pH 7.5 |
| Salvianolic acid B inhibits Aβ fibril formation and disaggregates preformed fibrils and protects against Aβ-induced cytotoxicity | 2008 | William L. Klein and Min Li | 50 mM phosphate buffer pH7.4, and 100 mM NaCl |
| Aβ inhibits the proteasome and enhances amyloid and tau accumulation | 2008 | Frank M. LaFerla | 50 mM Tris, pH 7.5, 25 mM KCl, 10 mM NaCl, 1 mM MgCl<sub>2</sub>, 0.03% SDS |
| Differences between normal and alpha-synuclein overexpressing SH-SY5Y neuroblastoma cells after Aβ(1-42) and NAC treatment | 2008 | Zsolt L. Datki | DMEM |
| Aβ Oligomers Induce Neuronal Cell Cycle Events in Alzheimer's Disease | 2008 | Bruce T. Lamb | Ham's F-12 (phenol red-free) |
| ApoE Promotes the Proteolytic Degradation of Aβ | 2008 | Gary E. Landreth | MilliQ water |
| Blocking Aβ42 Accumulation Delays the Onset and Progression of Tau Pathology via the C Terminus of Heat Shock Protein70-Interacting Protein: A Mechanistic Link between Aβ and Tau Pathology | 2008 | Frank M. LaFerla | MilliQ water |
| Aβ(1-42) injection causes memory impairment, lowered cortical and serum BDNF levels, and decreased hippocampal 5-HT<sub>2A</sub> levels | 2008 | S. Aznar | MilliQ water |
| Aggregation of Aβ(1-42) in the presence of short peptides: conformational studies | 2008 | Botond Penke | MilliQ water (pH 6) |
| Small Molecule, Non-Peptide p75<sup>NTR</sup> Ligands Inhibit Aβ-Induced Neurodegeneration and Synaptic Impairment | 2008 | Frank M. Longo | Oligomers - 0.5 mM NaOH or 0.2% NH<sub>4</sub>OH or PBS / Fibrils - 10mM HCl |
| Oligomer-specific Aβ toxicity in cell models is mediated by selective uptake | 2008 | Wiep Scheper | Oligomers - DMEM (phenol red - free)/ Fibrils - 10mM HCl |
| E2-25K/Hip-2 regulates caspase-12 in ER stress-mediated Aβ neurotoxicity | 2008 | Yong - Keun Jung | PBS |
| Alzheimer's disease-type neuronal tau hyperphosphorylation induced by Aβ oligomers | 2008 | Fernanda G De Felice | PBS, pH 7.4 |
| Grape-Derived Polyphenolics Prevent Aβ Oligomerization and Attenuate Cognitive Deterioration in a Mouse Model of Alzheimer's Disease | 2008 | Giulio M. Pasinetti | PBS, pH 7.4 |
| Anti-oligomeric Aβ Single-chain Variable Domain Antibody Blocks Aβ-induced Toxicity Against Human Neuroblastoma Cells | 2008 | Michael R. Sierks | PBS, pH 7.4 |
| Aβ-globulomers are formed independently of the fibril pathway | 2008 | Stefan Barghorn | PBS, pH 7.4 |
| A Two-Year Study with Fibrillar β-Amyloid (Aβ) Immunization in Aged Canines: Effects on Cognitive Function and Brain Aβ | 2008 | David Cribbs | PBS, pH 7.5 |
| GluN2B subunit-containing NMDA receptor antagonists prevent Aβ-mediated synaptic plasticity disruption in vivo | 2009 | Michael J. Rowan | 0.9% saline |
| Design and synthesis of newtrehalose-conjugated pentapeptides as inhibitors of Aβ(1-42) fibrillogenesis and toxicity | 2009 | Enrico Rizzarelli | 20 mM PBS or 25 mM NEM buffer, pH 7.4 |
| Comparison of Alzheimer Aβ(1-40) and Aβ(1-42) amyloid fibrils reveals similar protofilament structures | 2009 | Marcus Fändrich and Nikolaus Grigorieff | 50 mM Tris/HCl (pH 7.4) |
| Aβ-Dependent Inhibition of LTP in Different Intracortical Circuits of the Visual Cortex: The Role of RAGE | 2009 | Luciano Domenici | ACSF |
| Preparation and Characterization of a Monoclonal Antibody with High Affinity for Soluble Ab Oligomers | 2009 | Hong Tao | Ham's F-12 (phenol red-free) |
| Protection of synapses against Alzheimer's-linked toxins: Insulin signaling prevents the pathogenic binding of Aβ oligomers | 2009 | William L. Klein | Ham's F-12 (phenol red-free) |
| The binding of pullulan modified cholesteryl nanogels to Aβ oligomers and their suppression of cytotoxicity | 2009 | Dusica Maysinger | Ham's F-12 (phenol red-free) |
| CD14 and Toll-Like Receptors 2 and 4 Are Required for Fibrillar Aβ-Stimulated Microglial Activation | 2009 | Gary E. Landreth | MilliQ water |
| Autophagy protects neuron from Aβ-induced cytotoxicity | 2009 | Wen-Mei Fu | MilliQ water |
| TrkA pathway activation induced by amyloid-beta (Aβ) | 2009 | Massimo Masserini | Neurobasal, pH 7.4 |
| Pin1 affects Tau phosphorylation in response to Aβ oligomers | 2009 | Massimo Masserini | Neurobasal, pH 7.4 |
| Enlargement of Aβ aggregates through chemokine-dependent microglial clustering | 2009 | Huey - Jen Tsay | Oligomers - Ham's F-12 (phenol red-free) / Fibrils - 10mM HCl and 150mM NaCl |
| PP1 inhibition by Aβ peptide as a potential pathological mechanism in Alzheimer's disease | 2009 | Edgar F. da Cruz e Silva | Oligomers - possibly cell culture medium / Fibrils - 10mM HCl |
| Aβ peptide conformation determines uptake and interleukin-1α expression by primary microglial cells | 2009 | Greer M. Murphy Jr. | Oligomers and Fibrils - PBS / Protofibrils - Ham's F12 (all pH 7.4) |
| Human apolipoprotein A-I binds amyloid-β and prevents Aβ-induced neurotoxicity | 2009 | Sergio T. Ferreira | PBS |
| Single-domain antibodies recognize selectively small oligomeric forms of amyloid $\beta$, prevent Aβ-induced neurotoxicity and inhibit fibril formation | 2009 | François Rougeon | PBS |
| Diazoxide Reverses the Enhanced Expression of KATP Subunits in Cholinergic Neurons Caused by Exposure to Aβ1-42 | 2009 | Kejing Liu | PBS |
| A Role for Synaptic Zinc in Activity-Dependent Aβ Oligomer Formation and Accumulation at Excitatory Synapses | 2009 | Jorge Busciglio | PBS, pH 7.4 |
| Characterisation of two antibodies to oligomeric Aβ and their use in ELISAs on human brain tissue homogenates | 2009 | Seth Love | PBS; 0.15 M sodium chloride, 1.9 mM sodium dihydrogen orthophosphate 1-hydrate, 7.5 mM di-sodium hydrogen orthophosphate 12-hydrate, pH 7.1) or Dulbecco's Modified Eagle Medium |
| The PI3K-Akt-mTOR pathway regulates Aβ oligomer induced neuronal cell cycle events | 2009 | Bruce T. Lamb | Sodium phosphate buffer at pH 7.4 |
| Hydralazine modifies Aβ fibril formation and prevents modification by lipids in vitro | 2010 | Ian V. J. Murray | 100mM PBS, pH 7.4 |
| Aβ20–29 peptide blocking apoE/Aβ interaction reduces full-length Aβ42/40 fibril formation and cytotoxicity in vitro | 2010 | Zhuyi Li | 100mM Tris, pH 7.4 |
| TGF-β1 blockade of microglial chemotaxis toward Aβ aggregates involves SMAD signaling and down-regulation of CCL5 | 2010 | Huey - Jen Tsay | 10mM HCl and 150mM NaCl |
| Stability of Aβ (1-42) peptide fibrils as consequence of environmental modifications | 2010 | Francesco Mantegazza | 10mM HCl and switched to various buffers at pH 7.4 or 10 mM Tris HCl, 150 mM NaCl, pH 7.4 |
| Retardation of Aβ Fibril Formation by Phospholipid Vesicles Depends on Membrane Phase Behavior | 2010 | Sara Linse | 20 mM Tris/HCl, pH 7.4, 0.2 mM EDTA, 0.02% NaN<sub>3</sub> |
| Surface Plasmon Resonance Spectroscopy in Determination of the Interactions Between Amyloid $\beta$ Proteins (Aβ) and Lipid Membranes | 2010 | Marie - Isabel Aguilar | 20 mM sodium phosphate (pH 6.8) |
| The Native Copper- and Zinc- Binding Protein Metallothionein Blocks Copper-Mediated Aβ Aggregation and Toxicity in Rat Cortical Neurons | 2010 | Adrian K. West | 20mM Tris-HCl, 100mM NaCl, pH 7.4 |
| Sensitive detection of Aβ protofibrils by proximity ligation - relevance for Alzheimer's disease | 2010 | Ulf Landegren | 50 mM phosphate buffer and 100 mM NaCl, pH 7.4 |
| Preparation and characterization of toxic Aβ aggregates for structural and functional studies in Alzheimer's disease research | 2010 | Hilal A. Lashuel | 500 mM glycine-NaOH, pH 8.5 |
| Amyloid beta from axons and dendrites reduces local spine number and plasticity | 2010 | Roberto Malinow | ACSF |
| Aβ Oligomers Cause Localized Ca<sup>2+</sup> Elevation, Missorting of Endogenous Tau into Dendrites, Tau Phosphorylation, and Destruction of Microtubules and Spines | 2010 | Eckhard and Eva Mandelkow | Ham's F-12 (phenol red-free) |
| Soluble oligomeric forms of beta-amyloid (Aβ) peptide stimulate Aβ production via astrogliosis in the rat brain | 2010 | A. Gonzalo - Ruiz | Ham's F-12 (phenol red-free) |
| Activation of PERK Signaling Attenuates Aβ-Mediated ER Stress | 2010 | Sung Su Kim | MilliQ water |
| Human intravenous immunoglobulin provides protection against Aβ toxicity by multiple mechanisms in a mouse model of Alzheimer's disease | 2010 | Milla Koistinaho | MilliQ water |
| Conformation dependent monoclonal antibodies distinguish different replicating strains or conformers of prefibrillar Aβ oligomers | 2010 | Charles Glabe | MilliQ water/ HEPES buffered saline, pH 7.4 (HBS) / phosphate buffered saline pH 7.4 (PBS) all supplemented with 0.02% NaN<sub>3</sub> |
| Synthetic amyloid-β oligomers impair long-term memory independently of cellular prion protein | 2010 | Gianluigi Forloni | Oligomers - 50 mM phosphate buffer, 150 mM NaCl, pH 7.4 / Fibrils - 10mM HCl |
| Selenium attenuates Aβ production and Aβ-induced neuronal death | 2010 | Dong - Gyu Jo | PBS |
| Stabilization of neurotoxic Alzheimer amyloid-β oligomers by protein engineering | 2010 | Christopher Dobson and Torleif Hard | PBS, pH 7.2 |
| Linear and conformation specific antibodies in aged beagles after prolonged vaccination with aggregated Abeta | 2010 | Elizabeth Head | PBS, pH 7.5 |
| The presence of sodium dodecyl sulphate-stable Aβ dimers is strongly associated with Alzheimer-type dementia | 2010 | Dominic Walsh | Tris-buffered saline (TBS), containing 5 mM ethylenediaminetetraacetic acid (EDTA), 5 mM ethylene glycol tetraacetic acid (EGTA), 10 mg/ml leupeptin, 1 mg/ml aprotinin, 1 mg/ml pepstatin A, 1 mM Pefabloc, 2 mM 1,10-phenanthroline |
| The Osaka FAD Mutation E22Δ Leads to the Formation of a Previously Unknown Type of Amyloid $\beta$ Fibrils and Modulates Aβ Neurotoxicity | 2011 | Rudi Glockshuber | 10 mM H<sub>3</sub>PO<sub>4</sub>-NaOH, pH 7.4, 100 mM NaCl |
| Critical Role of Astroglial Apolipoprotein E and Liver X Receptor-α Expression for Microglial Aβ Phagocytosis | 2011 | Michael T. Heneka | 50 mM Tris-HCl, pH 7 |
| Resveratrol Acts Not through Anti-Aggregative Pathways but Mainly via Its Scavenging Properties against Aβ and Aβ-Metal Complexes Toxicity | 2011 | Paolo Zatta | Dialyzed against various metal solutions 10 mM ([CH<sub>3</sub>CH(OH)COO]<sub>3</sub>Al, FeCl<sub>3</sub>, CuCl<sub>2</sub>, ZnCl<sub>2</sub>) and then bidialyzed x3 in MilliQ water, pH 7. *In vitro* preps were diluted into 0.1 M Tris/HCl pH 7.4 buffer plus 0.15 M NaCl |
| Abeta Peptide Toxicity is Reduced After Treatments Decreasing Phosphatidylethanolamine Content in Differentiated Neuroblastoma Cells | 2011 | Massimo Masserini | DMEM/ PBS |
| Mechanism mediating oligomeric Aβ clearance by naïve primary microglia | 2011 | Huey - Jen Tsay | Ham's F-12 (phenol red-free) |
| The Second-Generation Active Aβ Immunotherapy CAD106 Reduces Amyloid Accumulation in APP Transgenic Mice While Minimizing Potential Side Effects | 2011 | Matthias Staufenbiel | Ham's F-12 (phenol red-free) |
| Cognitive effects of cell-derived and synthetically derived Aβ oligomers | 2011 | James P. Cleary | Ham's F-12 (phenol red-free) |
| Aβ Inhibition of Ionic Conductance in Mouse Basal Forebrain Neurons Is Dependent upon the Cellular Prion Protein PrPC | 2011 | Jack H. Jhamandas | Ham's F-12 (phenol red-free) |
| Membrane cholesterol enrichment prevents Aβ-induced oxidative stress in Alzheimer's fibroblasts | 2011 | Cristina Cecchi | Ham's F-12 (phenol red-free) |
| Specificity and sensitivity of the Abeta oligomer ELISA | 2011 | David A. Loeffler | LMW - 100mM Tris, pH 8.8<br>Oligomers - PBS; 0.01 M, pH 7.4, with 0.02% azide |
| Protection against Aβ-mediated rapid disruption of synaptic plasticity and memory by memantine | 2011 | Michael J. Rowan | MilliQ water |
| p75NTR Regulates Aβ Deposition by Increasing Aβ Production But Inhibiting Aβ Aggregation with Its Extracellular Domain | 2011 | Xin - Fu Zhou | Oligomers - DMEM Fibrils - DMEM with 10mM HCl |
| WNT5A Signaling Contributes to Aβ-Induced Neuroinflammation and Neurotoxicity | 2011 | Shao - Jun Tang | Oligomers - DMEM/F-12 Fibrils - 10mM HCl |
| The contribution of activated astrocytes to Aβ production: Implications for Alzheimer's disease pathogenesis | 2011 | Robert Vassar | Oligomers - Ham's F-12 (phenol red-free), Fibrils -10mM HCl |
| Downregulation of CREB expression in Alzheimer's brain and in Aβ-treated rat hippocampal neurons | 2011 | Christopher B. Eckman | Oligomers - Ham's F-12 (phenol red-free), Fibrils -10mM HCl |
| Functional Links Between Aβ Toxicity, Endocytic Trafficking, and Alzheimer's Disease Risk Factors in Yeast | 2011 | Susan Lindquist | Oligomers - MilliQ water Fibrils - 10mM HCL 6% DMSO |
| Lipid matrix plays a role in Abeta fibril kinetics and morphology | 2011 | Frances Separovic | PBS |
| Sinomenine inhibits microglial activation by Aβ and confers neuroprotection | 2011 | Shiv Kumar Sharma | PBS |
| Soluble Aβ Seeds Are Potent Inducers of Cerebral β-Amyloid Deposition | 2011 | Mathias Jucker | PBS, pH 7 |
| Silibinin: A novel inhibitor of Aβ aggregation | 2011 | Junzeng Zhang | PBS, pH 7.4 |
| ERK and p38 inhibitors attenuate memory deficits and increase CREB phosphorylation and PGC-1α levels in Aβ-injected rats | 2012 | Fariba Khodagholi | 0.1 M PBS |
| Aβ neurotoxicity depends on interactions between copper ions, prion protein, and N-methyl-d-aspartate receptors | 2012 | Gerald W. Zamponi | 0.9 % saline, pH 7.4 |
| The phosphodiesterase-4 inhibitor rolipram reverses Aβ-induced cognitive impairment and neuroinflammatory and apoptotic responses in rats | 2012 | Han - Ting Zhang | 0.9% saline |
| Amyloid-Beta (Aβ) D7H Mutation Increases Oligomeric Aβ42 and Alters Properties of Aβ-Zinc/Copper Assemblies | 2012 | Irene H. Cheng | 10 mM Tris-HCl, pH 7.4 |
| The binding of resveratrol to monomer and fibril amyloid beta | 2012 | Jiang - Ning Zhou | 50 mM phosphate buffer, pH 7.5, and 100 mM NaCl |
| Specific Binding of Alzheimer's Aβ Peptide Fibrils to Single-Walled Carbon Nanotubes | 2012 | Saeyoung Nate Ahn | 50mM MOPS, pH 7.4 |
| Aβ delays fibrin clot lysis by altering fibrin structure and attenuating plasminogen binding to fibrin | 2012 | Sidney Strickland | 50mM Tris pH 7.4, 0.1% NH<sub>4</sub>OH |
| p75NTR is mainly responsible for Aβ toxicity but not for its internalization: a primary study | 2012 | Xin - Fu Zhou | DMEM |
| Synapse-Binding Subpopulations of Aβ Oligomers Sensitive to Peptide Assembly Blockers and scFv Antibodies | 2012 | William L. Klein | Ham's F-12 (phenol red-free) |
| Aβ Oligomers-Induced Toxicity is Attenuated in Cells Cultured with NbActiv4™ Medium | 2012 | William L. Klein | Ham's F-12 (phenol red-free) |
| Redox-active Cu(II)-Aβ causes substantial changes in axonal integrity in cultured cortical neurons in an oxidative-stress dependent manner | 2012 | Roger S. Chung | MilliQ water |
| Aβ-induced formation of autophagosomes is mediated by RAGE-CaMKKβ-AMPK signaling | 2012 | Inhee Mook-Jung | Opti-MEM |
| An Effector-Reduced Anti-β-Amyloid (Aβ) Antibody with Unique Aβ Binding Properties Promotes Neuroprotection and Glial Engulfment of Aβ | 2012 | Andreas Muhs and Ryan J. Watts | PBS |
| NMDA receptors and BAX are essential for Aβ impairment of LTP | 2012 | Morgan Sheng | PBS |
| A critical role for the PAR-1/MARK-tau axis in mediating the toxic effects of Aβ on synapses and dendritic spines | 2012 | Bingwei Li | PBS |
| Neuropathogenic role of adenylate kinase-1 in Aβ-mediated tau phosphorylation via AMPK and GSK3β | 2012 | Yong - Keun Jung | PBS |
| The Microtubule-Associated Protein 1A (MAP1A) is an Early Molecular Target of Soluble Aβ-Peptide | 2012 | A. B. Klein | PBS containing 0.1% NH<sub>4</sub>OH |
| A Chemical Analog of Curcumin as an Improved Inhibitor of Amyloid Abeta Oligomerization | 2012 | David L. Vander Jagt | PBS, pH 7.2 |
| An anti-diabetes agent protects the mouse brain from defective insulin signaling caused by Alzheimer's disease-associated Aβ oligomers | 2012 | Fernanda G De Felice | PBS, pH 7.4 |
| Interaction between NH<sub>2</sub>-tau fragment and Aβ in Alzheimer's disease mitochondria contributes to the synaptic deterioration | 2012 | Pietro Calissano | PBS, pH 7.4 |
| Inhibition of Phosphorylation of JNK Suppresses Aβ-Induced ER Stress and Upregulates Prosurvival Mitochondrial Proteins in Rat Hippocampus | 2013 | Fatemeh Shaerzadeh | 0.1 M PBS |
| Aβ Association Inhibition by Transferrin | 2013 | Giuseppe Melacini | 15 mM potassium phosphate buffer at pH 7.4, with 10 % D<sub>2</sub>O and 0.02 % NaN<sub>3</sub> |
| Proteolytically Inactive Insulin-Degrading Enzyme Inhibits Amyloid Formation Yielding Non-Neurotoxic Aβ Peptide Aggregates | 2013 | Eduardo M. Castaño | 50 mM NaCl, 20 mM Tris-HCl, pH 7.4 |
| Ammonium hydroxide treatment of Aβ produces an aggregate free solution suitable for biophysical and cell culture characterization | 2013 | Blaine R. Roberts | 60mM NaOH - further diluted into: PBS, pH 7.4 - ThT / 50mM Tris, 10mM EDTA, pH 8 - DLS |
| Intracellular accumulation of aggregated pyroglutamate amyloid beta: convergence of aging and Aβ pathology at the lysosome | 2013 | Wiep Scheper | DMEM (phenol red - free) |
| Exosomes neutralize synaptic-plasticity-disrupting activity of Aβ assemblies in vivo | 2013 | Joung - Hun Kim | Ham's F-12 (phenol red-free) |
| Disruption of neocortical histone H3 homeostasis by soluble Aβ: implications for Alzheimer's disease | 2013 | Caterina M. Hernandez | Ham's F-12 (phenol red-free) |
| The CAMKK2-AMPK Kinase Pathway Mediates the Synaptotoxic Effects of Aβ Oligomers through Tau Phosphorylation | 2013 | Franck Polleux | Ham's F-12 (phenol red-free) |
| Blocking the Interaction between Apolipoprotein E and Aβ Reduces Intraneuronal Accumulation of Aβ and Inhibits Synaptic Degeneration | 2013 | Martin J. Sadowski | Ham's F-12 (phenol red-free) |
| Protection against the synaptic targeting and toxicity of Alzheimer's-associated Aβ oligomers by insulin mimetic chiro-inositols | 2013 | William L. Klein | Ham's F-12 (phenol red-free) |
| Leptin and ghrelin prevent hippocampal dysfunction induced by Aβ oligomers | 2013 | C.M.F. Pereira | Ham's F-12 (phenol red-free) |
| Aβ leads to Ca<sup>2+</sup> signaling alterations and transcriptional changes in glial cells | 2013 | Dmitry Lim | Ham's F-12 (phenol red-free) |
| Canonical Wnt signaling protects hippocampal neurons from Aβ oligomers: role of non-canonical Wnt-5a/Ca<sup>2+</sup> in mitochondrial dynamics | 2013 | Nibaldo C Inestrosa | MilliQ water |
| Oligomeric Aβ-Induced Microglial Activation is Possibly Mediated by NADPH Oxidase | 2013 | Chun Fu Wu | MilliQ water |
| The cellular prion protein traps Alzheimer's Aβ in an oligomeric form and disassembles amyloid fibers | 2013 | John H. Viles | Monomer - Water, pH 10.5 - maintained using NaOH<br>Fibril elongation - 160mM NaCl, 30mM HEPES, pH 7.4 |
| Memantine Rescues Transient Cognitive Impairment Caused by High-Molecular-Weight Aβ Oligomers But Not the Persistent Impairment Induced by Low-Molecular-Weight Oligomers | 2013 | Sergio T. Ferreira | PBS |
| Immobilized amyloid Aβ peptides support platelet adhesion and activation | 2013 | Mauro Torti | PBS |
| Characterization of a Single-Chain Variable Fragment Recognizing a Linear Epitope of Aβ: A Biotechnical Tool for Studies on Alzheimer's Disease? | 2013 | Dieter Willbold | PBS |
| Peripherally Applied Synthetic Peptide isoAsp7-Aβ(1-42) Triggers Cerebral β-Amyloidosis | 2013 | A.A. Makarov | PBS, pH 7, further diluted in surgical saline |
| Curcumin-conjugated nanoliposomes with high affinity for Aβ deposits: Possible applications to Alzheimer disease | 2013 | Charles Duyckaerts | PBS, with or without 10% Curcumin-conjugated nanoliposomes (CnLs), (20 µg/ml final concentration) |
| Schisantherin A recovers Aβ-induced neurodegeneration with cognitive decline in mice | 2014 | Ying Jia | 0.9% saline |
| Aβ induces its own prion protein N-terminal fragment (PrPN1)-mediated neutralization in amorphous aggregates | 2014 | Xavier Roucou | 10 mM Na<sub>2</sub>HPO<sub>4</sub>, pH 7.4 |
| Distinct synthetic Aβ prion strains producing different amyloid deposits in bigenic mice | 2014 | Stanley B. Prusiner | 10 mM sodium phosphate (NaP) buffer at neutral pH 7.4 with or without 3.47 mM [0.1% (wt/vol)] SDS (Comparison) |
| Immobilization of Homogeneous Monomeric, Oligomeric and Fibrillar Aβ Species for Reliable SPR Measurements | 2014 | Dieter Willbold | 10 mM sodium phosphate buffer pH 7.4 |
| Aβ-AGE aggravates cognitive deficit in rats via RAGE pathway | 2014 | S. Zhang | 10mM PBS, pH 7.2 |
| Self-Propagative Replication of Aβ Oligomers Suggests Potential Transmissibility in Alzheimer Disease | 2014 | Vijayaraghavan Rangachari | 20 mM Tris-HCl, pH 8.0 |
| Soluble Aβ Oligomers Are Rapidly Sequestered from Brain ISF In Vivo and Bind GM1 Ganglioside on Cellular | 2014 | Dennis J. Selkoe | 50 mM ammonium acetate, pH 8.5 |
| Interaction between Prion Protein and Aβ Amyloid Fibrils Revisited | 2014 | Witold K. Surewicz | 7.5mM PBS, pH 7.4 |
| Aβ promotes VDAC1 channel dephosphorylation in neuronal lipid rafts. Relevance to the mechanisms of neurotoxicity in Alzheimer's disease | 2014 | R. Marin | DMEM |
| A Sensitive Aβ Oligomer Assay Discriminates Alzheimer's and Aged Control Cerebrospinal Fluid | 2014 | Alexander McCampbell | Ham's F-12 (phenol red-free) |
| Enhancing Astrocytic Lysosome Biogenesis Facilitates Aβ Clearance and Attenuates Amyloid Plaque Pathogenesis | 2014 | Jin-Moo Lee | Ham's F-12 (phenol red-free) |
| Elucidating Molecular Mass and Shape of a Neurotoxic Aβ Oligomer | 2014 | William L. Klein | Ham's F-12 (phenol red-free) |
| Neurotrophin receptor p75 mediates the uptake of the amyloid beta (Aβ) peptide, guiding it to lysosomes for degradation in basal forebrain cholinergic neurons | 2014 | J. Oliver Dolly | Ham's F-12 (phenol red-free) |
| PKCε Promotes HuD-Mediated Neprilysin mRNA Stability and Enhances Neprilysin-Induced Aβ Degradation in Brain Neurons | 2014 | Daniel Alkon | MilliQ water |
| Interactions between Aβ oligomers and presynaptic cholinergic signaling: Age-dependent effects on attentional capacities | 2014 | Britney Yegla | MilliQ water followed by 1 mM NaHCO<sub>3</sub>, pH 10 after aggregation step |
| Cyclophilin D deficiency rescues Aβ-impaired PKA/CREB signaling and alleviates synaptic degeneration | 2014 | Shirley ShiDu Yan | PBS |
| Intracellular Accumulation of Amyloid-β (Aβ) Protein Plays a Major Role in Aβ-Induced Alterations of Glutamatergic Synaptic Transmission and Plasticity | 2014 | Claudio Grassi | PBS |
| Aβ and tau toxicities in Alzheimer's are linked via oxidative stress-induced p38 activation: Protective role of vitamin E | 2014 | J.Viña | PBS |
| Clioquinol promotes the degradation of metal-dependent amyloid-β (Aβ) oligomers to restore endocytosis and ameliorate Aβ toxicity | 2014 | Susan Lindquist | PBS, pH 7.2 |
| Liposomes bi-functionalized with phosphatidic acid and an ApoE-derived peptide affect Aβ aggregation features and cross the blood-brain-barrier: Implications for therapy of Alzheimer disease | 2014 | Francesca Re | PBS, pH 2 |
| Single Chain Variable Fragment Against Aβ Expressed in Baculovirus Inhibits Abeta Fibril Elongation and Promotes its Disaggregation | 2015 | Tao Hung | 0.1 M boric acid, 2.5 mM NaCl, 2.5 mM sodium borate, pH 8.5 |
| LC3 overexpression reduces Aβ neurotoxicity through increasing α7nAChR expression and autophagic activity in neurons and mice | 2015 | Wen-Mei Fu | 10 mM sodium phosphate buffer |
| QIAD assay for quantitating a compound's efficacy in elimination of toxic Aβ oligomers | 2015 | Dieter Willbold | 10 mM sodium phosphate buffer, pH 7.4 |
| SDS-PAGE analysis of Aβ oligomers is diserving research into Alzheimer's disease: appealing for ESI-IM-MS | 2015 | Natàlia Carulla | 10 mM sodium phosphate, pH 7.4 |
| MicroRNAs 99b-5p/100-5p Regulated by Endoplasmic Reticulum Stress are Involved in Abeta-Induced Pathologies | 2015 | Jun Wan | 20mM PBS |
| Donepezil attenuates Aβ-associated mitochondrial dysfunction and reduces mitochondrial Aβ accumulation in vivo and in vitro | 2015 | Hai Yang Zhang | 250 mM sucrose, 10 mM Tris, 2.5 mM KH<sub>2</sub>PO<sub>4</sub>, PH 7.4 |
| Peptide dimer structure in an Aβ(1-42) fibril visualized with cryo-EM | 2015 | Marcus Fändrich and Nikolaus Grigorieff | 50 mM Tris·HCl, pH 7.4 |
| Steady-state and time-resolved Thioflavin-T fluorescence can report on morphological differences in amyloid fibrils formed by Aβ(1-40) and Aβ(1-42) | 2015 | Elin K. Esbjörner | 50 mM sodium phosphate buffer, pH 7.4 |
| Aβ and NMDAR activation cause mitochondrial dysfunction involving ER calcium release | 2015 | A. Cristina Rego | Ham's F-12 (phenol red-free) |
| Aβ-dependent reduction of NCAM2-mediated synaptic adhesion contributes to synapse loss in Alzheimer's disease | 2015 | Lars M. Ittner, Vladimir Sytnyk | Ham's F-12 (phenol red-free) |
| Dendritic and axonal mechanisms of Ca<sup>2+</sup> elevation impair BDNF transport in Aβ oligomer-treated hippocampal neurons | 2015 | Michael A. Silverman | Ham's F-12 (phenol red-free) |
| Cu<sup>2+</sup> accentuates distinct misfolding of Aβ(1–40) and Aβ(1–42) peptides, and potentiates membrane disruption | 2015 | John H. Viles | Water at pH 10.5, using NaOH |
| Resveratrol inhibits oligomeric Aβ-induced microglial activation via NADPH oxidase | 2015 | Wei-Qi Li | MilliQ water |
| Involvement of Intracellular and Mitochondrial Aβ in the Ameliorative Effects of Huperzine A against Oligomeric Aβ42-Induced Injury in Primary Rat Neurons | 2015 | Hai Yang Zhang | Neurobasal |
| Minocycline attenuates Aβ oligomers-induced pro-inflammatory phenotype in primary microglia while enhancing Aβ fibrils phagocytosis | 2015 | Nabila Hamdi | Oligomers - Ham's F12, Fibrils - PBS |
| Aβ-induced degradation of BMAL1 and CBP leads to circadian rhythm disruption in Alzheimer's disease | 2015 | Inhee Mook-Jung | Opti-MEM |
| Alzheimer-associated Aβ oligomers impact the central nervous system to induce peripheral metabolic deregulation | 2015 | Fernanda G De Felice | PBS |
| Stabilization of Nontoxic Aβ-Oligomers: Insights into the Mechanism of Action of Hydroxyquinolines in Alzheimer's Disease | 2015 | Colin L. Masters | PBS |
| ApoE4 and Aβ Oligomers Reduce BDNF Expression via HDAC Nuclear Translocation | 2015 | Daniel Alkon | PBS without Ca<sup>2+</sup> or Mg<sup>2+</sup> |
| Lycopene abrogates Aβ(1–42)-mediated neuroinflammatory cascade in an experimental model of Alzheimer's disease | 2015 | Kanwaljit Chopra | PBS, pH 7.4 |
| Rapid α-oligomer formation mediated by the Aβ C terminus initiates an amyloid assembly pathway | 2016 | Ronald Wetzel | 1 x PBS, pH 7.4 |
| Heterotypic seeding of Tau fibrillization by pre-aggregated Aβ provides potent seeds for prion-like seeding and propagation of Tau-pathology in vivo | 2016 | Diederik Moechars and Ilse Dewachter | 10 mM HCl solution |
| The new β amyloid-derived peptide Aβ1–6A2V-TAT(D) prevents Aβ oligomer formation and protects transgenic C. elegans from Aβ toxicity | 2016 | Mario Salmona | 10 mM phosphate buffer containing 150 mM NaCl, pH 7.4 |
| Astrocytes from old Alzheimer's disease mice are impaired in Aβ uptake and in neuroprotection | 2016 | Dan Frenkel | 10 mM HEPES, 300 mM NaCl, in DDW, pH = 7.4 |
| Atomic-resolution structure of a disease-relevant Aβ(1–42) amyloid fibril | 2016 | Beat H. Meier and Roland Riek | 100 mM phosphate at pH of 7.4, 0/100 mM NaCl, 0/10/100 μM ZnCl<sub>2</sub> / 30 μM heparin. Either zinc or heparin not both. |
| Structure of Crenezumab Complex with Aβ Shows Loss of β-Hairpin | 2016 | Weiru Wang | 100 mM sodium phosphate buffer, pH 7.4 |
| Mechanisms of tau and Aβ-induced excitotoxicity | 2016 | Gail V.W. Johnson | 135 mM NaCl, 5.4 mM KCl, 0.33 mM NaH<sub>2</sub>PO<sub>4</sub>, 0.4 mM KH<sub>2</sub>PO<sub>4</sub>, 20 mM HEPES, 0.8 mM MgCl<sub>2</sub>, 1.2 mM CaCl<sub>2</sub>, 10 mM glucose, and pH 7.4 |
| Quantitative analysis of intrinsic and extrinsic factors in the aggregation mechanism of Alzheimer-associated Aβ-peptide | 2016 | Sara Linse | 20 mM sodium phosphate buffer, 200 μM EDTA, 0.02% NaN<sub>3</sub>, pH 8 |
| Presynaptic dystrophic neurites surrounding amyloid plaques are sites of microtubule disruption, BACE1 elevation, and increased Aβ generation in Alzheimer's disease | 2016 | Robert Vassar | 4 mM HEPES, pH 8 |
| Soluble Conformers of Aβ and Tau Alter Selective Proteins Governing Axonal Transport | 2016 | Sylvain E. Lesné | 50 mM Tris-HCl, 150 mM NaCl, 0.01% Triton X-100, pH 7.4 |
| Soluble prion protein and its N-terminal fragment prevent impairment of synaptic plasticity by Aβ | 2016 | Witold K. Surewicz | 50 mM sodium phosphate buffer, pH 7.4 |
| oligomers: Implications for novel therapeutic strategy in Alzheimer's disease |  |  |  |
| Aβ-Induced Synaptic Alterations Require the E3 Ubiquitin Ligase Nedd4-1 | 2016 | Gentry N. Patrick | MEM |
| The Tyr216 phosphorylated form of GSK3β contributes to tau phosphorylation at PHF-1 epitope in response to Aβ in the nucleus of SH-SY5Y cells | 2016 | Sabrina Ingrand | MiliQ Water |
| Dopamine agonists rescue Aβ-induced LTP impairment by Src-family tyrosine kinases | 2016 | Klaus G. Reymann | Oligomers - Ham's F12, Fibrils - Neurobasal |
| Identification of a Small Molecule Cyclophilin D Inhibitor for Rescuing Aβ-Mediated Mitochondrial Dysfunction | 2016 | Shirley ShiDu Yan | PBS |
| Antibody modified-silver nanoparticles for colorimetric immuno sensing of Aβ(1-40/1-42) based on the interaction between β-amyloid and Cu<sup>2+</sup> | 2016 | Xiurong Yang | PBS (50 mM NaH<sub>2</sub>PO<sub>4</sub>, 100 mM NaCl, pH 7.6) |
| Conversion of Synthetic Aβ to In Vivo Active Seeds and Amyloid Plaque Formation in a Hippocampal Slice Culture Model | 2016 | Mathias Jucker | PBS (added to cells so presumably pH 7.4) |
| Deacetylation of TFEB promotes fibrillar Aβ degradation by upregulating lysosomal biogenesis in microglia | 2016 | Jianguo Ji | PBS (added to cells so presumably pH 7.4) |
| RXR controlled regulatory networks identified in mouse brain counteract deleterious effects of Aβ oligomers | 2016 | Radosveta Koldamova | serum-free medium |
| Characterization of insulin-degrading enzyme-mediated cleavage of Aβ in distinct aggregation states | 2016 | Kerensa Broerson | ThT/TEM - PBS, pH 7.4 NMR - 50 mM phosphate buffer pH 6.8 containing 50 mM NaCl |
| Novel Curcumin loaded nanoparticles engineered for Blood-Brain Barrier crossing and able to disrupt Abeta aggregates | 2017 | Andreas M. Grabrucker | ThT - 'Assay buffer', Cell culture - NB+++ |
| Astrocytic LRP1 Mediates Brain Aβ Clearance and Impacts Amyloid Deposition | 2017 | Guojun Bu | DMEM with 10% FBS |
| Aβ accumulation causes MVB enlargement and is modelled by dominant negative VPS4A | 2017 | Gunnar K Gouras | |
| High-resolution bioelectrical imaging of Aβ-induced network dysfunction on CMOS-MEAs for neurotoxicity and rescue studies | 2017 | Luca Berdondini | 10 mM phosphate buffer, pH 7.4 |
| The Aβ oligomer eliminating D-enantiomeric peptide RD2 improves cognition without changing plaque pathology | 2017 | Dieter Willbold | 10 mM sodium phosphate buffer, pH 7.4. |
| Human Brain-Derived Aβ Oligomers Bind to Synapses and Disrupt Synaptic Activity in a Manner That Requires APP | 2017 | Tara Spires - Jones and Dominic Walsh | 124 mM NaCl, 2.8 mM KCl, 1.25 mM NaH<sub>2</sub>PO<sub>4</sub>, and 26 mM NaHCO<sub>3</sub>, pH 7.4. |
| A vaccine with Aβ oligomer-specific mimotope attenuates cognitive deficits and brain pathologies in transgenic mice with Alzheimer's disease | 2017 | Rui-tian Liu | 20mM PBS, pH 7.4 |
| Nec-1 alleviates cognitive impairment with reduction of Aβ and tau abnormalities in APP/PS1 mice | 2017 | YoungSoo Kim | DMEM with 0.5%FBS |
| Increased susceptibility to Aβ toxicity in neuronal cultures derived from familial Alzheimer's disease (PSEN1-A246E) induced pluripotent stem cells | 2017 | Claudio Soto | DMEM/F12 without serum |
| Norovirus P particle-based active Aβ immunotherapy elicits sufficient immunogenicity and improves cognitive capacity in a mouse model of Alzheimer's disease | 2017 | Wei Kong | Ham's F-12, ThT - 10 mM PB, 500 mM NaCl, pH 7.0 |
| Inhibition of Drp1 Ameliorates Synaptic Depression, Aβ Deposition, and Cognitive Impairment in an Alzheimer's Disease Model | 2017 | Dong-Gyu Jo | Ham's F-12 (phenol red-free) |
| Role of membrane GM1 on early neuronal membrane actions of Aβ during onset of Alzheimer's disease | 2017 | Luis Aguayo | MiliQ water |
| High-density lipoproteins suppress Aβ-induced PBMC adhesion to human endothelial cells in bioengineered vessels and in monoculture | 2017 | Cheryl L. Wellington | Monomer - DMEM Oligomer - Ham's F-12 (phenol red-free) Fibrils - 0.1μM HCl. |
| Strain-specific Fibril Propagation by an Aβ Dodecamer | 2017 | Vijayaraghavan Rangachari | Monomer - implied PBS LFAO - 50 mM NaCl and 5 mM C12:0 fatty acid, Fibrils - 150 mM NaCl with 0.01% NaN<sub>3</sub> |
| Astrocyte Transforming Growth Factor Beta 1 Protects Synapses against Aβ Oligomers in Alzheimer's Disease Model | 2017 | Flávia Carvalho Alcantara Gomes | PBS |
| Aβ seeding potency peaks in the early stages of cerebral β-amyloidosis | 2017 | Mathias Jucker | PBS (10%, w/v) |
| Colorimetric sandwich immunosensor for Aβ(1-42) based on dual antibody-modified gold nanoparticles | 2017 | Xiurong Yang | PBS (50 mM NaH<sub>2</sub>PO<sub>4</sub>, 100 mM NaCl), pH 7.4 |
| Involvement of GluN2B subunit containing N-methyl-d-aspartate (NMDA) receptors in mediating the acute and chronic synaptotoxic effects of oligomeric amyloid-beta (Aβ) in murine models of Alzheimer's disease (AD) | 2017 | Chris Parsons | Ringer's solution: 125 mM NaCl, 2.5 mM KCl, 25 mM NaHCO<sub>3</sub>, 2 mM CaCl<sub>2</sub>, 1 mM MgCl<sub>2</sub>, 25 mM d-glucose, and 1.25 mM NaH<sub>2</sub>PO<sub>4</sub>, bubbled with a 95% O<sub>2</sub>, 5% CO<sub>2</sub> mixture, pH of 7.3 |
| Anthocyanin-Loaded PEG-Gold Nanoparticles Enhanced the Neuroprotection of Anthocyanins in an Aβ1-42 Mouse Model of Alzheimer's Disease | 2017 | Myeong Ok Kim | Saline |
| A Novel, Multi-Target Natural Drug Candidate, Matrine, Improves Cognitive Deficits in Alzheimer's Disease Transgenic Mice by Inhibiting Aβ Aggregation and Blocking the RAGE/Aβ Axis | 2017 | Bin Zhao | ThT - 50-mM phosphate buffer (pH 6.0). Other experiments unclear |
| Amyloid β-Derived Diffusible Ligands (ADDLs) Induce Abnormal Autophagy Associated with Aβ Aggregation Degree | 2018 | Tao Hung | DMEM/F12 medium without phenol red |
| Inhibition of Phosphodiesterase-4 Reverses Aβ-Induced Memory Impairment by Regulation of HPA Axis Related cAMP Signaling | 2018 | Jiangchun Pan | 0.9% saline |
| The Aβ protofibril selective antibody mAb158 prevents accumulation of Aβ in astrocytes and rescues neurons from Aβ-induced cell death | 2018 | Anna Erlandsson | 10X PBS |
| Diffusible, highly bioactive oligomers represent a critical minority of soluble Aβ in Alzheimer's disease brain | 2018 | Dominic Walsh | 124 mM NaCl, 2.8 mM KCl, 1.25 mM NaH<sub>2</sub>PO<sub>4</sub>, 26 mM NaHCO<sub>3</sub>, pH 7.4 |
| Aβ truncated species: Implications for brain clearance mechanisms and amyloid plaque deposition | 2018 | Jorge Ghiso | CD- 10 mM phosphate buffer, pH 7.4, containing 150 mM sodium fluoride, ThT- 50 mM Tris-HCl buffer, pH 8.5 |
| CaMKII Metaplasticity Drives Aβ Oligomer-Mediated Synaptotoxicity | 2018 | Daniel Choquet | F12 medium |
| Soluble Aβ1-42 increases the heterogeneity in synaptic vesicle pool size among synapses by suppressing intersynaptic vesicle sharing | 2018 | Sunghoe Chang | Ham's F-12 (phenol red-free) |
| A novel Aβ epitope vaccine based on bacterium-like particle against Alzheimer's disease | 2018 | Wei Kong | Ham's F-12 (phenol red-free) |
| HDL Mimetic Peptide 4F Mitigates Aβ-Induced Inhibition of ApoE Secretion and Lipidation in Primary Astrocytes and Microglia | 2018 | Ling LI | Ham's F-12 (phenol red-free) |
| Salvianolic acid B attenuates mitochondrial stress against Aβ toxicity in primary cultured mouse neurons | 2018 | Heng Du | Ham's F-12 (phenol red-free) |
| Effect of Cholesterol on Membrane Fluidity and Association of Aβ Oligomers and Subsequent Neuronal Damage: A Double-Edged Sword | 2018 | Luis Aguayo | MiliQ water |
| Effects of n-3 PUFA enriched and n-3 PUFA deficient diets in naïve and Aβ-treated female rats | 2018 | Lugia Trabace | MiliQ water |
| Estrogen deficiency exacerbates Aβ-induced memory impairment through enhancement of neuroinflammation, amyloidogenesis and NF-κB activation in ovariectomized mice | 2018 | Jin Tae Hong | MiliQ water |
| Enantiomeric Aβ peptides inhibit the fluid shear stress response of PIEZO1 | 2018 | Philip Gottlieb | Monomer - THIP followed by DMSO then buffer as below, oligomer - 150 KCl, 10 HEPES, 1 MgCl<sub>2</sub>, 1 CaCl<sub>2</sub> at pH 7.4 |
| Astrocytic glutamatergic transporters are involved in Aβ-induced synaptic dysfunction | 2018 | Pingyi Xu | PBS |
| High-affinity interactions and signal transduction between Aβ oligomers and TREM2 | 2018 | Todd Golde | PBS |
| Estrogen Receptors Are Involved in the Neuroprotective Effect of Silibinin in Aβ1–42-Treated Rats | 2018 | Takeshi Ikejima | PBS, pH 7.4 |
| The diphenylpyrazole compound anle138b blocks Aβ channels and rescues disease phenotypes in a mouse model for amyloid pathology | 2018 | Andre Fischer | PBS, pH 7.4 |
| The NMDA receptor antagonist Radiprodil reverses the synaptotoxic effects of different amyloid-beta (Aβ) species on long-term potentiation (LTP) | 2018 | Chris Parsons | Ringer's solution: 125 mM NaCl, 2.5 mM KCl, 25 mM NaHCO<sub>3</sub>, 2 mM CaCl<sub>2</sub>, 1 mM MgCl<sub>2</sub>, 25 mM d-glucose, and 1.25 mM NaH<sub>2</sub>PO<sub>4</sub>, bubbled with a 95% O<sub>2</sub>, 5% CO<sub>2</sub> mixture, pH of 7.3 |
| Euxanthone Attenuates Aβ1–42-Induced Oxidative Stress and Apoptosis by Triggering Autophagy | 2018 | Gang Sun | Saline |
| Cubeben induces autophagy via PI3K-AKT-mTOR pathway to protect primary neurons against amyloid beta in Alzheimer's disease | 2019 | Ruijan Dong |  |
| Neuroprotective efficacy of thymoquinone against amyloid beta-induced neurotoxicity in human induced pluripotent stem cell-derived cholinergic neurons | 2019 | Ikuro Suzuki | Implied cell culture medium but unclear |
| A catalytic antioxidant for limiting amyloid-beta peptide aggregation and reactive oxygen species generation | 2019 | Tim Storr | 0.01M PBS pH7.4 |
| Serotonin type 6 receptor antagonist attenuates the impairment of long-term potentiation and memory induced by Abeta | 2019 | Alireza Komaki | 0.9% saline, pH 7.4 |
| Exendin-4 attenuates brain mitochondrial toxicity through PI3K/Akt-dependent pathway in amyloid beta (1–42)-induced cognitive deficit rats | 2019 | Jaya Verma | 147 mM NaCl, 2.9 mM KCl, 1.6 mM MgCl<sub>2</sub>, 1.7 mM CaCl<sub>2</sub> and 2.2 mM dextrose; pH 7.4 |
| Interactions of polyunsaturated fatty acids with amyloid peptides Aβ40 and Aβ42 | 2019 | Praveen Rao | 215mM sodium phosphate, pH 7.4 |
| Morphological changes induced in erythrocyte by amyloid beta peptide and glucose depletion: A combined atomic force microscopy and biochemical study | 2019 | Marco Girasole | 35 mM Na<sub>2</sub>SO<sub>4</sub>, 90 mM NaCl, 25 mM HEPES [N-(2-hydroxyethyl)-piperazine-N1-2-ethanesulfonic acid], 1.5 mM MgCl<sub>2</sub>, glucose 5 mM |
| Cryo-EM structure and polymorphism of Aβ amyloid fibrils purified from Alzheimer's brain tissue | 2019 | Mathias Jucker & Marcus Fändrich | 50 mM Tris, 10 mM ethylenediaminetetraacetic acid, pH 8 |
| Carbenoxolone Reverses the Amyloid Beta 1–42 Oligomer-Induced Oxidative Damage and Anxiety-Related Behavior in Rats | 2019 | Bimla Nehru | First dissolved in 100μl 100mM NaOH and further diluted in 900μl 10mM sodium phosphate |
| Amyloid beta-mediated KIF5A deficiency disrupts anterograde axonal mitochondrial movement | 2019 | Lan Guo | Ham's F-12 |
| Senolytic therapy alleviates Aβ-associated oligodendrocyte progenitor cell senescence and cognitive deficits in an Alzheimer's disease model | 2019 | Mark Matteson | Incubated in DMSO for 24hrs then diluted in DMEM, 20% FBS, L-glut, 1 mM sodium pyruvate, 0.1 mM β-mercaptoethanol, 0.1 mM non-essential amino acids, and 1,000 U ml<sup>-1</sup> leukemia inhibitory factor |
| 9-Methylfascaplysin Is a More Potent Aβ Aggregation Inhibitor than the Marine-Derived Alkaloid, Fascaplysin, and Produces Nanomolar Neuroprotective Effects in SH-SY5Y Cells | 2019 | Hongze Lang | MiliQ water/ PBS |
| In vitro studies of the neuroprotective activities of astaxanthin and fucoxanthin against amyloid beta (Aβ1–42) toxicity and aggregation | 2019 | Wei Zhang | PBS |
| Alborexin clears amyloid-β by inducing autophagy through PTEN-mediated inhibition of the AKT pathway | 2019 | Ram Vishwakarma, Ajay Kumar | PBS |
| Mesencephalic astrocyte-derived neurotrophic factor (MANF) protects against Aβ toxicity via attenuating Aβ-induced endoplasmic reticulum stress | 2019 | Yuxian Shen | PBS |
| α-Sheet secondary structure in amyloid β-peptide drives aggregation and toxicity in Alzheimer's disease | 2019 | Valarie Daggett | PBS, pH 7.6 |
| Amyloid-beta induced retrograde axonal degeneration in a mouse tauopathy model | 2019 | Shu-Wei Sun | Saline |
| Hesperetin Confers Neuroprotection by Regulating Nrf2/TLR4/NF-κB Signaling in an Aβ Mouse Model | 2019 | Myeong Ok Kim | Saline |
| Trodusquemine enhances Aβ42 aggregation but suppresses its toxicity by displacing oligomers from cell membranes | 2019 | Christopher Dobson | ThT - 5 mM sodium phosphate, 200 μM EDTA, pH 8.0 in the presence of either 75 or 150 mM NaCl, NMR- 5mM sodium phosphate, pH 7.4 |
| Inhibition of amyloid beta toxicity in zebrafish with a chaperone-gold nanoparticle dual strategy | 2019 | Sijie Lin | Undefined aqueous solution (ThT/TEM) deionized water (CD) PBS pH 7.4 |
| Designed Cell-Penetrating Peptide Inhibitors of Amyloid-beta Aggregation and Cytotoxicity | 2020 | Mazin Magzoub | PBS or DMEM (no serum). |
| Interaction between tissue transglutaminase and amyloid-beta: Protein-protein binding versus enzymatic crosslinking | 2020 | Benjamin Drukarch | 10 mM sodium-acetate buffer, pH 5.0 (SPR) / 70 mM NaCl, 2.5 mM CaCl<sub>2</sub>, 40 mM HEPES, pH 7.5, 1 mM DTT (cross-linking) |
| Amyloid-Beta Peptides Trigger Aggregation of Alpha-Synuclein In Vitro | 2020 | Stephan Schilling | 20 mM Tris/HCl, 100 mM NaCl, pH 7.0 |
| Sinomenine inhibits amyloid beta-induced astrocyte activation and protects neurons against indirect toxicity | 2020 | Shiv Kumar Sharma | PBS |

**Supplementary Table S2:** Morphometric parameters for individually 3D-reconstructed Aβ_42_ fibrils. Fibrils numbered #1-100 originate from 20mM sodium phosphate pH 8.0 assembly condition, fibrils #101-200 are from the sodium phosphate pH 7.4 assembly condition, fibrils #201-300 are from HEPES pH 7.4 assembly condition and fibrils #301-400 are from Tris pH 7.4 assembly condition. For each fibril, the following parameters are listed. *L*: Contour length of the segment of the fibril on AFM height image that was traced and its surface envelope 3D-reconstructed. *h*: The average height of the fibril segment. *cod*: The mean cross-over distance of peaks on the centre fibril height profile. *dpf*: directional periodic frequency. *hnd*: twist handedness of the fibril. *csa*: The filament mean AFM tip accessible cross-sectional area of the fibril. csr: The filament mean cross-sectional radius to the helical axis. csjz: The filament cross-sectional mean second polar moment of area. The fibril numbers is the index number of each of the individual fibrils and were used throughout.

| Fibril # | *L* / nm | *h* / nm | *cod* / nm | *hnd* | *dpf* / nm<sup>-1</sup> | *csa* / nm<sup>2</sup> | *csr* / nm | *csjz* / nm<sup>4</sup> |
| --- | --- | --- | --- | --- | --- | --- | --- | --- |
| 1 | 992.20 | 6.35 | 69.35 | left | -0.01442 | 29.77 | 3.00 | 140.84 |
| 2 | 482.45 | 7.62 | 60.49 | left | -0.01653 | 41.96 | 3.56 | 290.74 |
| 3 | 761.74 | 5.91 | 105.24 | left | -0.00950 | 25.89 | 2.80 | 108.56 |
| 4 | 240.25 | 6.60 | 75.88 | left | -0.01318 | 27.14 | 2.74 | 171.89 |
| 5 | 2929.72 | 4.97 | 44.65 | left | -0.02240 | 16.36 | 2.21 | 48.15 |
| 6 | 517.59 | 4.97 | 55.14 | left | -0.01813 | 14.08 | 1.97 | 42.76 |
| 7 | 308.60 | 4.66 | 50.63 | left | -0.00987 | 12.11 | 1.83 | 28.70 |
| 8 | 324.25 | 6.11 | 60.94 | left | -0.00821 | 22.72 | 2.54 | 99.15 |
| 9 | 740.24 | 7.25 | 124.22 | left | -0.00403 | 38.27 | 3.41 | 234.54 |
| 10 | 623.07 | 5.74 | 53.16 | left | -0.01881 | 19.53 | 2.23 | 56.55 |
| 11 | 1154.33 | 5.71 | 123.11 | left | -0.00812 | 24.44 | 2.75 | 91.03 |
| 12 | 546.91 | 7.28 | 60.99 | left | -0.01640 | 40.73 | 3.44 | 256.09 |
| 13 | 521.50 | 4.83 | 39.58 | left | -0.02527 | 15.59 | 2.13 | 43.44 |
| 14 | 501.98 | 8.59 | 53.00 | left | -0.01887 | 55.56 | 4.04 | 441.25 |
| 15 | 595.73 | 7.80 | 117.15 | left | -0.00854 | 34.37 | 3.14 | 236.05 |
| 16 | 388.69 | 7.70 | 98.96 | left | -0.00505 | 31.12 | 3.01 | 178.83 |
| 17 | 1033.27 | 5.78 | 67.88 | left | -0.01473 | 24.64 | 2.61 | 84.04 |
| 18 | 453.14 | 4.42 | 67.81 | left | -0.00737 | 9.82 | 1.69 | 16.84 |
| 19 | 816.42 | 8.51 | 113.74 | left | -0.00879 | 38.45 | 3.29 | 279.31 |
| 20 | 265.64 | 6.76 | 39.34 | right | 0.01271 | 30.54 | 3.06 | 148.20 |
| 21 | 603.53 | 5.11 | 47.62 | left | -0.02100 | 15.64 | 2.10 | 48.67 |
| 22 | 779.32 | 5.94 | 140.82 | left | -0.00710 | 23.01 | 2.59 | 97.50 |
| 23 | 232.43 | 4.79 | 36.17 | left | -0.02765 | 12.75 | 1.92 | 30.92 |
| 24 | 814.46 | 7.57 | 108.37 | left | -0.00923 | 38.91 | 3.40 | 249.27 |
| 25 | 355.49 | 7.65 | 94.99 | left | -0.00526 | 23.66 | 2.56 | 104.01 |
| 26 | 320.36 | 4.14 | 68.64 | left | -0.01457 | 10.28 | 1.60 | 28.90 |
| 27 | 390.70 | 6.95 | 57.36 | left | -0.01743 | 25.22 | 2.49 | 153.20 |
| 28 | 267.58 | 5.05 | 32.37 | right | 0.03090 | 15.82 | 2.20 | 43.75 |
| 29 | 361.35 | 7.26 | 85.16 | left | -0.01174 | 31.04 | 2.93 | 220.18 |
| 30 | 1507.99 | 6.96 | 108.04 | right | 0.00926 | 40.77 | 3.31 | 253.62 |
| 31 | 334.00 | 5.24 | 67.43 | left | -0.01483 | 15.52 | 2.07 | 47.92 |
| 32 | 601.58 | 5.02 | 42.88 | left | -0.02332 | 17.42 | 2.26 | 54.73 |
| 33 | 992.24 | 7.23 | 105.08 | left | -0.00952 | 30.88 | 3.02 | 173.52 |
| 34 | 605.48 | 4.87 | 76.20 | left | -0.01312 | 12.60 | 1.92 | 30.72 |
| 35 | 599.62 | 8.46 | 57.94 | left | -0.00863 | 38.91 | 3.42 | 264.93 |
| 36 | 1025.41 | 6.34 | 110.57 | right | 0.00904 | 28.34 | 2.94 | 128.52 |
| 37 | 955.14 | 6.14 | 131.15 | left | -0.00762 | 24.02 | 2.68 | 106.98 |
| 38 | 675.82 | 5.18 | 43.63 | left | -0.02292 | 16.46 | 2.10 | 50.78 |
| 39 | 455.11 | 6.28 | 98.68 | left | -0.01013 | 20.30 | 2.28 | 95.90 |
| 40 | 418.04 | 7.57 | 87.96 | left | -0.01137 | 27.09 | 2.68 | 172.74 |
| 41 | 414.09 | 10.03 | 74.02 | left | -0.00675 | 58.48 | 4.15 | 558.07 |
| 42 | 382.81 | 5.99 | 46.93 | left | -0.02131 | 17.82 | 2.23 | 69.34 |
| 43 | 521.50 | 6.42 | 139.20 | left | -0.00718 | 31.39 | 3.05 | 182.43 |
| 44 | 630.87 | 5.83 | 125.63 | left | -0.00796 | 21.45 | 2.51 | 85.03 |
| 45 | 392.68 | 4.75 | 31.02 | left | -0.03224 | 15.19 | 2.15 | 40.49 |
| 46 | 335.95 | 4.91 | 53.28 | left | -0.01877 | 11.40 | 1.73 | 31.55 |
| 47 | 583.99 | 4.48 | 88.28 | left | -0.01133 | 11.29 | 1.81 | 23.18 |
| 48 | 509.79 | 4.36 | 49.65 | left | -0.02014 | 14.41 | 2.03 | 33.10 |
| 49 | 515.64 | 6.47 | 72.79 | left | -0.01374 | 27.26 | 2.64 | 116.92 |
| 50 | 980.47 | 4.93 | 38.05 | left | -0.02628 | 15.74 | 2.21 | 41.91 |
| 51 | 453.16 | 5.05 | 40.41 | left | -0.02475 | 15.75 | 2.20 | 43.39 |
| 52 | 855.48 | 5.05 | 36.70 | left | -0.02725 | 17.46 | 2.32 | 51.52 |
| 53 | 437.52 | 5.58 | 46.09 | left | -0.01085 | 16.51 | 2.19 | 55.42 |
| 54 | 1570.33 | 5.60 | 47.93 | left | -0.02086 | 19.43 | 2.38 | 78.75 |
| 55 | 302.74 | 4.32 | 56.40 | left | -0.01773 | 11.48 | 1.82 | 26.53 |
| 56 | 558.64 | 9.50 | 78.58 | left | -0.01273 | 48.14 | 3.45 | 524.60 |
| 57 | 587.92 | 5.80 | 57.27 | left | -0.01746 | 18.41 | 2.20 | 71.14 |
| 58 | 373.06 | 7.76 | 55.31 | left | -0.01808 | 42.31 | 3.58 | 291.03 |
| 59 | 1048.85 | 4.75 | 39.05 | left | -0.02561 | 14.37 | 2.07 | 36.76 |
| 60 | 1128.96 | 10.05 | 124.56 | left | -0.00803 | 53.27 | 3.74 | 725.59 |
| 61 | 634.78 | 5.27 | 62.13 | left | -0.01610 | 14.10 | 1.96 | 48.14 |
| 62 | 949.24 | 6.06 | 39.46 | left | -0.02534 | 22.85 | 2.60 | 102.55 |
| 63 | 1304.78 | 4.09 | 24.33 | right | 0.04110 | 8.93 | 1.52 | 14.81 |
| 64 | 261.72 | 5.97 | 39.91 | left | -0.02506 | 20.97 | 2.47 | 91.28 |
| 65 | 269.53 | 4.68 | 58.89 | left | -0.01698 | 10.70 | 1.73 | 25.74 |
| 66 | 408.21 | 8.38 | 108.79 | right | 0.00919 | 36.98 | 3.20 | 258.27 |
| 67 | 421.88 | 6.55 | 61.17 | right | 0.01635 | 28.04 | 2.88 | 141.56 |
| 68 | 918.04 | 6.25 | 90.43 | right | 0.01106 | 23.73 | 2.50 | 134.45 |
| 69 | 1162.11 | 6.24 | 92.58 | right | 0.00540 | 20.57 | 2.51 | 70.78 |
| 70 | 468.76 | 4.97 | 46.27 | left | -0.02161 | 13.72 | 1.94 | 40.53 |
| 71 | 166.02 | 6.09 | 15.16 | left | -0.02199 | 27.27 | 2.93 | 120.28 |
| 72 | 837.92 | 8.61 | 86.35 | left | -0.01158 | 47.54 | 3.77 | 403.25 |
| 73 | 472.67 | 5.64 | 50.95 | left | -0.01963 | 19.36 | 2.33 | 71.76 |
| 74 | 310.55 | 5.80 | 46.39 | left | -0.01078 | 19.21 | 2.39 | 74.06 |
| 75 | 333.99 | 5.04 | 41.29 | left | -0.02422 | 16.42 | 2.20 | 51.93 |
| 76 | 503.93 | 7.55 | 82.52 | left | -0.01212 | 35.25 | 3.14 | 265.10 |
| 77 | 240.24 | 5.07 | 52.05 | left | -0.01921 | 15.40 | 2.11 | 47.75 |
| 78 | 890.63 | 3.55 | 55.13 | right | 0.01814 | 9.91 | 1.76 | 15.48 |
| 79 | 617.35 | 7.47 | 60.06 | left | -0.01665 | 30.80 | 2.93 | 177.06 |
| 80 | 548.97 | 6.01 | 28.16 | right | 0.03551 | 25.90 | 2.70 | 99.60 |
| 81 | 339.93 | 5.32 | 75.86 | left | -0.01318 | 16.60 | 2.15 | 55.25 |
| 82 | 425.90 | 7.42 | 53.33 | left | -0.01875 | 44.17 | 3.64 | 289.72 |
| 83 | 513.81 | 7.43 | 93.68 | left | -0.01068 | 32.05 | 3.05 | 188.42 |
| 84 | 259.84 | 6.37 | 70.55 | left | -0.01417 | 18.79 | 2.10 | 104.64 |
| 85 | 314.82 | 5.37 | 58.84 | left | -0.01699 | 15.21 | 2.06 | 54.10 |
| 86 | 392.68 | 4.66 | 42.61 | left | -0.02347 | 13.69 | 1.98 | 38.18 |
| 87 | 248.11 | 5.96 | 50.84 | left | -0.01967 | 20.62 | 2.48 | 82.71 |
| 88 | 203.19 | 5.70 | 55.87 | left | -0.00895 | 14.67 | 2.02 | 47.30 |
| 89 | 421.99 | 4.82 | 37.15 | left | -0.02692 | 15.84 | 2.20 | 42.50 |
| 90 | 482.54 | 8.07 | 39.98 | left | -0.01251 | 47.09 | 3.84 | 360.72 |
| 91 | 453.24 | 5.46 | 53.72 | left | -0.01861 | 17.75 | 2.27 | 63.60 |
| 92 | 212.95 | 5.25 | 41.76 | left | -0.02395 | 14.73 | 1.98 | 53.36 |
| 93 | 377.05 | 6.31 | 109.11 | left | -0.00917 | 21.66 | 2.49 | 100.14 |
| 94 | 1488.68 | 8.07 | 100.12 | left | -0.00999 | 49.10 | 3.82 | 360.05 |
| 95 | 435.67 | 8.68 | 120.54 | left | -0.00830 | 46.28 | 3.62 | 423.99 |
| 96 | 384.91 | 6.89 | 85.57 | left | -0.01169 | 22.26 | 2.35 | 142.28 |
| 97 | 574.37 | 6.36 | 49.32 | left | -0.02028 | 25.10 | 2.58 | 93.11 |
| 98 | 636.89 | 5.38 | 148.31 | left | -0.00674 | 18.97 | 2.38 | 62.14 |
| 99 | 339.94 | 7.77 | 84.40 | left | -0.01185 | 36.24 | 3.06 | 249.09 |
| 100 | 332.12 | 4.80 | 40.10 | left | -0.02494 | 14.13 | 2.05 | 36.76 |
| 101 | 437.51 | 4.81 | 70.00 | left | -0.01429 | 14.84 | 2.08 | 41.82 |
| 102 | 236.34 | 5.08 | 33.85 | left | -0.02954 | 22.48 | 2.61 | 86.99 |
| 103 | 380.87 | 4.74 | 44.82 | left | -0.02231 | 18.10 | 2.27 | 52.24 |
| 104 | 294.93 | 5.31 | 11.21 | left | -0.04460 | 20.78 | 2.56 | 67.90 |
| 105 | 332.03 | 4.36 | 31.71 | left | -0.03153 | 16.27 | 2.21 | 38.17 |
| 106 | 375.01 | 6.44 | 45.65 | left | -0.02191 | 33.66 | 3.10 | 158.53 |
| 107 | 585.94 | 5.48 | 48.01 | left | -0.02083 | 18.75 | 2.34 | 67.16 |
| 108 | 515.63 | 7.22 | 41.18 | left | -0.02428 | 33.44 | 3.18 | 199.28 |
| 109 | 259.77 | 7.34 | 57.81 | left | -0.00865 | 26.57 | 2.72 | 149.41 |
| 110 | 312.51 | 4.16 | 50.00 | left | -0.02000 | 11.13 | 1.75 | 23.40 |
| 111 | 644.55 | 6.35 | 40.53 | right | 0.02467 | 30.96 | 2.98 | 142.72 |
| 112 | 337.90 | 5.68 | 51.07 | left | -0.01958 | 18.97 | 2.29 | 65.16 |
| 113 | 453.15 | 6.88 | 74.80 | left | -0.00668 | 29.93 | 2.92 | 175.89 |
| 114 | 513.70 | 4.77 | 76.76 | left | -0.01303 | 18.80 | 2.39 | 52.22 |
| 115 | 359.34 | 4.36 | 47.66 | left | -0.02098 | 12.53 | 1.90 | 27.46 |
| 116 | 408.22 | 4.70 | 14.64 | left | -0.06830 | 16.19 | 2.21 | 43.28 |
| 117 | 451.18 | 4.22 | 53.71 | left | -0.01862 | 13.19 | 1.95 | 31.52 |
| 118 | 314.46 | 6.12 | 81.32 | left | -0.01230 | 32.74 | 3.15 | 170.96 |
| 119 | 392.60 | 7.63 | 83.30 | left | -0.00400 | 36.90 | 3.33 | 228.44 |
| 120 | 507.82 | 7.54 | 75.10 | left | -0.00666 | 34.58 | 3.23 | 201.48 |
| 121 | 297.02 | 4.43 | 50.39 | left | -0.01984 | 11.80 | 1.83 | 29.82 |
| 122 | 185.58 | 5.99 | 35.84 | left | -0.02790 | 18.54 | 2.23 | 77.32 |
| 123 | 359.39 | 4.91 | 60.40 | left | -0.01656 | 14.79 | 2.02 | 41.13 |
| 124 | 269.54 | 4.84 | 22.41 | right | 0.04463 | 10.77 | 1.67 | 28.08 |
| 125 | 236.35 | 5.30 | 55.53 | left | -0.01801 | 18.85 | 2.31 | 70.18 |
| 126 | 507.83 | 6.81 | 50.00 | left | -0.02000 | 36.39 | 3.24 | 218.56 |
| 127 | 474.61 | 3.95 | 126.97 | left | -0.00788 | 10.19 | 1.63 | 21.73 |
| 128 | 212.89 | 5.55 | 47.98 | left | -0.02084 | 18.89 | 2.38 | 67.69 |
| 129 | 484.38 | 5.35 | 22.25 | left | -0.04495 | 22.94 | 2.68 | 82.38 |
| 130 | 404.31 | 5.79 | 52.32 | left | -0.01911 | 22.42 | 2.54 | 93.83 |
| 131 | 376.96 | 5.13 | 60.08 | left | -0.01664 | 14.32 | 2.02 | 43.87 |
| 132 | 318.37 | 4.96 | 38.84 | left | -0.01287 | 19.09 | 2.36 | 58.59 |
| 133 | 310.55 | 5.64 | 39.73 | left | -0.02517 | 21.85 | 2.51 | 80.39 |
| 134 | 304.72 | 5.22 | 58.01 | left | -0.00862 | 15.55 | 2.14 | 44.76 |
| 135 | 287.17 | 5.10 | 66.46 | left | -0.01505 | 17.63 | 2.25 | 61.37 |
| 136 | 335.94 | 5.19 | 63.92 | left | -0.01565 | 16.72 | 2.24 | 52.10 |
| 137 | 341.80 | 7.08 | 40.06 | left | -0.01248 | 24.73 | 2.67 | 115.81 |
| 138 | 560.55 | 6.49 | 45.28 | left | -0.01104 | 28.42 | 2.96 | 131.20 |
| 139 | 367.19 | 4.78 | 44.95 | left | -0.02225 | 14.99 | 2.13 | 38.12 |
| 140 | 230.47 | 4.69 | 38.31 | right | 0.02610 | 14.33 | 2.07 | 35.84 |
| 141 | 228.52 | 5.91 | 37.27 | left | -0.01342 | 25.88 | 2.78 | 117.98 |
| 142 | 478.53 | 4.95 | 31.13 | left | -0.03212 | 17.67 | 2.31 | 55.50 |
| 143 | 253.91 | 7.30 | 70.70 | left | -0.01414 | 24.12 | 2.52 | 137.88 |
| 144 | 580.24 | 6.48 | 61.30 | left | -0.01631 | 25.46 | 2.72 | 121.93 |
| 145 | 300.87 | 7.36 | 72.02 | left | -0.01388 | 31.10 | 2.87 | 220.59 |
| 146 | 566.57 | 8.86 | 83.32 | left | -0.00600 | 42.48 | 3.48 | 298.28 |
| 147 | 384.88 | 6.20 | 44.91 | left | -0.02227 | 27.21 | 2.84 | 131.78 |
| 148 | 578.32 | 8.31 | 52.47 | left | -0.01906 | 45.30 | 3.38 | 300.73 |
| 149 | 443.54 | 6.71 | 41.00 | left | -0.02439 | 41.68 | 3.51 | 260.07 |
| 150 | 439.62 | 5.05 | 56.51 | left | -0.01769 | 19.45 | 2.34 | 78.00 |
| 151 | 334.08 | 5.49 | 62.03 | left | -0.01612 | 19.21 | 2.38 | 67.58 |
| 152 | 461.06 | 7.56 | 61.15 | right | 0.01635 | 41.48 | 3.50 | 283.69 |
| 153 | 970.95 | 6.59 | 52.13 | left | -0.01918 | 30.56 | 3.08 | 156.69 |
| 154 | 629.06 | 6.19 | 55.62 | left | -0.01798 | 24.75 | 2.74 | 109.61 |
| 155 | 779.49 | 6.39 | 50.31 | left | -0.00994 | 25.41 | 2.79 | 111.53 |
| 156 | 863.51 | 9.17 | 84.22 | left | -0.01187 | 48.89 | 3.62 | 516.99 |
| 157 | 435.66 | 6.99 | 83.76 | left | -0.01194 | 23.55 | 2.40 | 151.15 |
| 158 | 400.50 | 6.24 | 82.05 | left | -0.01219 | 24.52 | 2.42 | 109.67 |
| 159 | 420.15 | 7.41 | 75.17 | left | -0.00665 | 28.82 | 2.80 | 138.99 |
| 160 | 461.07 | 5.97 | 43.62 | left | -0.01146 | 20.99 | 2.43 | 74.42 |
| 161 | 900.62 | 6.05 | 53.91 | left | -0.01855 | 22.85 | 2.60 | 94.72 |
| 162 | 449.34 | 5.68 | 49.13 | left | -0.02035 | 24.15 | 2.70 | 96.57 |
| 163 | 255.93 | 5.51 | 48.76 | left | -0.02051 | 19.98 | 2.34 | 78.22 |
| 164 | 240.32 | 5.27 | 37.63 | right | 0.01329 | 14.92 | 1.97 | 50.52 |
| 165 | 525.55 | 8.25 | 88.93 | right | 0.01125 | 38.16 | 3.28 | 290.12 |
| 166 | 330.20 | 5.78 | 26.19 | left | -0.01909 | 19.38 | 2.38 | 68.49 |
| 167 | 385.04 | 5.79 | 30.88 | left | -0.03238 | 23.72 | 2.66 | 99.82 |
| 168 | 531.42 | 5.76 | 31.83 | left | -0.01571 | 18.65 | 2.32 | 59.28 |
| 169 | 205.14 | 6.13 | 18.54 | right | 0.05394 | 30.54 | 3.06 | 139.93 |
| 170 | 236.39 | 5.49 | 40.28 | left | -0.02482 | 18.82 | 2.37 | 62.70 |
| 171 | 212.95 | 5.96 | 49.36 | left | -0.02026 | 25.40 | 2.83 | 103.38 |
| 172 | 181.85 | 5.23 | 25.79 | right | 0.03878 | 17.34 | 2.32 | 49.44 |
| 173 | 568.50 | 5.33 | 77.59 | left | -0.01289 | 17.48 | 2.30 | 50.32 |
| 174 | 187.55 | 5.90 | 35.41 | left | -0.01412 | 23.41 | 2.69 | 86.55 |
| 175 | 402.44 | 5.56 | 53.95 | left | -0.01854 | 12.88 | 1.96 | 29.23 |
| 176 | 679.87 | 5.19 | 54.83 | left | -0.01824 | 14.54 | 1.99 | 49.61 |
| 177 | 402.45 | 5.17 | 42.25 | left | -0.02367 | 18.88 | 2.39 | 63.52 |
| 178 | 533.34 | 5.96 | 49.88 | left | -0.01002 | 30.19 | 3.04 | 158.30 |
| 179 | 554.84 | 5.96 | 44.44 | left | -0.01125 | 25.22 | 2.76 | 112.81 |
| 180 | 406.35 | 5.41 | 37.16 | left | -0.02691 | 21.16 | 2.55 | 74.29 |
| 181 | 332.12 | 4.91 | 51.48 | left | -0.01943 | 15.97 | 2.14 | 51.85 |
| 182 | 611.48 | 5.93 | 54.43 | left | -0.01837 | 23.39 | 2.65 | 104.59 |
| 183 | 236.99 | 4.92 | 39.70 | right | 0.02519 | 21.72 | 2.59 | 80.17 |
| 184 | 748.24 | 6.07 | 48.36 | left | -0.02068 | 28.11 | 2.94 | 134.94 |
| 185 | 314.55 | 6.45 | 52.32 | left | -0.01911 | 36.10 | 3.14 | 215.90 |
| 186 | 310.63 | 7.63 | 60.07 | left | -0.01665 | 40.12 | 3.49 | 290.35 |
| 187 | 539.22 | 5.89 | 52.31 | left | -0.01912 | 32.80 | 3.17 | 181.34 |
| 188 | 445.43 | 5.99 | 48.67 | left | -0.02055 | 25.59 | 2.78 | 111.78 |
| 189 | 818.58 | 4.78 | 57.15 | left | -0.01750 | 13.62 | 1.89 | 38.22 |
| 190 | 423.94 | 5.78 | 41.32 | left | -0.02420 | 26.70 | 2.84 | 129.76 |
| 191 | 703.30 | 6.29 | 48.60 | left | -0.02058 | 24.50 | 2.73 | 107.88 |
| 192 | 668.14 | 4.98 | 50.76 | left | -0.01970 | 14.78 | 2.07 | 42.54 |
| 193 | 336.03 | 4.95 | 27.47 | right | 0.01820 | 14.60 | 2.06 | 35.19 |
| 194 | 550.94 | 5.30 | 113.80 | left | -0.00879 | 17.35 | 2.13 | 63.89 |
| 195 | 427.85 | 5.76 | 50.65 | left | -0.00987 | 22.75 | 2.61 | 92.74 |
| 196 | 351.67 | 9.18 | 77.95 | left | -0.01283 | 37.16 | 3.00 | 364.54 |
| 197 | 320.40 | 4.20 | 111.45 | left | -0.00897 | 10.01 | 1.64 | 27.34 |
| 198 | 355.56 | 10.28 | 52.00 | left | -0.00962 | 68.42 | 4.58 | 754.87 |
| 199 | 252.02 | 6.12 | 51.53 | right | 0.01941 | 21.99 | 2.53 | 98.98 |
| 200 | 529.44 | 6.60 | 65.80 | right | 0.01520 | 24.57 | 2.52 | 108.79 |
| 201 | 660.17 | 4.85 | 79.85 | left | -0.01252 | 19.18 | 2.42 | 61.02 |
| 202 | 330.16 | 6.53 | 24.73 | left | -0.02022 | 19.94 | 2.44 | 61.18 |
| 203 | 787.15 | 3.90 | 44.49 | left | -0.02248 | 14.95 | 2.08 | 32.52 |
| 204 | 273.44 | 4.67 | 46.21 | left | -0.02164 | 16.91 | 2.19 | 51.18 |
| 205 | 292.98 | 4.82 | 35.35 | left | -0.02829 | 19.29 | 2.44 | 61.05 |
| 206 | 507.88 | 4.83 | 37.80 | left | -0.02645 | 22.05 | 2.61 | 76.68 |
| 207 | 353.52 | 3.70 | 57.62 | right | 0.00868 | 15.54 | 2.16 | 35.91 |
| 208 | 363.29 | 4.71 | 42.68 | left | -0.02343 | 19.05 | 2.40 | 55.69 |
| 209 | 496.11 | 6.02 | 46.41 | left | -0.02155 | 29.24 | 2.99 | 140.90 |
| 210 | 699.25 | 4.28 | 91.32 | right | 0.01095 | 18.70 | 2.34 | 58.17 |
| 211 | 373.06 | 4.98 | 33.15 | left | -0.01508 | 20.35 | 2.53 | 65.86 |
| 212 | 421.88 | 4.99 | 19.18 | left | -0.02607 | 17.41 | 2.32 | 49.49 |
| 213 | 736.40 | 5.95 | 23.83 | left | -0.02098 | 34.01 | 3.22 | 177.05 |
| 214 | 583.99 | 4.76 | 38.83 | left | -0.01288 | 15.93 | 2.23 | 40.54 |
| 215 | 734.40 | 4.69 | 69.53 | right | 0.01438 | 16.21 | 2.16 | 40.26 |
| 216 | 556.65 | 4.60 | 40.29 | left | -0.01241 | 13.88 | 2.05 | 31.11 |
| 217 | 396.49 | 4.33 | 40.17 | left | -0.01245 | 13.75 | 2.02 | 29.36 |
| 218 | 834.02 | 5.10 | 49.19 | left | -0.02033 | 24.59 | 2.65 | 107.58 |
| 219 | 517.59 | 5.60 | 41.67 | right | 0.01200 | 22.84 | 2.58 | 78.27 |
| 220 | 414.08 | 5.93 | 54.13 | right | 0.01847 | 24.28 | 2.69 | 101.40 |
| 221 | 507.82 | 4.94 | 30.20 | left | -0.01656 | 16.86 | 2.30 | 45.64 |
| 222 | 322.27 | 4.87 | 29.86 | right | 0.03349 | 16.49 | 2.26 | 45.16 |
| 223 | 652.35 | 6.27 | 140.63 | left | -0.00711 | 20.99 | 2.40 | 81.71 |
| 224 | 1167.97 | 6.08 | 139.93 | left | -0.00715 | 22.11 | 2.57 | 77.46 |
| 225 | 761.75 | 5.06 | 33.19 | left | -0.03013 | 26.11 | 2.85 | 110.12 |
| 226 | 529.32 | 4.66 | 39.63 | left | -0.02523 | 19.75 | 2.45 | 64.90 |
| 227 | 1492.56 | 4.83 | 66.04 | left | -0.00757 | 17.02 | 2.25 | 47.12 |
| 228 | 1695.42 | 4.19 | 77.54 | right | 0.01290 | 25.08 | 2.69 | 91.44 |
| 229 | 1015.64 | 4.89 | 51.50 | right | 0.01942 | 26.86 | 2.89 | 112.67 |
| 230 | 427.77 | 5.76 | 34.43 | left | -0.01452 | 22.35 | 2.57 | 75.20 |
| 231 | 320.48 | 5.88 | 31.96 | left | -0.01043 | 25.55 | 2.83 | 105.07 |
| 232 | 164.16 | 5.93 | 20.35 | left | -0.01638 | 27.59 | 2.94 | 126.14 |
| 233 | 267.72 | 5.88 | 15.11 | left | -0.03310 | 26.83 | 2.90 | 113.78 |
| 234 | 222.80 | 4.10 | 18.06 | left | -0.01846 | 15.39 | 2.21 | 37.31 |
| 235 | 162.20 | 5.10 | 19.40 | left | -0.01718 | 21.72 | 2.60 | 75.22 |
| 236 | 187.60 | 5.72 | 21.20 | left | -0.01572 | 25.11 | 2.80 | 98.27 |
| 237 | 232.58 | 4.69 | 26.38 | left | -0.01264 | 22.01 | 2.62 | 76.24 |
| 238 | 199.35 | 5.92 | 19.13 | left | -0.01307 | 33.06 | 3.22 | 174.64 |
| 239 | 412.25 | 6.62 | 25.68 | left | -0.01298 | 28.29 | 2.92 | 143.39 |
| 240 | 212.94 | 6.01 | 21.58 | left | -0.02317 | 28.43 | 2.98 | 129.57 |
| 241 | 218.80 | 5.51 | 21.36 | left | -0.02341 | 20.86 | 2.55 | 72.84 |
| 242 | 175.83 | 5.37 | 19.44 | left | -0.02572 | 20.08 | 2.50 | 63.35 |
| 243 | 254.11 | 5.41 | 21.35 | left | -0.02342 | 20.95 | 2.57 | 69.84 |
| 244 | 134.80 | 5.57 | 20.64 | left | -0.04844 | 21.17 | 2.56 | 75.71 |
| 245 | 263.76 | 3.87 | 28.67 | left | -0.01744 | 10.48 | 1.78 | 17.33 |
| 246 | 240.30 | 5.58 | 20.55 | left | -0.02433 | 20.09 | 2.50 | 67.09 |
| 247 | 463.01 | 5.53 | 20.02 | left | -0.02498 | 21.67 | 2.61 | 77.29 |
| 248 | 414.17 | 5.67 | 14.34 | left | -0.03487 | 22.02 | 2.62 | 80.64 |
| 249 | 259.84 | 5.37 | 14.90 | left | -0.06713 | 22.79 | 2.67 | 85.06 |
| 250 | 160.20 | 5.43 | 20.32 | left | -0.02461 | 19.61 | 2.47 | 61.50 |
| 251 | 226.62 | 6.02 | 9.09 | left | -0.10998 | 28.97 | 3.02 | 132.56 |
| 252 | 224.67 | 5.73 | 17.65 | left | -0.02833 | 22.05 | 2.64 | 77.26 |
| 253 | 177.78 | 5.82 | 10.84 | left | -0.02306 | 23.41 | 2.72 | 87.02 |
| 254 | 138.78 | 5.82 | 9.15 | left | -0.02731 | 24.12 | 2.70 | 88.10 |
| 255 | 160.20 | 5.81 | 10.02 | left | -0.01663 | 25.41 | 2.83 | 102.51 |
| 256 | 142.62 | 5.15 | 13.01 | left | -0.07686 | 21.93 | 2.63 | 76.51 |
| 257 | 226.63 | 5.64 | 14.05 | left | -0.01779 | 23.62 | 2.73 | 89.73 |
| 258 | 160.20 | 5.45 | 18.07 | left | -0.05534 | 19.96 | 2.50 | 66.47 |
| 259 | 548.97 | 4.59 | 36.59 | left | -0.02733 | 12.69 | 1.97 | 25.90 |
| 260 | 353.63 | 3.84 | 17.76 | left | -0.02816 | 11.29 | 1.87 | 19.83 |
| 261 | 169.98 | 5.89 | 8.90 | right | 0.11236 | 26.22 | 2.86 | 108.31 |
| 262 | 330.18 | 5.93 | 16.60 | left | -0.03013 | 26.44 | 2.88 | 111.63 |
| 263 | 105.50 | 5.59 | 11.33 | left | -0.04413 | 24.05 | 2.75 | 92.15 |
| 264 | 296.95 | 6.09 | 19.49 | left | -0.05130 | 29.53 | 3.04 | 138.36 |
| 265 | 191.45 | 5.95 | 14.78 | left | -0.01691 | 23.93 | 2.73 | 94.44 |
| 266 | 220.76 | 5.64 | 15.03 | left | -0.03328 | 21.54 | 2.59 | 74.79 |
| 267 | 347.75 | 5.48 | 21.83 | left | -0.02291 | 19.82 | 2.48 | 66.56 |
| 268 | 222.72 | 5.70 | 13.87 | left | -0.03605 | 24.14 | 2.76 | 92.35 |
| 269 | 277.41 | 5.69 | 13.65 | left | -0.03664 | 24.42 | 2.77 | 95.01 |
| 270 | 265.70 | 5.26 | 29.94 | left | -0.01670 | 19.37 | 2.46 | 60.21 |
| 271 | 478.65 | 5.92 | 17.75 | left | -0.02817 | 27.16 | 2.92 | 117.67 |
| 272 | 171.93 | 6.67 | 39.95 | left | -0.01252 | 25.16 | 2.74 | 122.74 |
| 273 | 369.63 | 6.08 | 16.64 | left | -0.03004 | 27.06 | 2.92 | 117.41 |
| 274 | 240.30 | 5.45 | 23.42 | left | -0.02135 | 20.08 | 2.49 | 65.96 |
| 275 | 490.36 | 5.35 | 22.04 | left | -0.02269 | 19.17 | 2.43 | 60.78 |
| 276 | 476.73 | 5.88 | 14.27 | left | -0.03505 | 24.20 | 2.75 | 93.71 |
| 277 | 273.51 | 5.80 | 15.20 | left | -0.03289 | 23.92 | 2.74 | 90.89 |
| 278 | 322.35 | 5.77 | 22.12 | left | -0.02260 | 22.28 | 2.64 | 83.60 |
| 279 | 343.84 | 5.69 | 21.61 | left | -0.02314 | 22.20 | 2.63 | 82.19 |
| 280 | 455.20 | 6.19 | 15.98 | left | -0.06256 | 29.79 | 3.07 | 141.82 |
| 281 | 316.49 | 6.15 | 13.58 | left | -0.03681 | 28.67 | 3.00 | 129.83 |
| 282 | 613.86 | 4.24 | 52.75 | left | -0.00948 | 8.97 | 1.60 | 14.36 |
| 283 | 359.47 | 5.82 | 15.76 | left | -0.03172 | 22.47 | 2.65 | 80.49 |
| 284 | 631.02 | 5.70 | 14.97 | left | -0.03340 | 24.60 | 2.78 | 95.75 |
| 285 | 363.38 | 6.33 | 20.42 | left | -0.01225 | 29.92 | 3.08 | 143.74 |
| 286 | 195.36 | 6.11 | 15.12 | left | -0.01653 | 25.18 | 2.78 | 106.97 |
| 287 | 304.84 | 6.05 | 13.86 | left | -0.03607 | 26.33 | 2.86 | 109.74 |
| 288 | 377.06 | 5.99 | 13.92 | left | -0.03591 | 25.66 | 2.84 | 105.25 |
| 289 | 205.14 | 5.82 | 17.35 | left | -0.01921 | 22.27 | 2.64 | 78.38 |
| 290 | 156.56 | 6.39 | 22.27 | left | -0.01497 | 27.29 | 2.89 | 127.17 |
| 291 | 176.71 | 5.74 | 14.13 | left | -0.03538 | 24.11 | 2.76 | 92.72 |
| 292 | 199.88 | 5.70 | 20.32 | left | -0.02461 | 23.04 | 2.68 | 88.02 |
| 293 | 189.60 | 5.34 | 23.39 | left | -0.02138 | 18.18 | 2.36 | 57.90 |
| 294 | 244.33 | 4.80 | 21.55 | left | -0.02320 | 14.96 | 2.14 | 36.27 |
| 295 | 204.85 | 7.52 | 46.76 | left | -0.01069 | 33.20 | 3.15 | 196.39 |
| 296 | 233.87 | 4.63 | 36.61 | left | -0.02731 | 14.85 | 2.11 | 39.14 |
| 297 | 527.70 | 5.70 | 14.01 | left | -0.03570 | 23.99 | 2.74 | 90.28 |
| 298 | 176.88 | 5.90 | 14.31 | left | -0.01747 | 24.56 | 2.76 | 94.67 |
| 299 | 179.28 | 6.05 | 16.16 | left | -0.03095 | 24.45 | 2.76 | 96.86 |
| 300 | 283.17 | 4.49 | 22.24 | left | -0.01499 | 12.60 | 1.96 | 26.03 |
| 301 | 398.46 | 5.06 | 40.63 | left | -0.02462 | 20.15 | 2.46 | 75.85 |
| 302 | 640.64 | 3.56 | 54.08 | right | 0.01849 | 12.14 | 1.93 | 24.46 |
| 303 | 425.79 | 5.17 | 49.68 | right | 0.02013 | 21.91 | 2.58 | 82.71 |
| 304 | 546.88 | 5.19 | 53.75 | left | -0.00930 | 12.89 | 1.94 | 29.47 |
| 305 | 375.01 | 5.68 | 32.16 | left | -0.01555 | 19.09 | 2.40 | 65.81 |
| 306 | 259.77 | 6.83 | 35.13 | left | -0.01423 | 28.04 | 2.90 | 146.15 |
| 307 | 283.22 | 4.34 | 15.92 | right | 0.06282 | 18.48 | 2.40 | 56.25 |
| 308 | 541.03 | 6.01 | 47.14 | left | -0.02121 | 24.53 | 2.69 | 105.08 |
| 309 | 433.60 | 6.70 | 30.02 | left | -0.03331 | 29.70 | 2.99 | 146.48 |
| 310 | 630.87 | 5.45 | 48.65 | left | -0.01028 | 14.64 | 2.05 | 44.47 |
| 311 | 433.60 | 4.67 | 41.15 | left | -0.02430 | 14.97 | 2.13 | 37.86 |
| 312 | 437.50 | 4.89 | 54.16 | left | -0.00923 | 13.33 | 1.96 | 36.60 |
| 313 | 804.72 | 5.06 | 55.40 | left | -0.01805 | 14.79 | 2.02 | 41.12 |
| 314 | 388.68 | 4.17 | 84.57 | left | -0.01182 | 9.00 | 1.55 | 18.15 |
| 315 | 355.47 | 5.12 | 52.77 | left | -0.01895 | 14.88 | 2.02 | 47.38 |
| 316 | 654.31 | 5.61 | 47.22 | left | -0.02118 | 21.09 | 2.47 | 81.53 |
| 317 | 715.01 | 5.06 | 33.67 | left | -0.01485 | 18.15 | 2.36 | 54.85 |
| 318 | 548.84 | 5.02 | 33.66 | left | -0.01485 | 17.55 | 2.32 | 49.49 |
| 319 | 1238.30 | 4.91 | 41.52 | left | -0.01204 | 17.70 | 2.35 | 51.07 |
| 320 | 916.03 | 5.67 | 50.07 | left | -0.01997 | 20.08 | 2.40 | 80.82 |
| 321 | 765.64 | 5.11 | 54.66 | left | -0.00915 | 13.16 | 1.94 | 34.61 |
| 322 | 1289.11 | 4.69 | 53.60 | left | -0.01866 | 17.78 | 2.16 | 48.64 |
| 323 | 476.58 | 4.92 | 30.19 | left | -0.01656 | 16.46 | 2.24 | 45.99 |
| 324 | 468.75 | 6.92 | 90.35 | left | -0.01107 | 29.48 | 2.97 | 155.93 |
| 325 | 648.46 | 5.04 | 32.62 | left | -0.03066 | 18.59 | 2.38 | 57.35 |
| 326 | 455.09 | 4.94 | 52.61 | left | -0.01901 | 14.93 | 2.12 | 40.22 |
| 327 | 589.85 | 8.04 | 47.57 | left | -0.02102 | 42.90 | 3.59 | 308.05 |
| 328 | 451.18 | 5.28 | 42.51 | left | -0.01176 | 15.54 | 2.14 | 43.04 |
| 329 | 396.54 | 4.65 | 62.91 | left | -0.01590 | 14.89 | 2.12 | 41.25 |
| 330 | 460.95 | 5.21 | 54.30 | left | -0.01842 | 15.55 | 2.11 | 45.42 |
| 331 | 443.38 | 7.19 | 52.18 | left | -0.01917 | 32.40 | 3.02 | 205.61 |
| 332 | 1336.00 | 4.61 | 73.85 | left | -0.01354 | 11.53 | 1.77 | 22.46 |
| 333 | 416.02 | 5.72 | 36.56 | left | -0.01368 | 23.32 | 2.69 | 88.30 |
| 334 | 447.27 | 4.82 | 37.98 | left | -0.01317 | 16.60 | 2.25 | 45.75 |
| 335 | 482.48 | 6.02 | 42.75 | left | -0.02339 | 21.10 | 2.38 | 89.03 |
| 336 | 380.87 | 5.71 | 41.73 | left | -0.01198 | 18.96 | 2.40 | 65.05 |
| 337 | 1048.85 | 5.71 | 77.65 | left | -0.01288 | 24.80 | 2.76 | 99.22 |
| 338 | 621.11 | 4.86 | 36.59 | left | -0.01367 | 17.46 | 2.32 | 50.84 |
| 339 | 568.37 | 6.69 | 58.09 | left | -0.00861 | 29.28 | 2.99 | 151.96 |
| 340 | 582.14 | 4.90 | 37.81 | left | -0.02645 | 18.17 | 2.36 | 56.74 |
| 341 | 601.57 | 4.77 | 35.09 | left | -0.01425 | 17.53 | 2.33 | 50.71 |
| 342 | 828.15 | 5.14 | 36.81 | left | -0.01358 | 19.51 | 2.46 | 63.48 |
| 343 | 1111.34 | 5.10 | 45.45 | left | -0.01100 | 16.97 | 2.27 | 49.09 |
| 344 | 683.60 | 5.06 | 37.42 | left | -0.01336 | 17.71 | 2.34 | 49.42 |
| 345 | 806.65 | 4.74 | 41.55 | left | -0.01203 | 15.35 | 2.17 | 37.65 |
| 346 | 970.73 | 4.24 | 50.51 | left | -0.00990 | 11.99 | 1.89 | 23.40 |
| 347 | 681.66 | 3.13 | 162.30 | left | -0.00616 | 6.41 | 1.38 | 7.93 |
| 348 | 539.07 | 5.48 | 40.97 | left | -0.01221 | 18.06 | 2.34 | 58.89 |
| 349 | 812.51 | 4.74 | 35.02 | left | -0.01428 | 14.54 | 2.11 | 36.37 |
| 350 | 580.09 | 4.82 | 29.43 | left | -0.01699 | 12.59 | 1.89 | 29.12 |
| 351 | 724.64 | 5.74 | 40.76 | left | -0.01227 | 16.50 | 2.08 | 49.40 |
| 352 | 761.77 | 4.32 | 19.63 | right | 0.05095 | 10.92 | 1.74 | 23.09 |
| 353 | 603.53 | 5.04 | 34.39 | left | -0.01454 | 17.42 | 2.30 | 51.84 |
| 354 | 832.04 | 5.02 | 46.79 | left | -0.02137 | 15.11 | 2.08 | 45.64 |
| 355 | 517.60 | 5.45 | 23.74 | left | -0.01404 | 20.44 | 2.48 | 75.16 |
| 356 | 509.79 | 5.72 | 40.07 | left | -0.01248 | 16.64 | 2.08 | 67.80 |
| 357 | 666.03 | 7.41 | 41.65 | left | -0.02401 | 42.68 | 3.65 | 296.76 |
| 358 | 1802.76 | 4.70 | 37.01 | left | -0.01351 | 14.74 | 2.12 | 35.53 |
| 359 | 939.47 | 4.93 | 41.19 | left | -0.02428 | 17.18 | 2.20 | 47.11 |
| 360 | 728.53 | 5.54 | 27.73 | left | -0.01202 | 19.16 | 2.42 | 63.49 |
| 361 | 369.15 | 6.63 | 41.52 | left | -0.01204 | 25.14 | 2.72 | 123.41 |
| 362 | 589.85 | 4.76 | 43.98 | left | -0.02274 | 15.20 | 2.14 | 41.18 |
| 363 | 439.46 | 5.69 | 23.42 | left | -0.01424 | 23.76 | 2.71 | 93.69 |
| 364 | 779.31 | 2.97 | 164.99 | left | -0.00606 | 6.23 | 1.36 | 7.00 |
| 365 | 1111.36 | 4.76 | 35.41 | left | -0.01412 | 16.05 | 2.22 | 42.64 |
| 366 | 748.06 | 4.81 | 36.41 | left | -0.02747 | 15.65 | 2.19 | 42.52 |
| 367 | 917.97 | 5.00 | 67.37 | left | -0.00742 | 12.87 | 1.93 | 31.25 |
| 368 | 419.93 | 3.70 | 90.76 | left | -0.01102 | 6.28 | 1.26 | 9.36 |
| 369 | 1423.87 | 4.26 | 22.33 | right | 0.04478 | 9.57 | 1.65 | 17.20 |
| 370 | 464.85 | 4.81 | 24.98 | left | -0.02002 | 15.93 | 2.21 | 42.94 |
| 371 | 1125.05 | 4.70 | 31.33 | left | -0.03191 | 16.53 | 2.24 | 44.95 |
| 372 | 709.40 | 5.82 | 40.15 | left | -0.00830 | 19.66 | 2.38 | 76.14 |
| 373 | 1390.65 | 5.07 | 35.01 | left | -0.02857 | 17.05 | 2.21 | 47.29 |
| 374 | 806.69 | 4.41 | 21.31 | right | 0.04693 | 14.20 | 2.05 | 34.36 |
| 375 | 1171.95 | 5.63 | 54.98 | left | -0.01819 | 22.84 | 2.52 | 98.22 |
| 376 | 656.26 | 4.84 | 43.57 | left | -0.02295 | 20.73 | 2.54 | 71.56 |
| 377 | 1001.98 | 4.03 | 93.55 | left | -0.01069 | 17.43 | 2.30 | 48.21 |
| 378 | 435.56 | 5.10 | 41.84 | right | 0.01195 | 17.51 | 2.28 | 52.48 |
| 379 | 875.03 | 4.15 | 25.67 | right | 0.03895 | 13.54 | 1.98 | 30.32 |
| 380 | 1048.90 | 4.64 | 57.81 | left | -0.01730 | 21.03 | 2.43 | 69.52 |
| 381 | 486.39 | 6.15 | 60.46 | left | -0.00827 | 25.80 | 2.74 | 129.57 |
| 382 | 687.55 | 5.16 | 58.54 | left | -0.01708 | 18.30 | 2.29 | 64.67 |
| 383 | 361.34 | 6.26 | 50.68 | left | -0.01973 | 21.21 | 2.33 | 102.69 |
| 384 | 939.55 | 4.93 | 52.61 | left | -0.01901 | 20.82 | 2.42 | 73.75 |
| 385 | 488.30 | 4.78 | 32.81 | left | -0.03048 | 19.78 | 2.47 | 63.74 |
| 386 | 513.69 | 5.15 | 56.93 | left | -0.00878 | 19.27 | 2.38 | 66.70 |
| 387 | 388.06 | 5.21 | 26.37 | left | -0.01264 | 20.39 | 2.48 | 68.32 |
| 388 | 566.42 | 5.98 | 47.78 | left | -0.01046 | 24.15 | 2.67 | 105.15 |
| 389 | 451.28 | 3.71 | 67.70 | left | -0.01477 | 14.74 | 2.09 | 39.70 |
| 390 | 597.72 | 5.55 | 78.13 | left | -0.01280 | 21.06 | 2.43 | 94.08 |
| 391 | 441.52 | 4.36 | 40.03 | left | -0.01249 | 14.24 | 2.10 | 32.54 |
| 392 | 285.23 | 4.77 | 37.87 | left | -0.01320 | 15.63 | 2.19 | 42.59 |
| 393 | 1011.97 | 4.85 | 58.42 | right | 0.01712 | 14.49 | 2.09 | 39.20 |
| 394 | 416.13 | 4.79 | 38.59 | left | -0.02591 | 16.90 | 2.28 | 48.29 |
| 395 | 380.96 | 4.24 | 48.76 | left | -0.01026 | 13.34 | 2.02 | 29.37 |
| 396 | 597.82 | 4.62 | 33.29 | left | -0.01502 | 15.02 | 2.15 | 35.58 |
| **397** | 476.69 | 4.64 | 38.88 | left | -0.01286 | 15.17 | 2.15 | 38.48 |
| **398** | 412.22 | 4.54 | 38.45 | left | -0.01300 | 13.43 | 2.01 | 30.68 |
| **399** | 453.24 | 4.57 | 36.67 | left | -0.02727 | 14.36 | 2.10 | 33.95 |
| **400** | 418.08 | 4.28 | 51.35 | left | -0.01947 | 11.97 | 1.89 | 25.14 |

